# Investigating Bio-Nano Interactions of PEGylated Cationic Polyamidoamine (PAMAM) Dendrimers within Synovial Joints

**DOI:** 10.1101/2025.08.09.668990

**Authors:** Simone A. Douglas-Green, Juan A. Aleman, Victor M. Damptey, Bhuvna Murthy, Brandon M. Johnston, Joon Ho Park, Alan J. Grodzinsky, Paula T. Hammond

**Affiliations:** Department of Chemical Engineering at MIT; Koch Institute of Integrative Cancer Research at MIT; Department of Biological Engineering at MIT; Department of Mechanical Engineering at MIT; Department of Electrical Engineering and Computer Science at MIT; Wallace H. Coulter Department of Biomedical Engineering at Georgia Tech and Emory University

**Keywords:** dendrimers, protein corona, bio-nano interactions, osteoarthritis

## Abstract

Delivering therapeutics directly to synovial joints to treat osteoarthritis (OA) is challenging due to the dense negatively charged cartilage matrix and rapid turnover of synovial fluid, leading to high clearance rates. Our lab has identified polyamidoamine (PAMAM) dendrimers as optimal nanocarriers to overcome delivery challenges to cartilage due to their positive charge and small size, which enables them to bind to cartilage and diffuse through tissue to deliver therapeutics to chondrocytes. Previously, we have developed and characterized dendrimers functionalized with polyethylene glycol (PEG), demonstrating improved biocompatibility and enhanced transport through cartilage matrix. To improve the design of our therapeutic system, in this next phase of work, we characterized dendrimer-protein interactions or protein coronas that form on dendrimers after being immersed in synovial fluid. We also analyzed how synovial fluid protein coronas affect biological outcomes of dendrimers in synovial joints, specifically uptake in cartilage and internalization by chondrocytes. We identified that protein coronas can reduce dendrimer uptake in cartilage and chondrocytes; however, uptake reduction is mitigated by varying PEG chain length and density. Although protein coronas can be perceived as “biological barriers” to uptake, we demonstrate that dendrimers conjugated with insulin-like growth factor 1 (IGF-1) have better engagement with IGF-1 receptors after being pre-coated with a synovial fluid protein corona. Overall, these studies offer further insight into the mechanisms of how positively charged dendrimers target and transport through cartilage, bridging knowledge gaps between *ex vivo* and *in vivo* work.

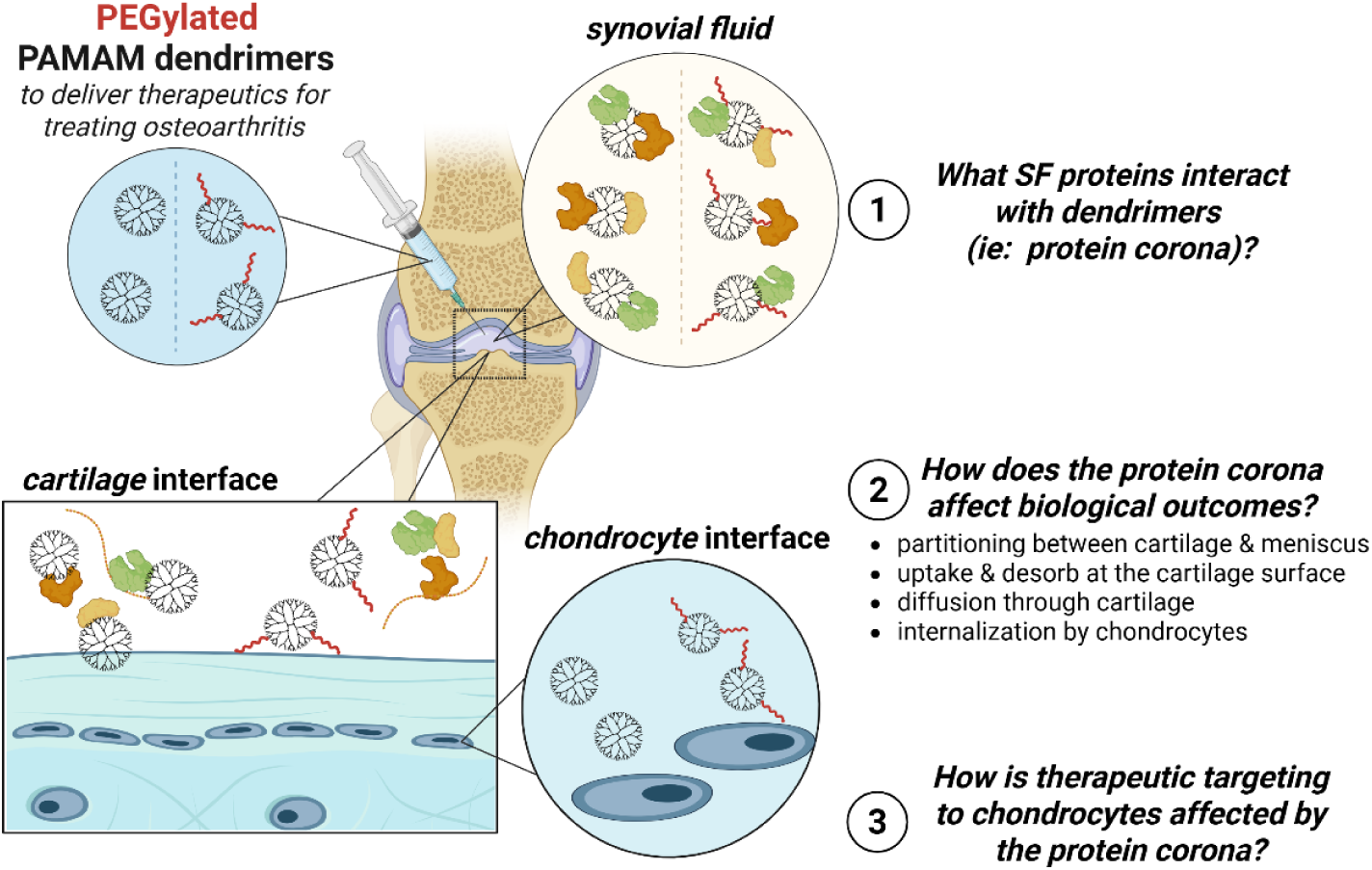

## 1. Introduction

Osteoarthritis (OA) is a debilitating joint disease characterized by the loss of articular cartilage, which leads to pain and reduced mobility^1^. OA is a leading cause of disability in the United States, affecting ∼33 million people with an overall economic burden of $136.8 billion^1–4^. There are no disease-modifying osteoarthritic drugs (DMOADs) on the market^5,6^. Current treatments, such as viscosupplementation with hyaluronic acid, provide short-term pain relief but do not reverse tissue damage^7–9^. While some DMOADs have therapeutic potential, clinical effectiveness in treating OA has not been successful, and there is a need to develop treatments that can repair and regenerate tissues^5^. Although biological drugs that stimulate cartilage proliferation exist, and injecting therapeutics directly into joints (intra-articularly) is possible^6,8^, physiological barriers make transport to the chondrocytes (“cartilage cells”) difficult. Cells are minimally dispersed throughout articular cartilage, which has a small mesh size (<15nm), strong negative charge, and high clearance rate due to rapid turnover of synovial fluid^6,8,10,11^. Thus, it is difficult for drugs to penetrate cartilage deep enough to reach cells, and molecules that reach are cleared rapidly and do not reside long enough to have any therapeutic effect. Our lab has identified polyamidoamine (PAMAM) dendrimers as optimal nanocarriers to overcome challenges with delivering therapeutics targeting cartilage to treat OA, including improved half-life from hours to months^12^ and therapeutic outcomes^12^ (Park et al submitted).

Generation 4 PAMAM dendrimers are positively charged, which facilitates transport through negatively charged matrices, and are small enough (∼4.5 nm) to diffuse through cartilage. Our dendrimer therapeutic system includes polyethylene glycol (PEG) and therapeutic proteins such as insulin-like growth factor 1 (IGF-1) or interleukin-1 receptor antagonist (IL1-RA). PEG helps reduce cytotoxicity, but more importantly, we identified that PEG improves cartilage matrix transport, where the cationic surface groups on dendrimers interact with the PEG corona during transport^13^. Our group has also demonstrated that dendrimers conjugated to IGF-1 and IL1-RA have increased joint retention time and therapeutic outcomes in rat knees compared to free IGF-1^12^ and IL1-RA. To further improve the design of our nanocarrier system, we need to understand the biological interactions at the nanoscale (bio-nano) interactions of dendrimers in synovial joints. In the scope of this paper, we define bio-nano interactions as dendrimers interacting with synovial fluid proteins (i.e., protein corona formation) and how those protein coronas affect interactions with cartilage and chondrocytes.

Analysis of dendrimer delivery efficacy must consider how synovial fluid proteins interact with dendrimers. When nanoparticles come into contact with biological fluids, proteins, lipids, or other biomolecules adsorb on the surface, forming a nanoparticle-protein complex called the protein corona^14–16^. This complex is the “biological identity” of nanoparticles *in vivo*^14,17^. In other words, after delivery, protein coronas alter the intrinsic properties of the nanoparticles. Intrinsic properties, such as size and zeta potential (surface charge), of nanoparticles are characterized and typically reported as part of delivery design. However, protein coronas can alter intrinsic properties and affect the biological outcomes, such as targeting^18,19^, biodistribution^20,21^, and pharmacokinetics^20,22^ of nanoparticles. Protein corona formation is determined by nanoparticle characteristics such as size, material, and surface chemistry^23–25^, as well as the biological environment including protein composition, pH, and temperature^26^. Protein coronas can be a barrier to efficacious nanoparticle transport and trafficking, although in some cases associated proteins can enable preferential tissue accumulation and biodistributions^27^. Through identifying proteins that adsorb onto nanoparticle surfaces, knowledge about protein corona composition can be leveraged to improve drug delivery strategies.

Dendrimers are delivered directly to joints through intra-articular injections; thus, protein corona characterization studies are needed to recapitulate the synovial joint environment, starting from the synovial fluid to cartilage to chondrocytes. We characterized dendrimer-synovial fluid protein interactions (protein coronas) and understand how protein coronas affect biological outcomes of dendrimers, including cartilage uptake and diffusion, as well as interactions with chondrocytes. Further, we assessed how protein coronas affect how dendrimers conjugated with therapeutic protein (IGF-1) interact with chondrocytes. Our central hypothesis is that after injection into the intra-articular space, synovial fluid-derived proteins adsorb to the dendrimer’s surface, forming a protein corona that affects targeting and transport to cartilage and chondrocytes. Overall, we aim to improve our understanding of the mechanism of delivery of cationic dendrimers through analyzing bio-nano interactions.

## 2. Materials and Methods

### 2.1. Materials

#### Dendrimer Conjugation

Generation 4 PAMAM dendrimers with ethylenediamine cores were purchased from Dendritech through Sigma as solutions in methanol (G4–10 wt% in methanol). Methanol was removed via dry ice rotary evaporation from the dendrimer solution. After, the dendrimer solution was washed 4 times with DI H2O using ultracentrifugal filtration (Amicon Ultra 10k, 4 mL, Fischer Scientific). Purified dendrimer was lyophilized and reconstituted in MilliQ H2O and stored at 4°C. Methoxy PEG succinimidyl carboxymethyl ester (mPEG-SCM, MW=550Da, 1000kDa), NHS-PEG_4_-DBCO, and NHS-PEG_8_-azide were purchased from Creative PEGworks. Human insulin-like growth factor 1 (IGF-1) recombinant protein was purchased from PeproTech. Atto 488 NHS ester was purchased from Sigma-Aldrich. Alexa Fluor™ 647 NHS ester (succinimidyl ester) and Alexa Fluor™ 568 NHS ester (succinimidyl ester) were purchased from ThermoFisher Scientific. Anhydrous dimethyl sulfoxide (DMSO, HPLC grade, 99.9+%) was purchased from Alfa Aesar. Sodium bicarbonate (molecular biology grade) was purchased from Sigma. Deuterium oxide (99.9 atom% % D, glass distilled) was purchased from Sigma-Aldrich. Type I Ultrapure water (MilliQ water) was generated with a Milli-Q IQ 7000 Ultrapure Lab Water System equipped with a Biopak polisher (Millipore).

#### Native PAGE

1x phosphate-buffered saline (1x PBS, 150 mM salt, without calcium or magnesium) was purchased from Lonza. 4–20% Mini-PROTEAN® TGX Stain-Free™ Gel-15 well, 4–20% Mini-PROTEAN® TGX™ Precast Protein -15 well, 2X Native Sample Buffer, 10X Tris/Glycine Buffer, and Coomassie Brilliant Blue R-250 Staining Solution were purchased from BioRad. Destain solution was prepared with 10% acetic acid (Sigma Aldrich) and 10% isopropanol (Sigma Aldrich) in MilliQ water.

#### Tissue and Cell Culture

Bovine knee joints were purchased from Research 87 (Boylston, MA). Primary chondrocytes were isolated from bovine joint cartilage; for enzymatic isolation of chondrocytes, protease from Streptomyces griseus was purchased from MilliporeSigma and collagenase was purchased from Worthington. Penicillin-streptomycin (100x), MEM non-essential amino acid solution (100x, without L-glutamine, cell culture grade), L-proline (non-animal source, cell culture grade), and ascorbic acid (20-200 mesh, cell culture grade) were purchased from Sigma. Dulbecco’s Modified Eagle Medium (DMEM) without phenol red (4.5 g/L glucose, without sodium pyruvate), fetal bovine serum (qualified), (4-(2-hydroxyethyl)-1-piperazineethanesulfonic acid) (HEPES, 1 M), and sodium pyruvate (100 mM) were purchased from Gibco.

#### Ex Vivo and In Vitro Assays

Bovine Synovial Fluid was purchased from Lampire Biological Laboratories. 1x phosphate-buffered saline (1x PBS, 154 mM NaCl, 5.6 mM Na2HPO4, 1.06 mM KH2PO4, ∼160 mM total salt, pH 7.4, without calcium or magnesium) was purchased from Lonza and 10x phosphate-buffered saline (10x PBS, 1368.9 mM NaCl, 26.83 mM KCl, 101.437 mM Na2HPO4, 17.64 mM KH2PO4, ∼1500 mM salt, pH 7.4, without calcium or magnesium, molecular biology grade) was purchased from Corning. Other PBS concentrations used (0.5X to 9X) were made by mixing 10x PBS with 1x PBS or 1x PBS with sterile water. CellTiter-Blue® cell viability assay was purchased from Promega. Human recombinant IGF-1 recombinant protein was purchased from PeproTech (acquired by ThermoScientific). Poly-L-lysine (PLK) was purchased from Sigma. Antibodies used include Hoechst (Invitrogen), Wheat Germ Agglutinin-Alexa Fluor 555 (ThermoFischer), Polyclonal Rabbit anti-Human IGF1R / IGF1 Receptor Antibody (LSBio), Goat anti-Rabbit IgG (H+L) Cross-Adsorbed Secondary Antibody, Alexa Fluor 488 (Invitrogen). VECTASHIELD Antifade Mounting Medium was purchased from Vector Laboratories. Formalin was purchased from ThermoFisher.

### 2.2. Dendrimer PEGylation

Generation 4 polyamidoamine (PAMAM) dendrimers were conjugated with PEG8 (mPEG-SCM, MW=550Da) at various densities-25%, 35%, and 45% or PEG22 (mPEG-SCM, MW=1000Da) at various densities-5%, 10%, 24%, and 34%, as previously described^13,28^. Nomenclature for formulations G4PX-Y%, where X is the PEG molecular weight (PEG8 or PEG22) and Y is the PEG density (percent PEGylation). For example, G4P8-25% is a generation 4 dendrimer conjugated with PEG8 at a density of 25%. 12.5 mM of PAMAM G4 dendrimer solution was made in a 10% v/v solution in 1M sodium bicarbonate solution and adjusted to pH 8 using hydrochloric acid. The amount of PEG needed to reach the desired % PEGylation of end groups was calculated and solubilized in anhydrous DMSO. Dendrimer solutions were diluted in 0.1M sodium bicarbonate, ensuring DMSO was no more than 10% of the final volume. The PEG and dendrimer solutions were combined, vortexed, and reacted while covered and shaking at room temperature for 2 hours. Dendrimers were purified using ultracentrifugal filtration (washed 4 times in DI water, 10k molecular weight cutoff filter) at 3500x g. Following lyophilization, dendrimers were reconstituted to 1000µM in MilliQ water and stored at 4°C.

### 2.3. NMR

%PEGylation of PAMAM dendrimers was quantified using ^1^H NMR (Bruker AVANCE, 400 MHz or 500 MHz) as previously described^13,28^. Samples were lyophilized to remove water and reconstituted in D2O. Proton chemical shifts are reported in ppm (δ): ^1^H NMR (D_2_O) δ 2.39-2.44 (PAMAM, NCH_2_C**H_2_**CONH), δ 2.61-2.68 (PAMAM, CH_2_C**H_2_**N(CH_2_)_2_), δ 2.80-2.84 (PAMAM, NC**H_2_**CH_2_CONH), δ 3.00-3.15 (PAMAM, NH_2_C**H_2_**C**H_2_**NH), δ 3.36-3.39 (PEG, OC**H_3_**), δ 3.68-3.71 (PEG, OC**H_2_**C**H_2_**), δ 4.05-4.08 (PAMAM-PEG, NHCOC**H_2_**O). The integral ratio between PEG (OC**H_2_**C**H_2_**) and PAMAM methylene (NCH_2_C**H_2_**CONH) protons was used to calculate %PEG.

### 2.4. Dendrimer Characterization

Generation 4 PAMAM dendrimers are reported to be 4.2nm, as measured by DLS, with a zeta potential of 34.6mV^29,30^. Dynamic light scattering (DLS) (Malvern ZS90 Particle Analyzer, λ=633 nm, θ=90) was used to measure the hydrodynamic size of unPEGylated and PEGylated (PEG8-25%, 35%, and 45% and PEG22-5%, 10%, 24%, and 34%) dendrimer. Data and additional information about DLS measurements can be found in **Supplemental Table 1**. Our group previously identified that the number of primary amines, termed accessible charged amines, on dendrimers correlates with how well positively charged dendrimers stick to negatively charged matrices, such as cartilage^13^. Generation 4 PAMAM dendrimers have 64 primary amines on the surface. We calculated that at 25, 35, and 45% PEGylated (PEG8) PAMAM dendrimers have approximately 36, 29, and 21 accessible charged amines (ACAs), respectively, and 5, 10, 24, and 34% PEGylated (PEG22) PAMAM dendrimers have approximately 47, 40, 21, and 8 ACAs, respectively.

### 2.5. Fluorescent Labeling of Dendrimers

Following PEGylation, dendrimers were conjugated with Atto 488 or Alexa Fluor 647, as previously described^13,28^. 1mM dendrimer was diluted in 0.1M bicarbonate solution at a 1:2 v/v ratio and vortexed. Atto 488 or Alexa Fluor 647 were prepared in anhydrous DMSO to a final concentration of 10mg/mL. Fluorophore was added to the dendrimer solution at a 1:1 molar ratio, vortexed, and reacted, covered while shaking at room temperature for 2 hours. Fluorescently labeled dendrimers were purified using ultracentrifugal filtration (washed 4 times in PBS, 10k molecular weight cutoff filter) at 3500 x g. Final solutions were stored at 4°C. To quantify fluorescent tags, the absorbance of fluorophores was measured using UV-Vis on a NanoDrop 1000 Spectrophotometer (ThermoFisher) with Atto 488 absorbance at 488nm (ε = 90,000 M-1cm-1) and AF647 absorbance at 650nm (ε = 270,000 M-1cm-1).

### 2.6. Dendrimer-IGF-1 Conjugation

PEG was conjugated to dendrimers and purified as described above in the Dendrimer PEGylation methods section. In a similar manner, 5% of dendrimer end groups were modified with NHS-PEG_4_-DBCO, with the reaction covered in foil for 2 hours, shaking at room temperature. Dendrimer-PEG-DBCO was purified using ultracentrifugal filtration (washed 4 times in DI water, 10k molecular weight cutoff filter) at 3500x g. Following lyophilization, dendrimers were reconstituted to 1000µM in MilliQ water and stored at 4°C. After dendrimer-PEG-DBCO was conjugated with Alexa Fluor 568 as described in the Fluorescent Labeling of Dendrimers method section. To quantify the dendrimer-AF568 ratio, the absorbance of fluorophores was measured using UV-Vis on a NanoDrop with AF 568 absorbance at 579nm (ε = 88,000 M-1cm-1). Separately, NHS-PEG8-azide was added to 1mg/mL of IGF-1 in 1M of sodium bicarbonate, which was pHed to 8 in a 2.5:1 azide to protein ratio with the reaction covered in foil for 30 minutes, shaking at room temperature. Alexa Fluor 647 was added to the IGF-1-azide in a 2:1 molar ratio, vortexed, and reacted covered in foil for 2 hours at room temperature. IGF-1-azide was purified using a dextran desalting size-exclusion column with an FPLC (AKTA pure 25M) with 0.1 M phosphate buffer (pH 6.8) as the eluent. The first peak (shorter retention time) was collected as the purified IGF-1 azide product, and the IGF-1 to AF657 ratio was measured using UV-Vis with AF647. Dendrimer-PEG-DBCO and IGF-1-azide were reacted in a 3:1 azide to DBCO ratio in 1x PBS for 48 hours at 4⁰C on a carousel shaker. Dendrimer-IGF-1 was purified using cationic exchange chromatography with an FPLC (AKTA pure 25M) using a salt gradient of PBS as the eluent. The IGF-1-to-dendrimer ratio was measured using UV-Vis with AF568 and AF647. Dendrimer-IGF-1 conjugates were concentrated using ultrafiltration (3k molecular weight cutoff filter, 0.5 ml) and stored at 4 °C.

### 2.7. Incubation of Fluorescently Labeled Dendrimers with Synovial Fluid and Native PAGE

Bovine synovial fluid was purchased and the stock concentration of protein was measured to be 15mg/mL; thus 1, 5, and 15mg/mL were selected as representative low, medium, and high synovial fluid concentrations used in studies (note: human synovial fluid is ∼25mg/mL^31^). Fluorescently labeled dendrimers (G4-0%, G4P8-25 and 45%, and G4-P22-5 and 34%) were incubated in bovine synovial fluid (1, 5, or 15 mg/mL) for 1 hour at 37⁰C to ensure protein adsorption. Gels were prepped, run on native PAGE gels, and imaged on a ChemiDoc Imaging System (BioRad), as previously described^28^. 1x Native Sample Buffer was prepared using 2X Sample Buffer in a 1:1 v/v ratio in 1x PBS. Samples were prepped in 1x Native Buffer in a 1:1 v/v ratio. Prepped samples were run on Native PAGE on a 4–20% Mini-PROTEAN® TGX Stain-Free™ gel or 4–20% Mini-PROTEAN® TGX™ Precast Protein gel at 120V in 1x Tris-Glycine Buffer. Stain-Free^TM^ gels were imaged on a ChemiDoc Imaging System (BioRad) with the blot 488 setting and protein gel Coomassie Stain-Free^TM^ settings to image dendrimers and protein, respectively. Densitometry was done on Stain-Free^TM^ gels using ImageJ to measure the fluorescent intensity of dendrimer and protein bands. “Regular” Precast Protein gels were imaged with the blot 647 setting to visualize dendrimers, and then stained with Coomassie Brilliant Blue R-250 Staining Solution for 1 hour and de-stained in de-staining solution overnight. Gels were then imaged with rapid auto-exposure on the Coomassie Blue stain setting to image proteins before preparation for mass spectrometry.

### 2.8. Mass Spectroscopy

*Digestion:* Protein isolation was performed following protocols previously described. The top or shielded bands were excised from the native PAGE gel. Proteins were reduced with 20mM dithiothreitol (Sigma) for 1h at 56oC and then alkylated with 60mM iodoacetamide (Sigma) for 1h at 25oC in the dark. Proteins were then digested with 12.5ng/uL modified trypsin (Promega) in 50uL 100mM ammonium bicarbonate, pH 8.9 at 25⁰C overnight. Peptides were extracted by incubating the gel pieces with 50% acetonitrile/5%formic acid, then 100mM ammonium bicarbonate, repeated twice, followed by incubating the gel pieces with 100% acetonitrile, then 100mM ammonium bicarbonate, repeated twice. Each fraction was collected, combined, and reduced to near dryness in a vacuum centrifuge. Peptides were desalted using Pierce Peptide Desalting Spin Columns (cat#89852), then vacuum centrifuged to dryness.

*LC-MS/MS*: The tryptic peptides were separated by reverse phase HPLC (Thermo Ultimate 3000) using a Thermo PepMap RSLC C18 column(2um tip, 75umx50cm PN# ES903) over a 90 minute gradient before nano electrospray using an Orbitrap Exploris 480 mass spectrometer (Thermo). Solvent A was 0.1% formic acid in water and solvent B was 0.1% formic acid in acetonitrile. The gradient conditions were 1% B (0-10 min at 300nL/min) 1% B (10-15 min, 300 nL/min to 200 nL/min) 1-3% B (15-15.5 min, 200nL/min), 3-23% B (15.5-35 min, 200nL/min), 23-35 B (35-40.8 min, 200nL/min), 35-80% B (40.8-43 min, 200 nL/min), 80% B (43-46 min, 200nL/min), 80-1% B (46-46.1 min, 200nL/min), 1% B (46.1-60 min, 200nL/min). The Thermo Orbitrap Exploris 480 mass spectrometer was operated in a data-dependent mode. The parameters for the full scan MS were: resolution of 60,000 across 375-1500 m/z and maximum IT 25 ms. The full MS scan was followed by MS/MS for the top 15 precursor ions with a NCE of 28, dynamic exclusion of 20 s and resolution of 30,000.

### 2.9. Proteomic Analysis

Raw mass spectral data files (.raw) were searched using Sequest HT in Proteome Discoverer (Thermo) against a Bovine protein database(Uniprot) and a contaminates database(made in house) with the following search parameters: 10 ppm mass tolerance for precursor ions; 0.02 Da for fragment ion mass tolerance; 2 missed cleavages of trypsin; fixed modification were carbamidomethylation of cysteine, variable modifications were methionine oxidation, methionine loss at the N-terminus of the protein, acetylation of the N-terminus of the protein and also Met-loss plus acetylation of the protein N-terminus.

The percent total of spectrum counts was calculated by dividing the total spectrum count of an individual protein by the sum total spectrum count for each experimental group. If the same protein was identified in the top and shielded bands, the total spectrum count for both bands was summed and then divided by the total spectrum count of the individual protein. UniProt and PantherDB were used to identify the biological functions of identified proteins. ExPASy-Compute pI/Mw tool was used to sort identified proteins by isoelectric point (pI).

### 2.10. Bovine Cartilage and Meniscus Explant Harvest and Culture

Cartilage was harvested from the trochlear groove of bovine knee joints within 12 hours of euthanasia. Using a biopsy punch, explants with 3 mm or 6 mm diameters were collected from cartilage. Explants were washed with 1x PBS and trimmed into cylinders with a diameter of 3 mm or 6 mm and a height of 1 mm. These cartilage explants contain intact superficial zone tissue and middle zone cartilage. 3 mm explants were used for uptake experiments, and 6 mm explants were used for diffusion experiments. Meniscus was collected and trimmed into cylinders with the same dimensions as the cartilage explants, and cultured in multi-well plates following a previously established protocol^12,13,32,33^. Before experiments were performed, explants were cultured for 24 hours to give time for tissue recovery from the harvesting procedure. Media (DMEM without phenol red, 10% FBS, 1% PS, 1% non-essential amino acids, 1% sodium pyruvate, and 1% HEPES buffer supplemented with 0.4% fresh L-proline and 0.4% ascorbic acid) was changed every other day. Explants were used in experiments no more than 2 weeks post-harvest to ensure the tissue is healthy.

### 2.11. Chondrocyte Isolation from Bovine Cartilage

Primary chondrocytes were isolated from the cartilage of bovine joints following a previously published protocol^12,13^. After exposing the trochlear groove of the joint, a scalpel was used to cut cartilage into thin slices. Cartilage slices were digested in a pronase solution, stirring for 1 hour in an incubator (37⁰C, 5% CO2) and then digested in a collagenase solution, stirring overnight in the incubator. Pronase solution was prepared the day of isolation by adding 0.1g of pronase to culture medium (High Glucose DMEM containing sodium pyruvate supplemented with 2.5% FBS, 1% Pen-Strep, 1% non-essential amino acids, 1% sodium pyruvate, and 1% HEPES buffer supplemented with 0.4% fresh L-proline and 0.4% ascorbic acid). Collagenase solution was prepared day of isolation by adding 0.0125g of collagenase to culture medium (see above for materials). Following tissue digestion, cells were extracted using a 70µm strainer followed by a 40 µm, and centrifuged for 8 min at 1900xg. The supernatant was aspirated and the pellet was resuspended in PBS and centrifuged again. Following centrifugation, PBS was aspirated and the pellet was suspended in media.

### 2.12. Co-uptake of Fluorescently Labeled Dendrimers into Cartilage and Meniscus

Each PAMAM dendrimer formulation was incubated in FBS, PBS, hyaluronic acid (HA), or synovial fluid for 1 hour at 37⁰C to form a protein corona on dendrimers. The bovine cartilage (3mm diameter) and meniscus explants (2 technical replicates and 4 biological replicates) were rinsed twice with PBS. In a 24-well plate (uptake plate), meniscus and cartilage explants (without tissue overlap) were placed in the same well and dosed with dendrimers with the pre-formed coronas, and placed in the incubator for 24 hours for dendrimer uptake. After 24 hours, 200 µL of dendrimer solution from each experimental well in the uptake plate was transferred to a 96-well plate, and fluorescence was read on a plate reader to calculate percent uptake. In a separate well plate, calibration curves were made using 1.25 µM, 0.625 µM, 0.3125 µM, 0.15625 µM, and 0.078125 µM dendrimer concentrations in 1x PBS. The calibration curves were read by fluorescence intensity with a plate reader (BioTek, excitation of 645nm, emission of 675nm, top mode, 20000µm z-position, 2x2 reads per well with a 750µm border, 25 flashes with a 0ms settle time, 0µs lag time, and 20µs integration time) and the optimal gain from the calibration curves was used for reading the experimental plates. In another 96-well plate (desorb plate), 10X PBS was added, and cartilage or meniscus explants from the uptake plate were placed in separate wells. The desorb plate was placed into the incubator for 24 hours for dendrimer desorption. After 24 hours of desorb, fluorescence was measured by a plate reader. Using the calibration curve, fluorescence was converted to moles of dendrimer desorbed from cartilage and meniscus, which were used to calculate the partitioning ratio.

### 2.13. Cartilage Uptake and Desorption (Salt Screening Assay) of Fluorescently Labeled Dendrimers

A fluorescence-based cartilage uptake and salt screening assay^13^ was used to assess electrostatic binding affinity between dendrimers and cartilage. Fluorescently tagged unPEGyalted (G4-0%) and PEGylated (G4P8-25%, G4P8-35%, G4P8-45%, G4P22-5%, G4P22-10%, G4P22-24%, and G4P22-34%) were incubated in synovial fluid (1, 5, or 15 mg/mL) at a final concentration of 0.6µM in 1x PBS solution for 1 hour at 37⁰C to form a protein corona. Cartilage explants were rinsed with PBS to remove residual proteins from the surface. Dendrimers with pre-formed protein coronas were incubated with bovine cartilage explants in a black bottom 96 well plate (uptake plate), and the last 2 wells contained conjugate only. Calibration curves were made as described in the co-uptake methods section. After a 24 hour incubation, a salt screening assay was done. Explants were placed in a second black (desorb plate) bottom 96 well plate with increasing PBS concentrations and returned to the CO2 incubator overnight. Fluorescence in the uptake plate was read on a plate reader to quantify percent uptake. The following day, the explants were removed and fluorescence in the desorb plate was read on the plate reader. The concentration of dendrimer remaining in the second plate was calculated using a calibration curve of known concentration. Percent uptake and percent desorption were calculated with equations 1 and 2:

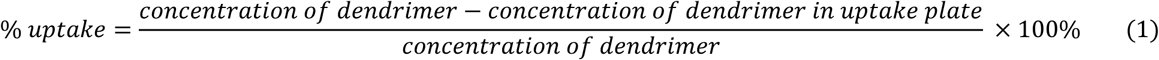

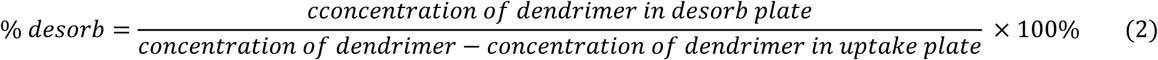

Using the results from the salt screening assay, we quantified the “critical salt concentration,” which is defined as the minimum concentration of salt (PBS) to screen electrostatic interactions between the dendrimer and cartilage that causes the dendrimer to desorb. The percent desorption was plotted against salt concentration, and the salt concentration where percent desorption sharply increases is the critical salt concentration, where higher critical salt concentrations are correlated to higher binding affinity/adsorption to cartilage.

### 2.14. Kinetic Uptake of Fluorescently Labeled Dendrimers

To perform kinetic uptake, cartilage explants were dosed with dendrimer conjugates as described above in the Cartilage Uptake and Salt Screening Assay section. Fluorescence was read at 0, 1, 2, 4, 8, 12, 24, 48, and 72 hours. Percent uptake was calculated using equation 1 and plotted against time to calculate first-order rate kinetics.

### 2.15. Diffusion of Fluorescently Labeled Dendrimers into Cartilage

Fluorescently tagged unPEGylated (G4-0%) and PEGylated (G4P8-25%, G4P8-35%, G4P8-45%, G4P22-5%, G4P22-10%, G4P22-24%, and G4P22-34%) were incubated in synovial fluid (1, 5, or 15 mg/mL) diluted in PBS solution at total volume 300uL with a concentration of 10uM for 1 hour at 37⁰C to form a protein corona. Cartilage explants (6mm diameter) were rinsed with PBS to remove residual proteins from the surface. Dendrimers with pre-formed protein coronas were incubated with bovine cartilage explants in a 48 well plate, and the last 2 wells contained conjugate only. Calibration curves were made as described in the co-uptake methods section. After 24 hours and seven days, explants were removed from the solutions and washed with 1x PBS. Explants were sectioned (with a #11 blade) longitudinally into ∼100 µm slices and washed with 1x PBS. Cross sections of explants were imaged at 10x magnification with a confocal microscope (Olympus FluoView FV1000). Z-stacks were taken using a 5 µm voxel depth with a Z-stack totaling 100 µm of cartilage depth. Images were flattened to 2D using the Z project function in the FIJI software package. If the cartilage explant did not fit into the field of view, two overlapping Z stacks were taken and stitched together using FIJI. Next, the 2D projection was flattened to 1D using the plot profile function in FIJI, and pixel intensities on the x-axis were averaged. A weighted average of pixel intensity on the y-axis was calculated to determine the average diffusion depth, which correlates with the diffusion depth of fluorescently labeled dendrimers.

### 2.16. Primary Chondrocytes Studies with Fluorescently Labeled Dendrimers and Dendrimer-IGF-1

8-chamber slides (LABTEK II, ThermoFisher) were coated with 0.01% poly-L-lysine (PLK). Primary chondrocytes were plated (5 x 10^4^ cells/ml) onto the PLK-coated chamber slide and cultured overnight. Fluorescently tagged unPEGyalted (G4-0%) and PEGylated (PEG8-25%, 35%, and 45% and PEG22-5%, 10%, 24%, and 34%) were incubated in synovial fluid (5 mg/mL) diluted in DMEM (no phenol red) with a concentration of 0.6µM for 1 hour at 37⁰C to form a protein corona. For experiments using dendrimer-IGF-1 and (free) IGF-1, a final protein concentration of 1 ng/mL was used; and for inhibitor studies, Polyclonal Rabbit anti-Human IGF1R / IGF1 Receptor Antibody (1 ng/mL) was added for 1 hour at 37⁰C before dosing with dendrimers. Cells were dosed with dendrimers for 4 hours at 37⁰C. Cells were washed in blocking buffer (1% bovine serum albumin in 1x PBS). For dendrimer only experiments, the cell membrane was stained using Wheat Germ Aggulutinin-555 (5 μg/ml). For denrimer-IGF-1 experiments, IGF-1 receptors were stained using Polyclonal Rabbit anti-Human IGF1R / IGF1 Receptor Antibody (5 µg/mL) and secondary antibody goat anti-rabbit IgG – AF488 (1:500). Nuclei were stained using Hoechst (1.25 μg/ml). Cells were fixed using 4% formalin and VECTASHIELD Antifade Mounting Medium (Vector Laboratories) was added before imaging (Olympus FV1200 confocal microscope).

## 3. Results

### 3.1. Characterizing interactions between PAMAM dendrimers and synovial fluid

First, we wanted to characterize dendrimer-synovial fluid protein interactions and identify which proteins make up the composition of the protein corona. Generation 4 dendrimers (G4) with short or long chain PEG (PEG8 or PEG22, respectively) and grafting densities; nomenclature for formulations G4PX-Y% where X is the PEG degree of polymerization (PEG8 or PEG22) and Y is the PEG graft density (percent PEGylation). The full library includes unPEGylated dendrimer (G4-0%) and PEGylated dendrimers (G4P8-25%, 35%, and 45%, and G4P22-5%, 10%, 24%, and 34%). These formulations were selected based on our lab’s previous work characterizing accessible charged amines (ACA) on dendrimer surfaces and *in vivo* studies (**Supplemental Table 1**)^12,13^. Bovine synovial fluid was purchased, and the stock concentration was measured to be 15 mg/mL; concentrations of 1, 5, and 15mg/mL were selected as representative low, medium, and high concentrations. Dendrimers were incubated in bovine synovial fluid for 1 hour at 37⁰C to form a protein corona. An incubation time of one hour was selected, as it has been previously reported that relatively stable protein coronas form during this time^34^; thus, a hard corona (more stable, irreversible protein-nanoparticle interactions) is being characterized^14,26^. Following protein corona formation, samples are run on native PAGE (**Figure 1A**) to separate dendrimer-protein interactions based on a previous protocol developed in our lab^28^. Briefly, samples separate under non-denaturing conditions to preserve dendrimer-protein interactions, migrating from top to bottom, where the top of the gel is negative and the bottom of the gel is positive. As such, more positively charged complexes are at the top of the gel, while less positively charged complexes migrate to the bottom. Samples are run on Stain-Free^TM^ gels to detect how dendrimers migrate with proteins (**Figure 1B** and Supplemental Figures 1A and 1B) or regular gels (**Supplemental Figures 2A and 2B**) to remove and digest bands for mass spectroscopy.

**Figure 1:**
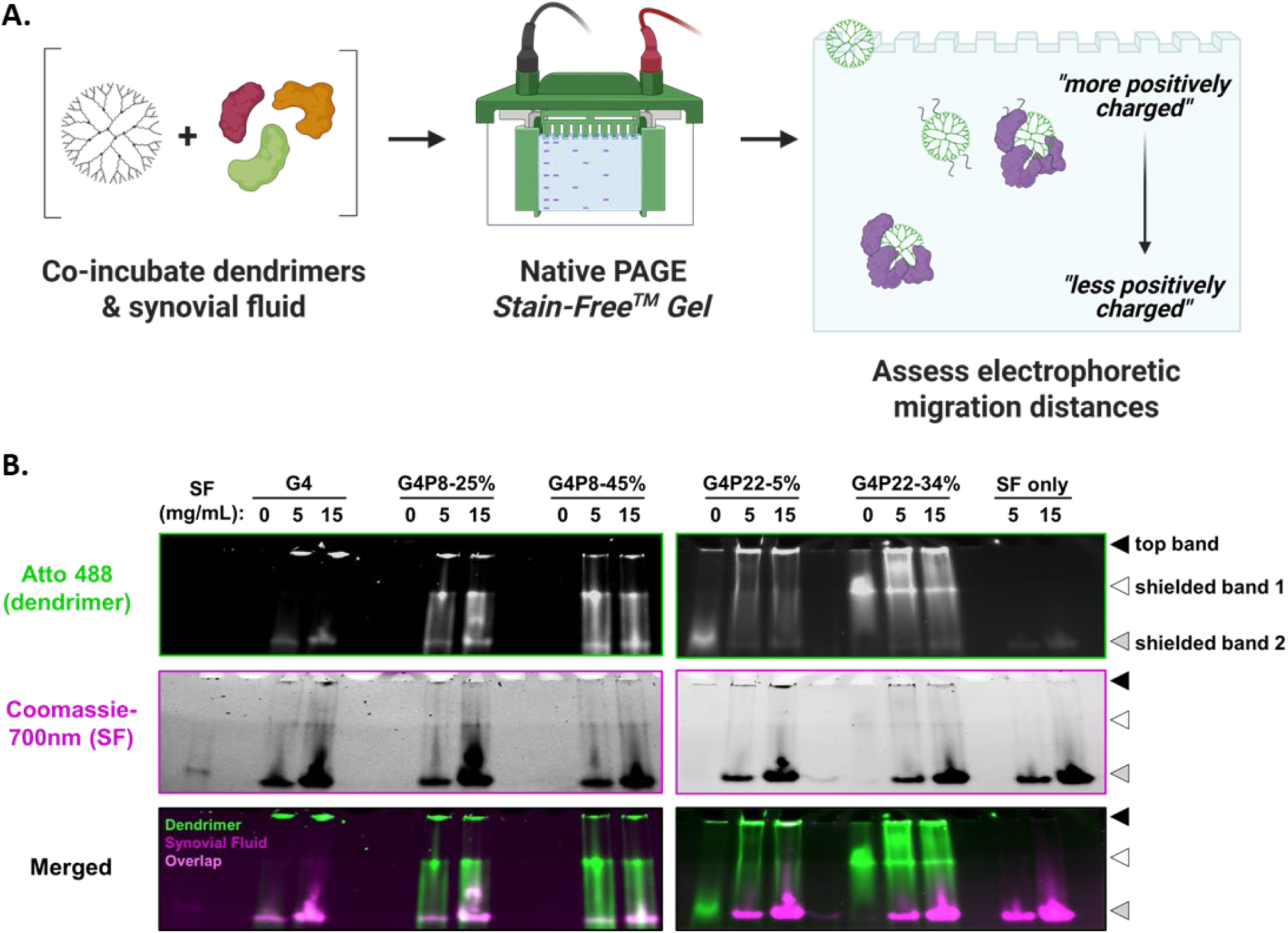
Electrophoretic migration of the dendrimers was measured to determine interactions between dendrimers and synovial fluid proteins. **(A)** UnPEGylated dendrimers (G4-0%), short chain PEGylated dendrimers (G4P8-25% and G4P8-45%), and long chain PEGylated dendrimers (G4P22-5% and G4P22-34%) tagged with Atto 488 were incubated in synovial fluid (5 and 15 mg/mL) for 1 hour at 37°C prior to separation using native PAGE on a Stain-Free™ gel. **(B)** Using Stain-Free™ gel, dendrimers are identified at 488nm and proteins are identified at Coomassie-700nm. Arrows (top and shielded bands) indicate migration distances of dendrimers.

**Figure 2:**
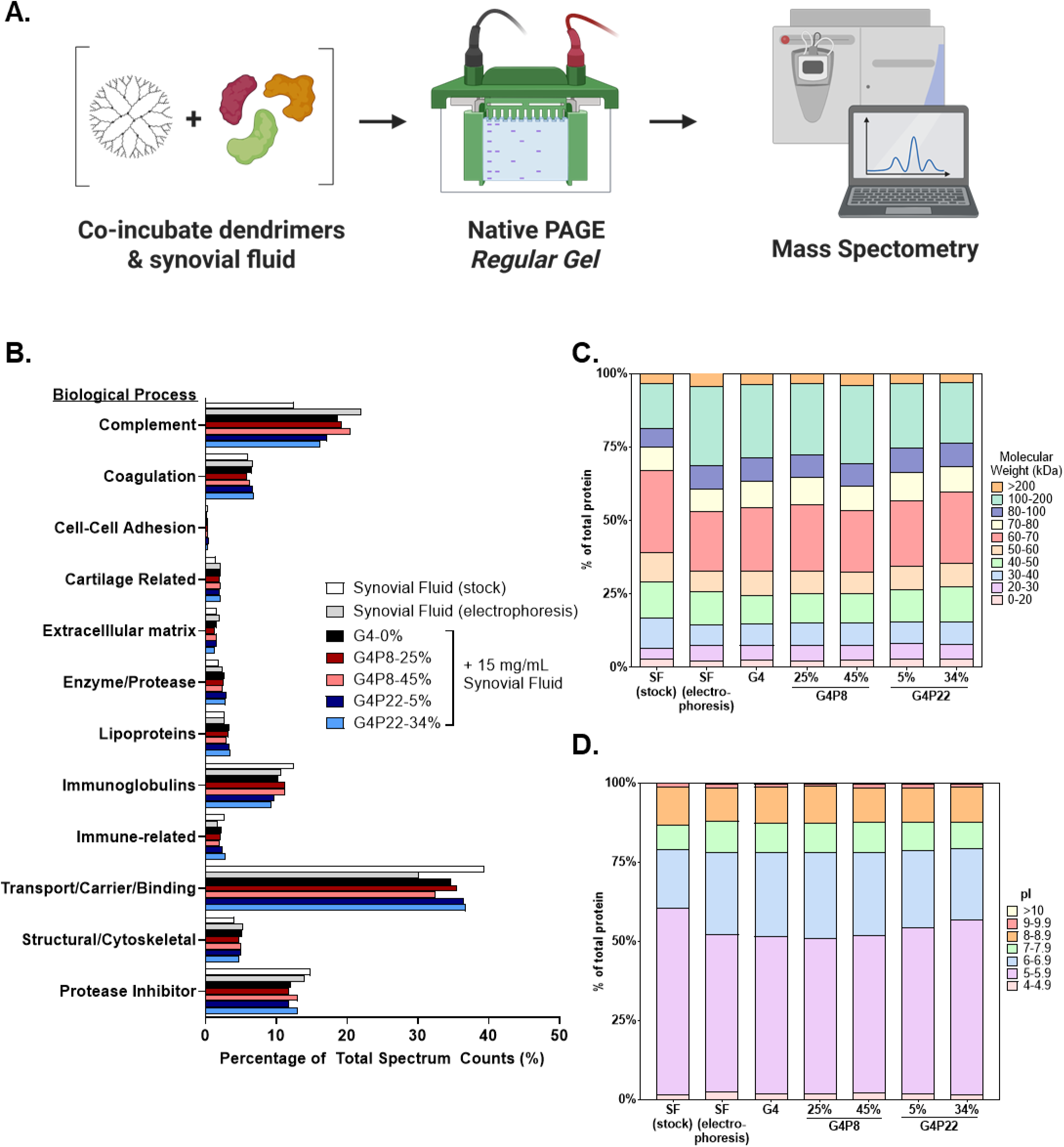
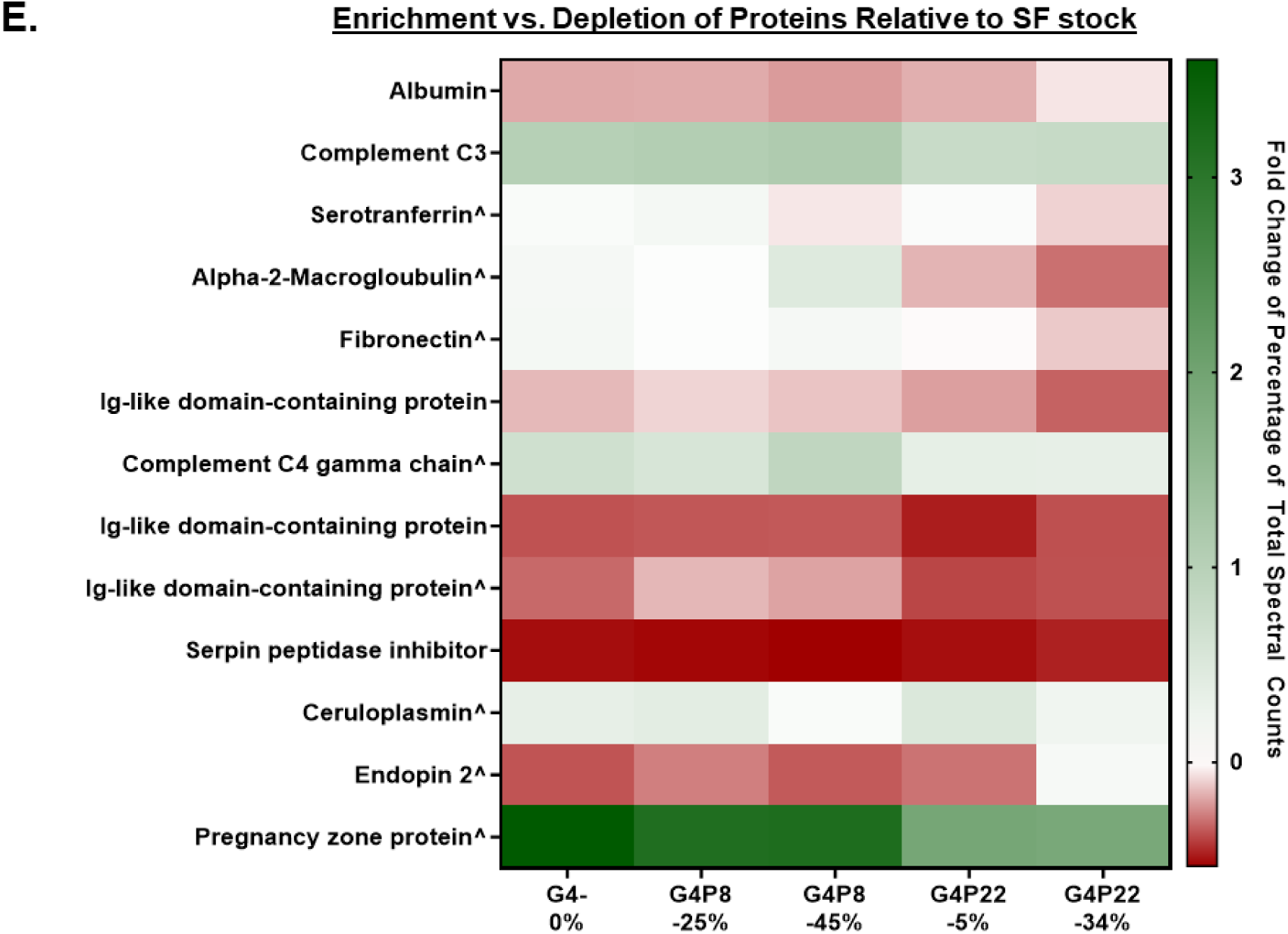
Mass spectroscopy analysis of dendrimers-synovial fluid protein interactions. **(A)** UnPEGylated dendrimers (G4-0%), short chain PEGylated dendrimers (G4P8-25% and G4P8-45%), and long chain PEGylated dendrimers (G4P22-5% and G4P22-34%) tagged with Atto 488 were incubated in synovial fluid (15 mg/mL) for 1 hour at 37°C prior to separation using native PAGE on a regular gel. The top band, shielded band 1, shielded band 2 (black, white, and grey arrows; Supplemental Figures 2A and 2B) from regular native PAGE gels for all experimental conditions were excised and run on mass spectroscopy. **(B)** All identified proteins were sorted by biological process and reported as the percentage of total spectral counts in a bar graph. **(C)** Molecular weight and **(D)** isoelectric point (pI) were analyzed based on mass spectroscopy data from Scaffold 5 and Expasy pI/MW tool, respectively. **(E)** Although dendrimers were mostly associated with albumin, a highly abundant protein (∼22% in the stock synovial fluid), compared to the stock concentration, there is a depletion of albumin. Compared to the synovial fluid stock, there is enrichment of proteins (complement C3, alpha-2 macroglobulin, and complement C4 gamma chain) or depletion of proteins (Ig-like domain-containing proteins).

Stain-Free^TM^ gels are used to simultaneously detect dendrimers with short (**Figure 1B, left**) or long (**Figure 1B, right**) chain PEG in the Atto 488 channel (green box) and proteins in the Coomassie 700nm channel (magenta box). Using Stain-Free^TM^ gels, we can better detect how dendrimers migrate with proteins and further understand dendrimer-protein interactions^28^. Top bands indicate the migration distances of unmodified PAMAM dendrimers, whereas shielded bands correspond to dendrimers that migrate through the gel due to shielding from PEG and/or proteins. Densitometry was done on the Stain-Free^TM^ gels to quantify dendrimer fluorescence in the top and shielded bands (**Supplemental Figures 1C – 1E and 1G – 1I**). For the Atto 488/dendrimer channel, data were normalized to the maximum fluorescence intensity. To account for albumin autofluorescence in shielded band 2, the densitometry from the synovial fluid only controls, matched to 5 or 15 mg/mL, were subtracted (**Supplemental Figure 1F and 1J**).

One of the advantages of using a Stain-Free^TM^ gel is being able to detect dendrimers and proteins simultaneously without using Coomassie stain. However, we noticed that for Stain-Free^TM^ gels in the Atto 488 channel (**Figure 1**), dendrimer only controls for G4, G4P8-25%, and G4P8-45% were not detected, however, this does not mean dendrimers were not present (note: in Supplemental Figures 2A and 2B dendrimer only controls were observed for all formulations). To normalize loading across all the lanes, the same concentration of dendrimer is added to each well. Fluorescence intensity is relative to fluorescence across all samples within the same gel; if a sample has significantly less fluorescence relative to a sample that has a very high fluorescence, the lower fluorescence sample will not appear as bright. When assessing the Coomassie channels in the Stain-Free^TM^ gels, there were not as many proteins observed in Stain-Free^TM^ gels compared to our previous work with FBS. This is likely due to the lower protein concentration of synovial fluid (15 mg/mL) compared to FBS (36 mg/mL). To confirm the presence of proteins, we followed up by staining the Stain-Free^TM^ gels with Coomassie Blue, which improved the detection of proteins and corroborated what was observed in the regular gels in **Supplemental Figures 2A and 2B**. Our electrophoresis-based approach is an advantageous method for separating dendrimer-protein interactions with potential use for other small nanoparticles. Although protein concentration is a limiting factor for use with Stain-Free^TM^ gels, we have identified ways to mitigate this as the method becomes more adapted.

In the Atto 488 channel, as the amount of synovial fluid increases, there is an increase in dendrimer fluorescence in the top band. This suggests more dendrimers are present due to increased protein interactions. After incubating G4P8-25% and G4P8-45% with synovial fluid, there is not as much of a difference in migration distances between them (**Figure 1B, left**). This corroborates our previous findings^28^ demonstrating that the mass-to-charge ratios of these PEGylated dendrimers are not as different. However, when comparing long PEG chains (PEG 22) with varying percent PEGylation, differences in migration distances are observed. Without synovial fluid, G4P22-5% migrates further (**Figure 1B, left-shielded band 2**) than G4P22-34% (**Figure 1B, left-shielded band 1**). We hypothesize this occurs because 5% PEG22 has a lower mass-to-charge ratio compared to 34% PEG22, which increases the migration distance at a lower percent PEGylation. When G4P22-5% is incubated in synovial fluid, we see that dendrimers do not migrate as far and mostly remain in the top band. This was unexpected, as we expect proteins to shield the dendrimer’s positive charge, resulting in migration down the gel. We hypothesize that the combined PEG and protein interactions result in a higher mass-to-charge ratio, thus reducing the migration distance of G4P22-5%. For dendrimers incubated with synovial fluid, more smearing is observed as synovial fluid concentration increases, suggesting changes in dendrimer migration due to dendrimer-protein interactions.

In the 488nm channel and Coomassie-700nm channel (**Figure 1B**), the dendrimers and proteins migrate similar distances, which means there are interactions between the two. Notably, there is an overlap of the dendrimer with shielded band 2, which was later identified as albumin with mass spectroscopy (**Figure 1B, grey arrow**). We observed that albumin auto-fluoresces in the Atto 488 channel, as evident by the synovial fluid only control and confirmed with mass spectroscopy; thus, to determine fluorescence only due to dendrimer, auto-fluorescence from albumin in the synovial fluid only sample was subtracted, demonstrating dendrimer associating with albumin (**Supplemental Figures 1F and 1J**). Based on densitometry, albumin is mostly associated with short-chain PEGylated dendrimers and results were further corroborated with mass spectroscopy. Interestingly, we see a reduction in the amount of PEG22-5% associated with albumin as indicated by the decrease in dendrimer fluorescence, suggesting PEG reduces albumin adsorption.

### 3.2. In synovial fluid, dendrimers mostly interact with albumin, in addition to complement proteins

After running native PAGE on a regular gel, top and shielded bands are isolated, digested, and each band was analyzed separately on mass spectroscopy (**Figures 2A**). Proteome software Scaffold 5 was used to analyze mass spectroscopy data; total spectrum count, accession number, and molecular weight were reported. The percentage of spectrum counts was calculated for each identified protein. A complete list of identified proteins is in **Supplemental Table 2**. In total, 187 proteins were identified. If the same protein was identified in multiple bands, total spectrum counts were combined. Top hits are reported in **Table 1**. The most common protein found to interact with dendrimers bands was albumin, however, we also observed complement C3, serotransferrin, and alpha-2-macroglobulin. When data was sorted by biological function (**Figure 2B**), factoring in that albumin is ∼22% of transport proteins identified, we observed that dendrimers mostly interact with complement proteins, which is similar to previous results for dendrimers-FBS^28^ and plasma protein^35^ interactions.

**Table 1:**
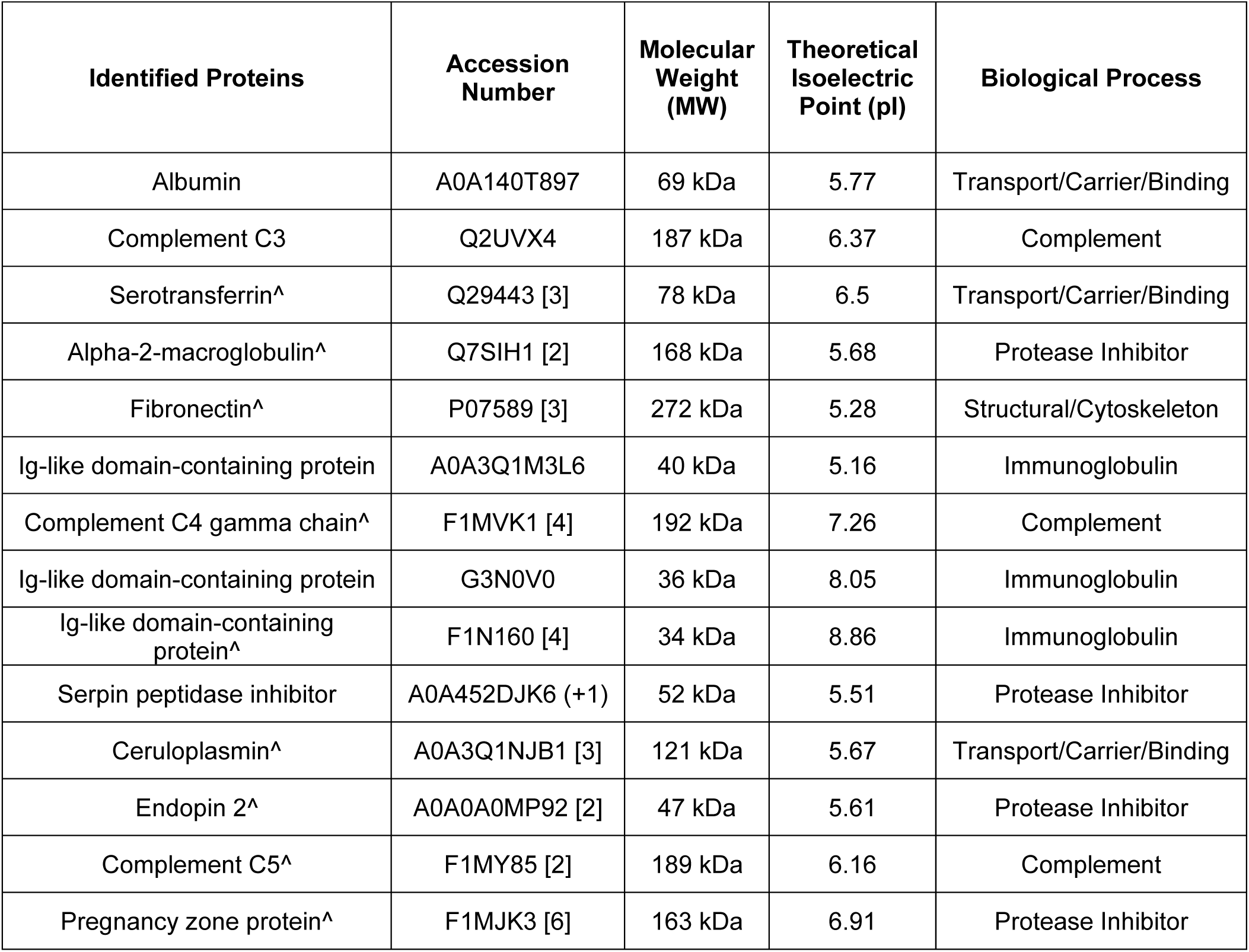
Top hits from mass spectrometry on PAMAM dendrimer-synovial fluid protein complexes in “top band” and “shielded bands” for dendrimers. The top or shielded bands (black, grey, and white arrows, respectively, Figure 2B and C) from native PAGE gels for dendrimers incubated in 15mg/mL of synovial fluid were excised, digested, and run on mass spectroscopy.

Proteins were also sorted by molecular weight (**Figure 2C**), where there was no observed difference between the distribution of the molecular weight of proteins across the various dendrimer groups. Distribution of dendrimer-associated proteins, based on molecular weight, varies across dendrimer groups compared to synovial fluid stock. In our previous work where dendrimers were incubated in FBS, we observed that dendrimers associated more with larger proteins and hypothesized that due to the dendrimers’ small size (4.5nm), they act as cargo on proteins^28^. However, these synovial fluid protein studies suggest otherwise as we do not observe this behavior; rather in synovial fluid we still observe interactions with smaller proteins. We also assessed theoretical isoelectric point (pI) (**Figure 2D**) and observed that a majority of the proteins identified have pIs between 5 and 7 (this is considering the relatively larger percent abundance of albumin). This also corroborates previous data; however, unlike previous data where compared to FBS stock we noticed an increase in proteins with pIs of 6-6.9 and decrease for pIs 7-7.9, we did not observe a major shift in the distribution of pIs between synovial fluid stock and dendrimers incubated in synovial fluid.

Finally, we compared the enrichment or depletion of dendrimer-associated proteins relative to synovial fluid stock for the top hit proteins (**Figure 2E**). We chose to compare to synovial fluid stock to identify protein corona enriched/dependent proteins that were undetected using mass spectroscopy alone. We acknowledge that comparing identified proteins to dendrimer-synovial fluid electrophoresis accounts for processing changes caused by electrophoresis, however, this analysis suggests the combined dendrimer and electrophoresis process could enhance proteome screening using mass spectroscopy. Dendrimers enrich lower abundance proteins (note: we define lower abundance proteins relative to albumin, which is <22% of stock SF); this included top hit complement proteins for all dendrimers and alpha-2-macroglobulin (A2M) for G4P8-45% which were found in previous discovery-based proteomic profiling of synovial fluid^36–38^. We also identified these same proteins in our proof-of-concept technique paper, where we characterized dendrimer interactions with FBS. Complement proteins have been previously identified in protein coronas of other types of nanoparticles, including liposomes^39^, gold^40^, superparamagnetic iron oxide^41^, and polymeric^18,42^ nanoparticles. It is also known that PEG can generate complement activation proteins^43^, and work has suggested that complement proteins, specifically C3, can bind to existing nanoparticle protein coronas^41^. A2M plays a role in immunity as a broad-spectrum protease inhibitor that acts by luring in active proteases and flagging them for elimination^44^. Higher amounts of A2M have been identified in the protein corona on transferrin-modified PEGylated polystyrene nanoparticles and were associated with increased cellular uptake^45^. Pregnancy zone protein (PZP) was enriched for all dendrimers. PZP is part of the alpha macroglobulin family, sharing 71% homology with A2M; it is an antiprotease-resistant immunosuppressant with trace amounts found in both males and females, and plasma levels increase during pregnancy^46^. Both A2M and PZP interact with the low-density lipoprotein receptor-associated protein (LRP), which can induce receptor-mediated endocytosis. In the context of leveraging nanoparticle protein coronas to improve delivery, A2M and PZP could warrant further studies, as LRP is a mediator of cell signaling in chondrocyte differentiation^47^ and A2M induces cartilage matrix synthesis and chondrocyte proliferation.

### 3.3. Proteins aid in the partitioning of PAMAM dendrimers into cartilage

Therapeutics to treat knee OA are delivered via local intra-articular injection in the synovial joint space and the target site for therapeutics is chondrocytes located within articular cartilage. For this reason, dendrimers need to localize to cartilage for efficacious treatment. The space within the synovial joint between meniscus and cartilage contains synovial fluid (SF), which is comprised of numerous proteins and biomolecules, as well as hyaluronan, which contributes to the viscosity and negative charge of SF^31,48^. After dendrimers are injected into this space, proteins and other biomolecules such as hyaluronan adsorb to the surface, forming a protein corona that can alter transport and targeting. Now that we have identified synovial fluid proteins that comprise the protein corona on dendrimers, we wanted to determine how environmental proteins and protein coronas affect dendrimer partitioning into cartilage, our target tissue, compared to the meniscus.

The first question we wanted to address was whether environmental proteins are important for dendrimers partitioning into cartilage. To model dendrimer uptake specificity to the cartilage in a joint, a co-uptake assay was developed where fluorescently tagged dendrimers in varying media conditions were added to wells containing cartilage and meniscus bovine explants for 24 hours (**Figure 3A**). For these studies, a protein corona was not formed-we wanted to see if the presence of proteins in the media affected partitioning to the target tissue, cartilage, compared to meniscus. Media conditions included synovial fluid (5mg/mL), hyaluronic acid (HA, 1 or 7.5mg/mL), FBS (10%), or PBS. Synovial fluid concentration *in vivo* is ∼25mg/mL, and we selected 5mg/mL to match the representative medium concentration in previous protein corona characterization studies. To decouple the effects of synovial fluid proteins and HA on partitioning, we included high and low concentrations of HA in our studies; 7.5 mg/mL of HA was used to match the negative charge in meniscus^32^ and a low concentration of 1 mg/mL of HA was used to equal the charge within synovial fluid ^49,50^. Additionally, 10% FBS (in DMEM) was selected as it contains a protein composition similar to synovial fluid but is not viscous. The dendrimers used were G4P4-95%, G4P4-70%, G4P13-39%, and G4P40-28%. This library was selected to capture a range of PEG chain lengths, as well as the number of accessible charged amines on the dendrimer surface, a key characteristic our group has reported that correlates with the effective charge of the dendrimer and stronger electrostatic interactions between cationic dendrimer and anionic cartilage.

**Figure 3:**
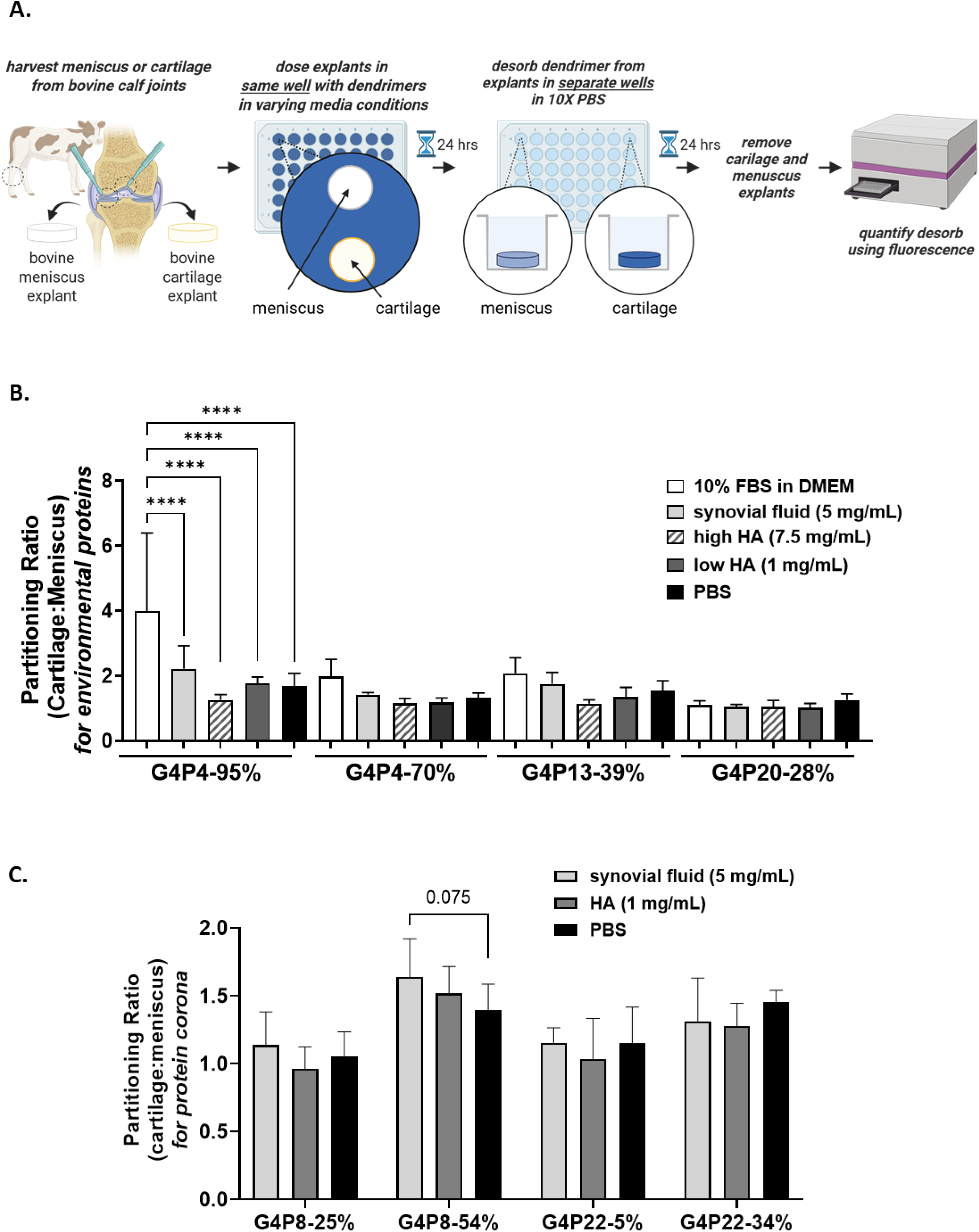
Proteins aid in the partitioning of dendrimers into cartilage. **(A)** A co-uptake assay was used to determine how environmental proteins or protein coronas affect the partitioning of dendrimers into cartilage versus meniscus. **(B)** Bovine cartilage and meniscus explants, in the same well, were dosed with dendrimer formulations with a wide range of PEG chain lengths and densities (G4P4-95%, G4P4-70%, G4P13-39%, and G4P40-28%) in varying media conditions (10% FBS in DMEM, synovial fluid at 5mg/mL, hyaluronic acid at 7.5 mg/mL or 1 mg/mL, and PBS). For the short chain PEGylated dendrimers, there is an increase in partitioning into cartilage over meniscus in media that contains proteins (10% FBS in DMEM or synovial fluid). G4P4-95% had a significant increase in partitioning ratio from PBS compared to 10% FBS in DMEM. **(C)** To evaluate how protein coronas affect dendrimer partitioning into cartilage, unPEGylated dendrimers (G4-0%), short chain PEGylated dendrimers (G4P8-25% and G4P8-45%), and long chain PEGylated dendrimers (G4P22-5% and G4P22-34%) were incubated with varying media conditions (synovial fluid at 5mg/mL, hyaluronic acid at 1 mg/mL, and PBS) to form a protein corona. Although there is a small increase in partitioning for the short PEGylated chain dendrimers in synovial fluid compared to HA and PBS, there were no statistically significant differences in partitioning of dendrimers with protein coronas across all media conditions.**|** Two-way ANOVA ****p<0.0001.

Dendrimer uptake into cartilage or meniscus was quantified using fluorescence (**Figure 3B**). Partitioning into cartilage compared to meniscus was higher for media supplemented with 10% FBS compared to 5mg/mL SF; this was specifically observed for dendrimers with shorter PEG chains compared to longer PEG chains. However, dendrimer formulations in low or high HA concentration had reduced partitioning ratios and had little to no increase in partitioning relative to PBS. With FBS and synovial fluid having overall higher partitioning coefficients compared to HA, this suggests that proteins in synovial fluid, not the anionic charge of HA, facilitate dendrimer partitioning into cartilage. Based on these observations, we hypothesize that partitioning is higher for short chain PEGs due to less PEG shielding of protein interactions, thus more amines are present on the surface of the dendrimers, which improves dendrimer cartilage uptake. We note that FBS and synovial fluid have varying compositions (**Supplemental Table 3**) and amounts of proteins, where the total number of overlapping proteins is 117 (∼55%).

It is worth noting that synovial fluid is more viscous than FBS. Healthy synovial fluid exhibits a zero shear rate viscosity from 1 to 175 Pa**·**s; this is largely due to HA, which contributes to the viscoelastic properties of synovial fluid^51,52^. Differences in viscosity could explain why partitioning is higher in FBS compared to synovial fluid. This hypothesis is supported by conditions with low molecular weight HA having a higher partitioning compared to high molecular weight HA, which are at concentrations that recapitulate the charges *in vitro*. Increased viscosity can slow down the movement of nanoparticles, which would reduce the diffusion rate^53^. Higher viscosity in a phase can block or hinder diffusion of materials, which would in turn slow down partitioning between phases. Further, diffusion limitations have been observed in phase separation of organic aerosols^54^ and chromatography with porous particles^55^. Overall, these data suggest that the partitioning of dendrimers into cartilage decreases under higher viscosity conditions.

Next, we wanted to know if protein coronas that form on dendrimers would alter partitioning between cartilage and meniscus. After observing that synovial fluid proteins promoted dendrimer partitioning to cartilage, we assessed whether pre-formed protein coronas affected partitioning between cartilage and meniscus (**Figure 3C**). To do this, dendrimers (G4P8-25%, G4P8-54%, G4P22-5%, and G4P22-34%) were incubated with synovial fluid (5 mg/mL), HA (1 mg/mL), or PBS for 1 hour at 37⁰C to form a preformed protein corona. We note the differences between the dendrimer libraries used in the previous partitioning studies. Although the first library included a range of PEG chain lengths and number of accessible charged amines on the dendrimer surface, we moved back to our more therapeutically relevant formulations. We determined there are no statistically significant differences in the partitioning of dendrimers with protein coronas across SF, HA, and PBS. However, there is a small increase in partitioning for the short PEG chain dendrimers in synovial fluid compared to HA and PBS that was not observed for the longer PEG chains. The overall increase in the partitioning ratio for G4P8-54% compared to G4P8-25% is consistent with data from **Figure 3B**, where higher PEG grafting density correlates with an increase in partitioning ratio. The same was observed for G4P22-34% compared to G4P22-5%. We note the number of accessible charged amines or ACA for short chain PEGylated dendrimers is 36 for G4P8-25% and 15 for G4P8-54%, and for long chain PEGylated dendrimers is 47 for G4P22-5% and 8 for G4P22-34%, suggesting this partitioning the presence of protein coronas is independent of ACA number and further confirms difference in partitioning is likely due to PEG grafting density. All of these observations indicate that pre-adsorbed protein coronas on PAMAM dendrimers do not alter dendrimer partitioning to cartilage over meniscus, which further demonstrates the strong affinity between dendrimers and cartilage.

### 3.4. Dendrimer uptake into cartilage decreases in synovial fluid

After interacting with synovial fluid, dendrimers reach the cartilage interface. We wanted to see how protein coronas affect dendrimer cartilage uptake. To do this (**Figure 4A**), fluorescently tagged (AF 647) dendrimers (G4, G4P8, and G4P22) were incubated in bovine synovial fluid at representative low, medium, and high concentrations (1, 5, or 15 mg/mL) for 1 hour at 37⁰C to form a protein corona. Following this, bovine cartilage explants were added to the solution of dendrimers with pre-formed protein coronas for 24 hours. We optimized our protocol initially using FBS as a medium for incubation of dendrimers with cartilage (**Supplemental Figure 4A**). We decided to use PBS instead of DMEM to dilute dendrimers (**Supplemental Figure 4B**) as we identified that unPEGylated (G4-0%) dendrimers had significantly reduced cartilage uptake in DMEM compared to PBS, suggesting biomolecules and nutrients from DMEM could form a corona on the dendrimers. We also rinsed the cartilage explants before dosing with dendrimers to remove any residual proteins on their surfaces from the culture media (**Supplemental Figure 4C**). Fluorescence was read to quantify the percent uptake of dendrimers on the cartilage surface (**Figure 4B**).

**Figure 4:**
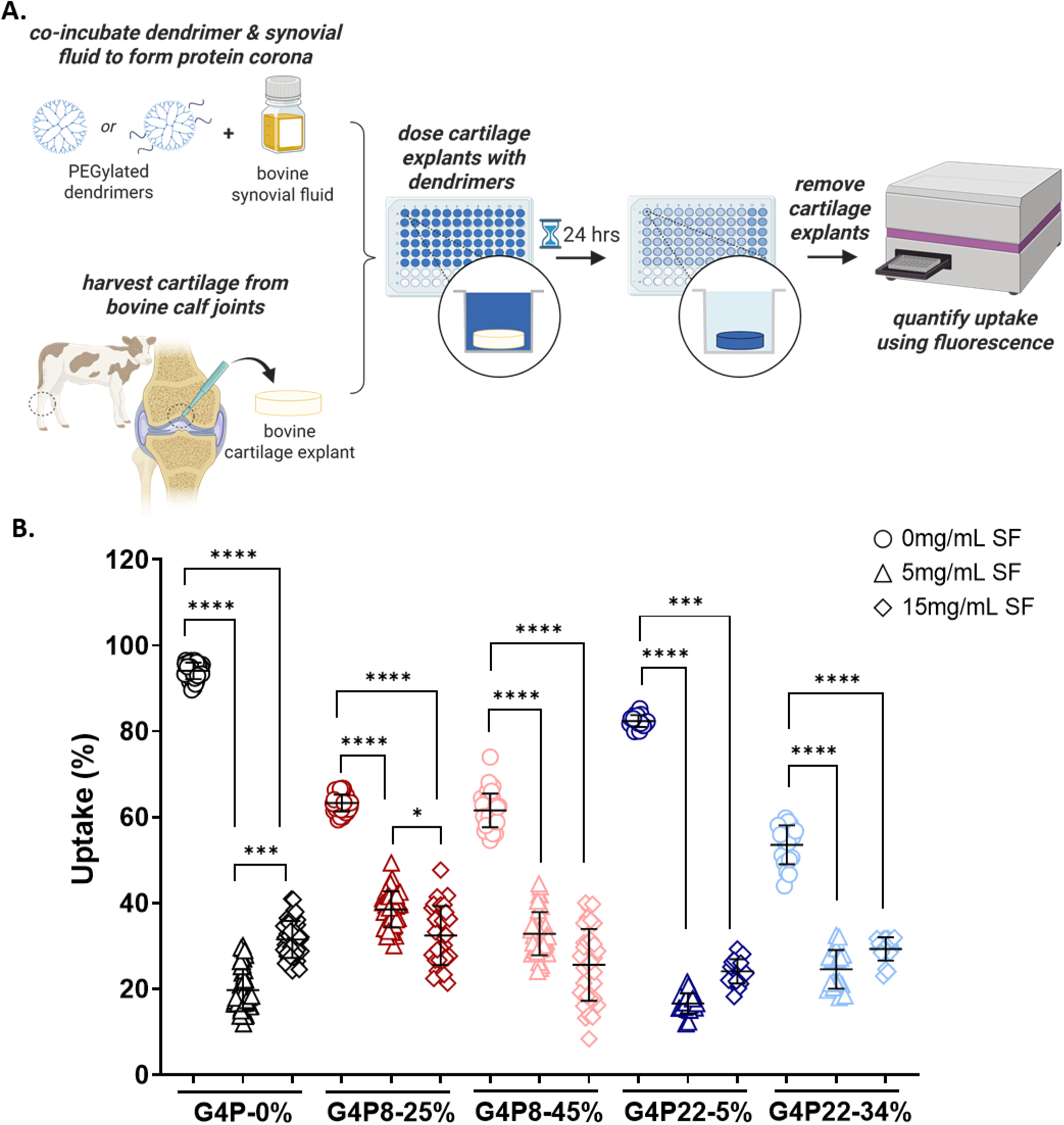
Dendrimer uptake into cartilage decreases in synovial fluid to a degree mitigated by varying PEG chain length or density. **(A)** Fluorescently tagged dendrimers with varying PEG chain length and density (G4, G4P8-25%, G4P8-45%, and G4P22-5%, and G4P22-34%) were incubated in 5 or 15 mg/mL of synovial fluid (SF) or PBS control (0 mg/mL SF) for 1 hour at 37°C to form a protein corona, then added to bovine cartilage explants for 24 hours and percent uptake was quantified using fluorescence. **(B)** As synovial fluid concentration increases, there is less uptake across all formulations relative to PBS only control. For G4P8-45%, G4P22-5%, and G4P22-34%, there is no significant difference in uptake between 5 and 15mg/mL of SF. 2-4 biological replicates with 10 technical replicates; ANOVA, Kruskal-Wallis Test for each dendrimer formulation | ****p<0.0001, ***p<0.001, **p<0.05.

Without synovial fluid pre-incubation, as PEG chain length or density (%) increases, percent uptake on cartilage decreases; the trends observed here are consistent with recently reported studies on the influence of PEG shielding and the availability of free unshielded amines on the PAMAM dendrimer surface^13^. For unPEGylated and PEGylated dendrimers, after incubation with synovial fluid, there is a significant decrease in percent uptake between 0 and 5 mg/mL synovial fluid, as well as between 0 and 15 mg/mL. While there is a significant difference in uptake between 5 and 15mg/mL of synovial fluid for G4 and G4P8-25%, there was no significant difference for G4P8-45%, G4P22-5%, and G4P22-34%. In short, incubation at the higher SF concentration did not lead to increased biomolecule adsorption and resultant PAMAM amine shielding, indicating that at the lower 5 mg/ml concentration we are already approaching a plateau in the amount of biomolecular adsorption taking place on the PAMAM dendrimer for the cases examined here. Notably, conditions without synovial fluid exhibit a lower distribution, while conditions with synovial fluid are more spread out; this degree of variance suggests a distribution of adsorbed coronas that add to biological outcomes such as like percent uptake. We also observed that dendrimers with a longer PEG chain (PEG22) were more clustered together compared to dendrimers with a shorter PEG chain (PEG8), indicating that longer PEG chains likely reduce biomolecular adsorption more uniformly, leading to more consistent – though lower - dendrimer uptake even in the presence of proteins. Uptake data for the full library of dendrimers is in **Supplemental Figure 5**.

### 3.5. PEG shields dendrimers from protein adsorption while sustaining electrostatic interactions with cartilage

Dendrimers have a net positive charge, which aids in the binding of dendrimers to negatively charged cartilage matrix. However, past work^13^ from our lab has determined that simply quantifying the total number of charged groups on the dendrimer is not a reliable metric to evaluate nanocarrier efficacy for binding to negatively charged matrices. Instead, quantifying the number of unshielded primary amines that present charge on the dendrimer surface is a better physiochemical property to characterize dendrimer “stickiness” on cartilage. We have termed this value the number of accessible charged amines (ACA), which changes based on the density and length of PEG conjugated to the dendrimer surface due to both covalent bonds and non-covalent interactions of amine groups with PEG. We wanted to assess how pre-formed protein coronas affect the binding affinity of dendrimers to cartilage. To do this, we used a salt screening assay (**Figure 5A**). Following uptake, explants were placed in a solution with increasing salt (PBS) concentrations. The “critical salt concentration” is defined as the minimum concentration of salt to screen the electrostatic interaction between the dendrimer and cartilage sufficiently to cause the dendrimer to desorb. Through this assay, we determined how well dendrimers with pre-formed protein coronas stick to cartilage, which correlates to more ACA on the dendrimer surface (**Figure 5B**).

**Figure 5:**
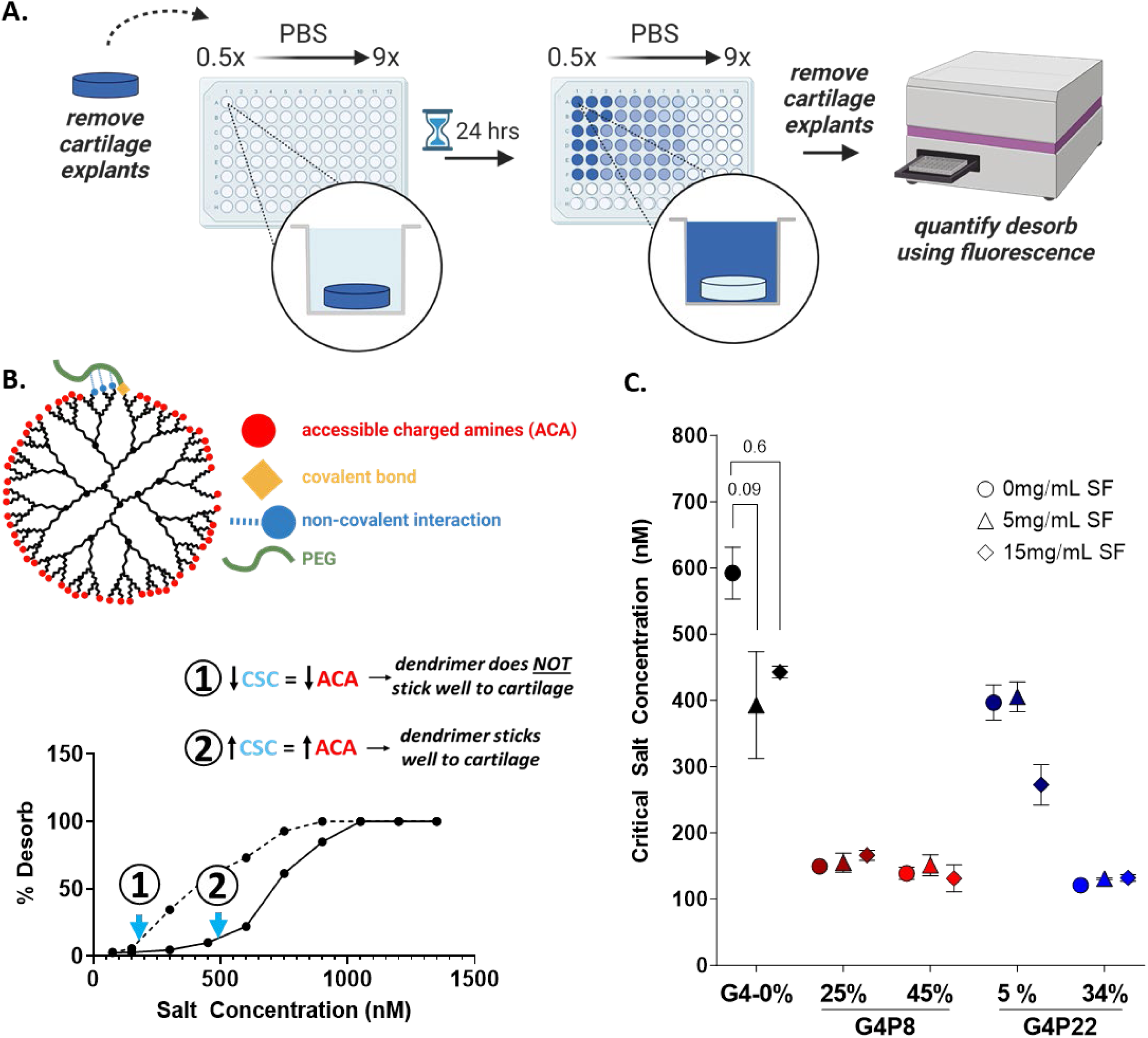
PEG shields dendrimers from protein adsorption while sustaining electrostatic interactions with cartilage. **(A)** After uptake, a salt screening assay was done by placing cartilage explants in increasing PBS concentrations (0.5X to 9X) for 24 hours to measure dendrimer desorption and calculate critical salt concentration (CSC). **(B)** CSC is defined as the lowest salt concentration to screen electrostatic interactions between dendrimer and cartilage. This value correlates to the number of accessible charged amines (ACA) or amines available on the dendrimer surface accessible to the physiological environments-a key property governing the degree of dendrimer interactions with cartilage. **(C)** For unPEGylated dendrimers, as synovial fluid (SF) is added, there is a decrease in the critical salt concentration, suggesting protein adsorption to dendrimer is blocking ACAs on the surface. However, for PEGylated dendrimers, despite synovial fluid proteins and biomolecules being present, there is little to no change in critical salt concentration. | 2 biological replicates with 2 technical replicates; ANOVA, Kruskal-Wallis Test for each dendrimer formulation.

Without synovial fluid, as PEG chain length or density (%) increases, the critical salt concentration decreases relative to unPEGylated dendrimers. When unPEGylated dendrimers are incubated in synovial fluid, the critical salt concentration decreases, suggesting adsorbed biomolecules are shielding the primary amines on the dendrimer’s surface (**Figure 5C**). For G4P8-25, G4P8-45%, and G4P22-34% dendrimers incubated in synovial fluid, we see there is little to no change in critical salt concentration. It appears that for these cases, the presence of PEG repels the adsorption of biomolecules that might otherwise further shield the PAMAM amine groups, leaving the ACA effectively unchanged. The critical salt concentration for G4P22-5% is the highest of all the PEGylated dendrimers, there is little change as 5mg/mL synovial fluid is added, however, a decrease is observed with 15 mg/mL. For this dendrimer, which has a much lower PEG density than the other samples, and thus a higher initial ACA, the higher 15 mg/ml SF concentration yielded biomolecular adsorption sufficient to increase shielding of amines and lower affinity to cartilage. Results demonstrate how varying PEG chain length and density affect protein adsorption.

Using the critical salt concentration from **Figure 5A** and **Supplemental Figure 5,** combined with “critical salt concentration vs. ACA” graphs from our lab’s previous studies^13^, we estimate the number of ACAs for unPEGylated dendrimers with synovial fluid protein coronas to be approximately 55, 40, and 60 in 1, 5, and 15mg/mL synovial fluid, respectively. For PEGylated dendrimers, despite the presence of proteins, there is little change in salt concentrations, likely due to PEG repelling proteins. Through accessing ACA, for the first time, we demonstrate that proteins block the primary amines on the dendrimer surface, resulting in dendrimer-protein interactions and protein corona formation. Further, through these studies, we have also proven that PEG8 and 25% density and PEG22 at 5% density can efficiently repel proteins. These results can help inform dendrimer design considerations, specifically developing dendrimers with higher ACAs to offset potential protein adsorption.

### 3.6. PEGylated dendrimers have improved kinetics of uptake in synovial fluid compared to unPEGylated dendrimers

In separate work, we examined the role of PEG chain length and density on the kinetics of PAMAM uptake and diffusion through cartilage under PBS conditions (Johnston et al. in preparation). There is rapid turnover in synovial fluid, which could affect the pharmacokinetics of dendrimers and the degree of dendrimer retention in the joint. Here we investigate how the protein coronas formed in SF affect kinetic uptake of dendrimers over time. For kinetic studies, unPEGylated and PEGylated dendrimers were incubated in bovine synovial fluid (5 mg/mL) for 1 hour at 37⁰C to form a protein corona, then added to cartilage explants. Fluorescence was read at 0, 1, 2, 4, 8, 12, 24, 48, and 72 hours and plotted to calculate the first-order rate constant (K). Overall, the first-order rate constant varies as the PEG chain length and density (%) changes; G4P22-5% had the highest rate constant yet only had 10% uptake (**Figure 6**). Adding synovial fluid decreases first-order rate constants for unPEGylated and PEGylated dendrimers. G4P22-34% had a 2x decrease in first-order rate constant, whereas G4P8-25% and G4P8-45% had a 3x decrease and the unPEGylated dendrimer had a 5x decrease. Although G4P22-5% had the highest rate constant without synovial fluid, there was a 6x decrease with synovial fluid. Overall, with synovial fluid proteins, PEGylated dendrimers have higher first-order rate constants compared to unPEGylated dendrimers. This suggests that while proteins slow down dendrimer uptake, PEG can help mitigate this effect, potentially by repelling proteins from the dendrimer surface while enabling more active diffusion via shielding effects as reported previously (Johnston et al. in preparation).

**Figure 6:**
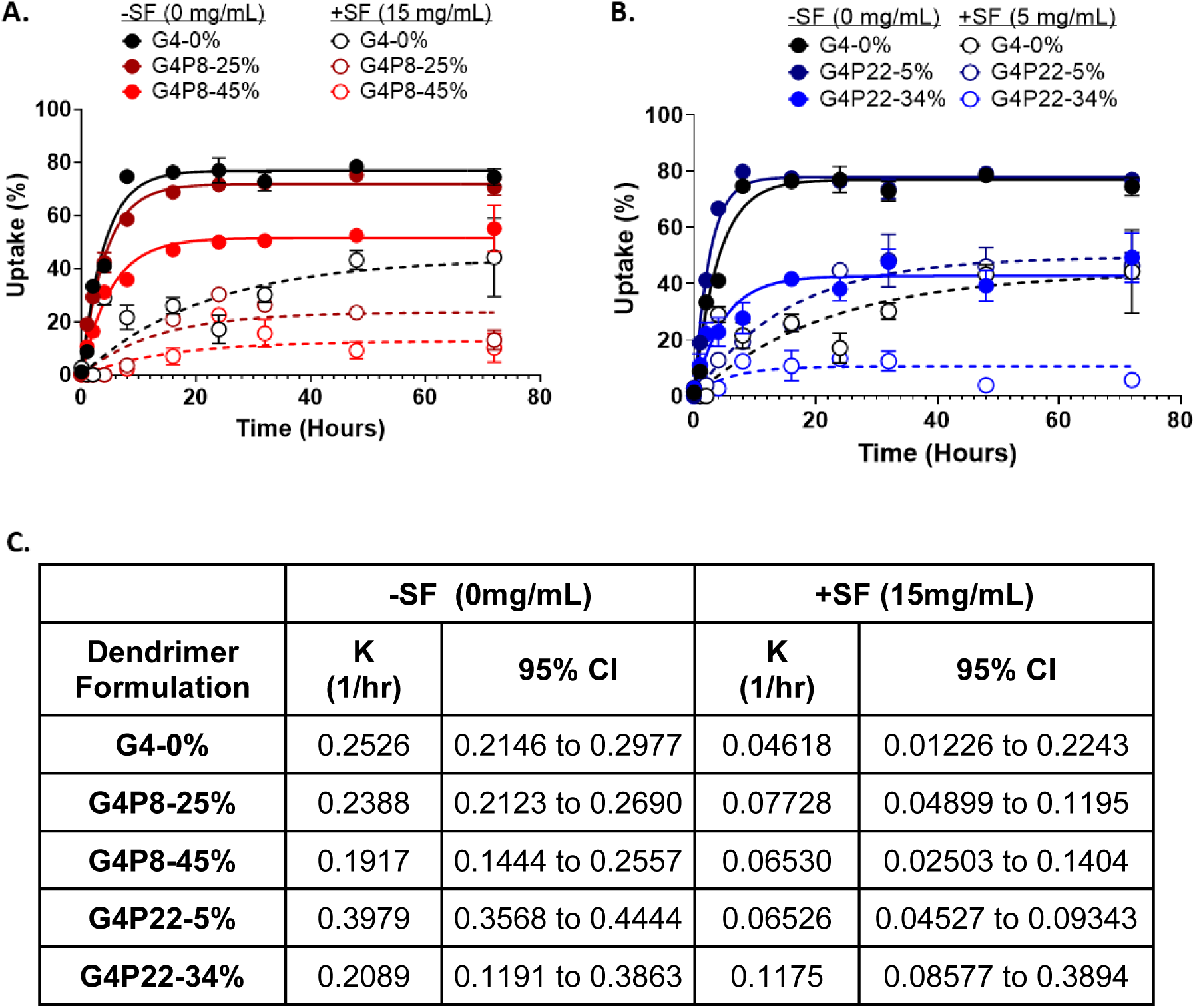
Protein-dendrimer interactions reduce the first-order rate constant of dendrimer uptake, however, this is mitigated by varying PEG density or chain length. **(A)** Short (G4P8-25% and 45%) and **(B)** long (G4P22-5% and 34%) chain PEGylated dendrimers were incubated in synovial fluid (SF) for 1 hour, then added to cartilage explants; fluorescence was read at various time points over 72 hours and plotted. **(C)** First-order rate constant varies based on varying PEG chain length and density.

### 3.7. Protein corona does not change dendrimer diffusion through cartilage

After dendrimers adsorb to the cartilage surface, they need to diffuse through the cartilage matrix to reach chondrocytes and deliver therapeutic proteins. To assess how protein coronas affect dendrimer diffusion, we used the same uptake assay as previously described using cartilage explants that were 6mm in diameter. At 24 hours and 7 days, cartilage explants were removed, sectioned, and imaged on confocal microscopy (**Figure 7A**), and the average diffusion depth was quantified using fluorescence (**Figures 7B and 7C**). Representative images of diffusion of unPEGylated and PEGylated dendrimers with or without pre-formed synovial fluid protein coronas are in **Supplemental Figure 6**. For unPEGylated dendrimers, after 24 hours, there was little difference in diffusion distance with or without synovial fluid; however, after 7 days, we see there is a slight increase in diffusion distance with synovial fluid. For short chain PEGylated dendrimers (PEG8) with protein coronas, there was a slight increase in diffusion distance after 24 hours; however, for long chain PEGylated dendrimers (PEG22), there was a decrease after 24 hours. After 7 days for all PEGylated dendrimers, there was no significant increase or decrease with or without synovial fluid. Although there was a slight increase in diffusion depth for unPEGylated dendrimers after 7 days, the difference was not significant. Collectively, this leads us to conclude that with synovial fluid, there is no difference in the diffusion of dendrimers in cartilage over longer periods of time.

**Figure 7:**
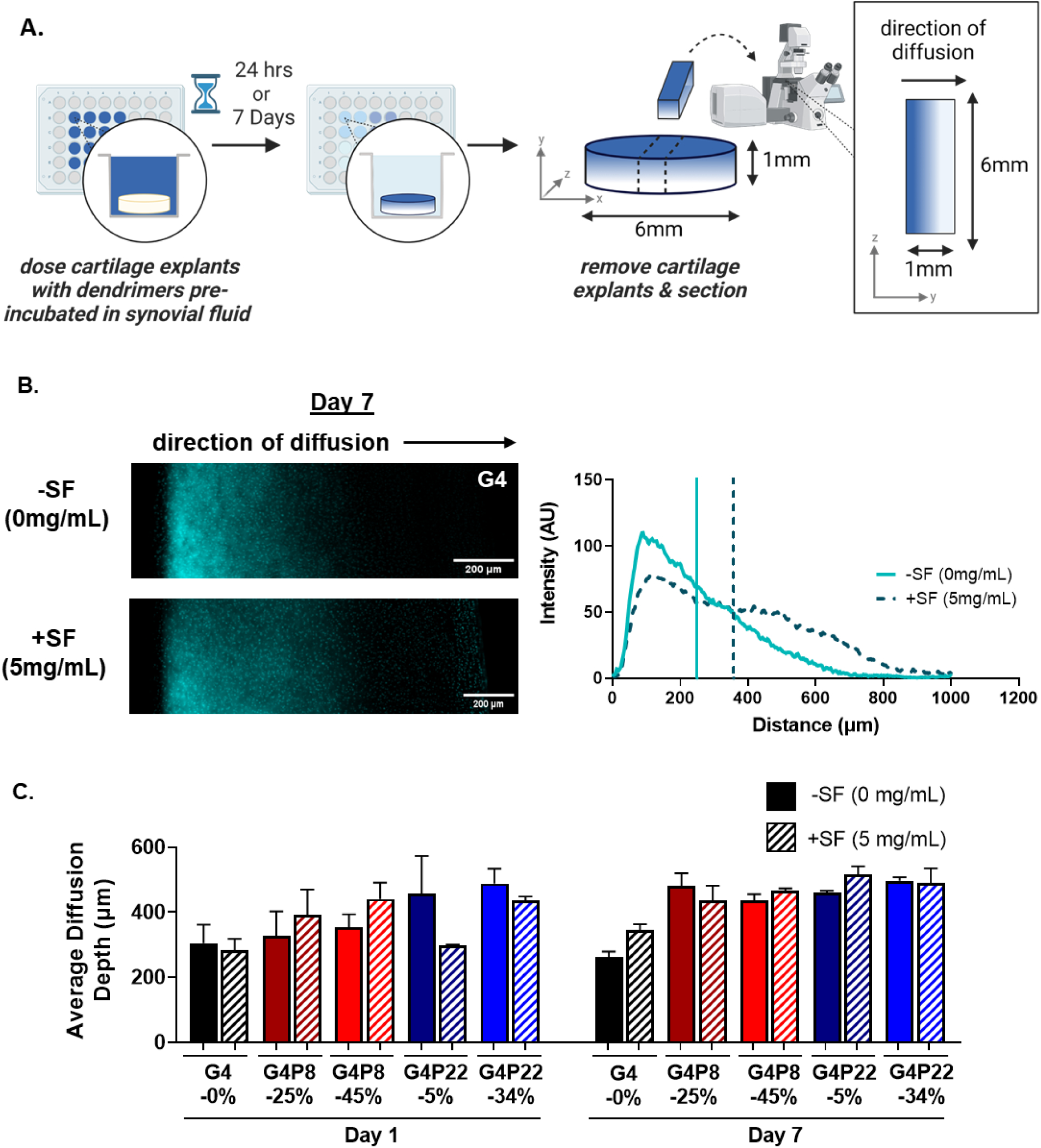
Protein corona does not hinder dendrimer diffusion in cartilage. **(A)** Short (G4P8-25% and 45%) and long (G4P22-5% and 34%) chain PEGylated dendrimers were incubated in synovial fluid (SF) for 1 hour, then added to cartilage explants. After 24 hours or 7 days, explants were removed, sectioned, and **(B)** imaged on microscopy to analyze dendrimer diffusion through cartilage. **(C)** After 24 hours and 7 days, there was no significant increase or decrease in average diffusion depth for dendrimer formulations with or without synovial fluid. Representative images of diffusion of unPEGylated and PEGylated dendrimers (with or without synovial fluid protein coronas) through cartilage are in Supplemental Figure 6.

Previous work shows that weak and reversible binding between dendrimers and cartilage is needed to enhance diffusion. Specifically, PEG with longer chains improves the diffusion distance of dendrimers through cartilage due to the longer chains disrupting the dendrimer-cartilage interactions more efficiently, which allows for more reversibility for binding, where dendrimers repeatedly bind and unbind to cartilage, moving deeper into cartilage^13^. However, in our protein corona studies with G4P8 and G4P22 dendrimers, there was little to no evidence of protein coronas adding to the shielding effect of weak and reversible binding between dendrimers and cartilage. For G4-0% at day 7 there was an increase with in average diffusion distance with synovial fluid, however this was not significant. We wanted to further test the hypothesis that proteins can shield charges on primary amines and facilitate weak and reversible^32,33,56^ interactions to aid in dendrimer diffusion. To test this hypothesis, we used unPEGylated generation 6 (G6) PAMAM dendrimers, which contain 256 primary amines; highly cationic, which makes them more difficult to diffuse through negatively charged cartilage (**Supplemental Figure 7**). Cartilage uptake experiments were set up as previously described for the G4 dendrimer library. After 24 hours, G6 dendrimers with and without synovial fluid protein had little diffusion in cartilage. However, after 7 days, G6 dendrimer with protein coronas had a higher (p=0.09) average diffusion depth compared to no protein corona. This demonstrates how proteins shield charged amines on the surface of dendrimers to facilitate weak, reversible interactions within cartilage to aid dendrimer diffusion.

### 3.8. Dendrimer internalization by chondrocytes decreases in synovial fluid

Dendrimers can be conjugated with therapeutic proteins, such as insulin-like growth factor-1 (IGF-1), that target receptors on chondrocytes, promoting chondrocyte proliferation and enhancing matrix production to help with cartilage repair^57,58^. Based on observations from cartilage uptake kinetic studies, dendrimer-protein interactions may be sustained within cartilage matrices; thus, these protein coronas could affect dendrimers targeting chondrocytes. To study how such dendrimer-protein conjugates might interact with cartilage cells if the bound proteins remain present, dendrimers were incubated in synovial fluid (5 mg/mL) to form a protein corona, as previously described, and bovine primary chondrocytes were dosed for 4 hours, then prepared for confocal microscopy (**Figure 8**). Before *in vitro* studies, cytotoxicity experiments were done to confirm that bovine synovial fluid protein coronas were not toxic to cells (**Supplemental Figure 8**). Without pre-formed synovial fluid protein coronas, unPEGylated and PEGylated dendrimers are internalized by chondrocytes, likely due to the positive charge facilitating uptake through the membrane^59–61^. For unPEGylated dendrimers with synovial fluid protein coronas, there is a decrease in dendrimer internalization, with some dendrimers aggregating outside the cell. G4P8-25%, G4P8-45%, and G4P22-5% dendrimers are still internalized even with synovial fluid protein coronas, and no dendrimer aggregates were identified outside the cells. This was in contrast to G4P22-5%, where there was more dispersion of dendrimers outside the cell, which could be due to dendrimer-protein interactions reducing dendrimer internalization. We also observed the dendrimers colocalized with membrane stain and hypothesize this could be due to synovial fluid membrane proteins associated with the dendrimer. Collectively, this suggests that in the presence of synovial fluid proteins, PEG shields dendrimers from being opsonized, so dendrimers are internalized. However, G4P22-34% with pre-formed synovial fluid protein corona were not internalized by chondrocytes. We hypothesize these dendrimers were not internalized due to the combined long PEG chain length and high density, as well as the protein corona shielding the positive charge of the dendrimer. Thus, the dendrimer could not go past the cell membrane to be internalized.

**Figure 8:**
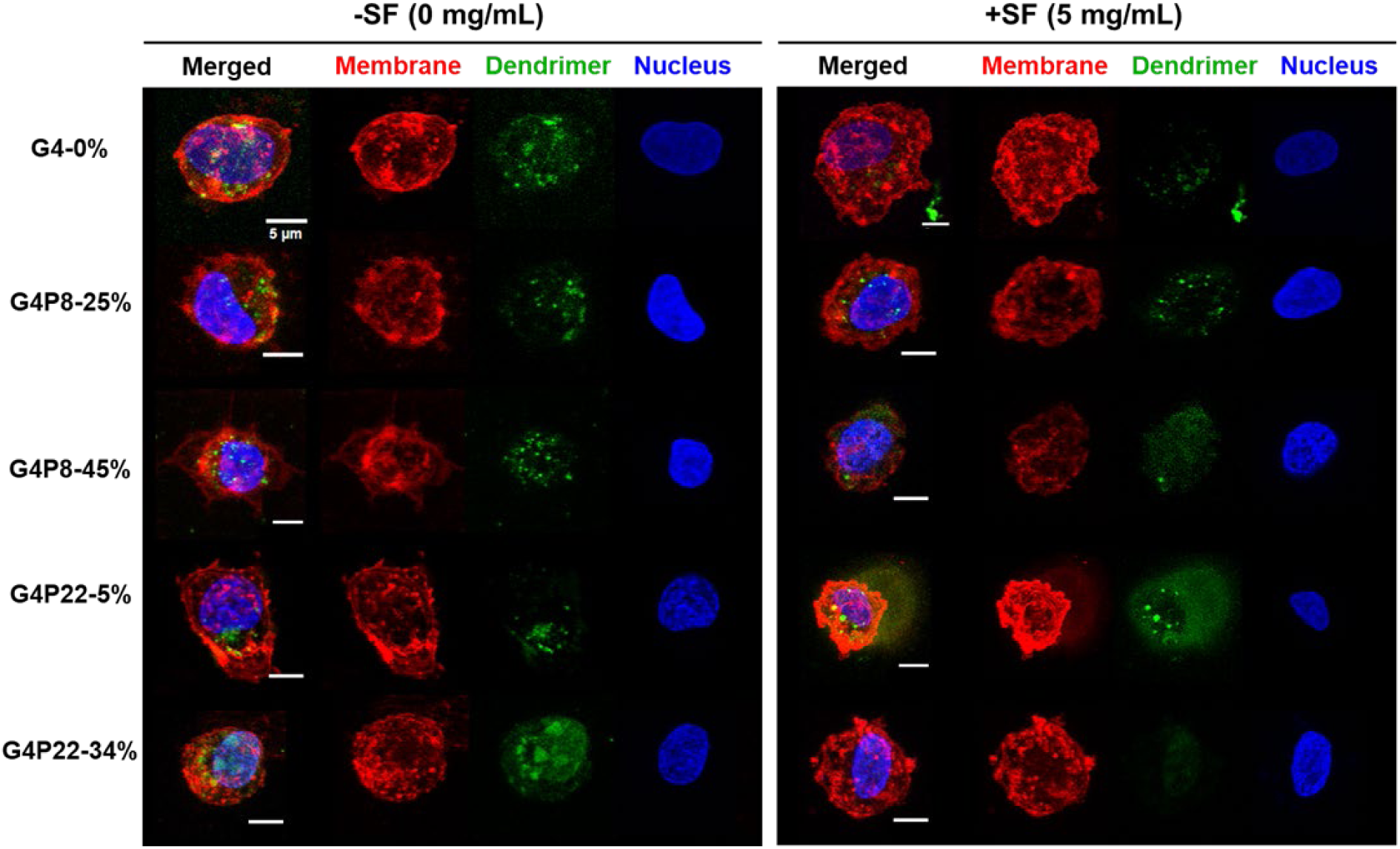
Chondrocytes internalize dendrimers within 4 hours, however, this is reduced with pre-formed synovial fluid protein coronas. Dendrimers (green) were incubated in 5mg/mL of synovial fluid for 1 hour at 37°C, chondrocytes were dosed for 4 hours, and prepped for confocal microscopy. Nuclei were visualized with Hoechst (blue) and membrane was visualized with wheat germ agglutinin (red). Without synovial fluid, unPEGylated and PEGylated dendrimers are internalized by chondrocytes. In synovial fluid, G4, G4P8-25%, G4P845%, and G4P22-34% are still taken up, although it is reduced. However, G4P22-34% with synovial fluid proteins are not internalized. | Scale bar = 5µm.

### 3.9 Pre-formed protein coronas aid dendrimer-IGF-1 conjugates with IGF-1 receptor engagement

Finally, we wanted to see how protein coronas affect the targeting of dendrimers conjugated with therapeutic protein IGF-1. We selected G4P8-25%, as this has previously been shown to have a high amount of accessible charged amines^13^ and performed well in our protein corona studies for cartilage and chondrocyte uptake. G4P8-25% dendrimers were conjugated with IGF-1 (dendrimer-IGF-1) in a 1:1 ratio of dendrimer to protein. Dendrimers were fluorescently labeled with AF 568, and IGF-1 was labeled with AF 647. Conjugates were incubated with synovial fluid (5mg/mL) to form a protein corona, as described above. Bovine primary chondrocytes were dosed for 4 hours and prepared for confocal microscopy. Full confocal images are in **Supplemental Figure 9**. We observed that DAPI overlaps with IGF-1 receptor without inhibitor (**Figures 9A and 9C**); this is likely because IGF-1 receptor can translocate^62^ to the nucleus, where it can regulate gene expression and influence DNA repair pathways^63,64^.

**Figure 9:**
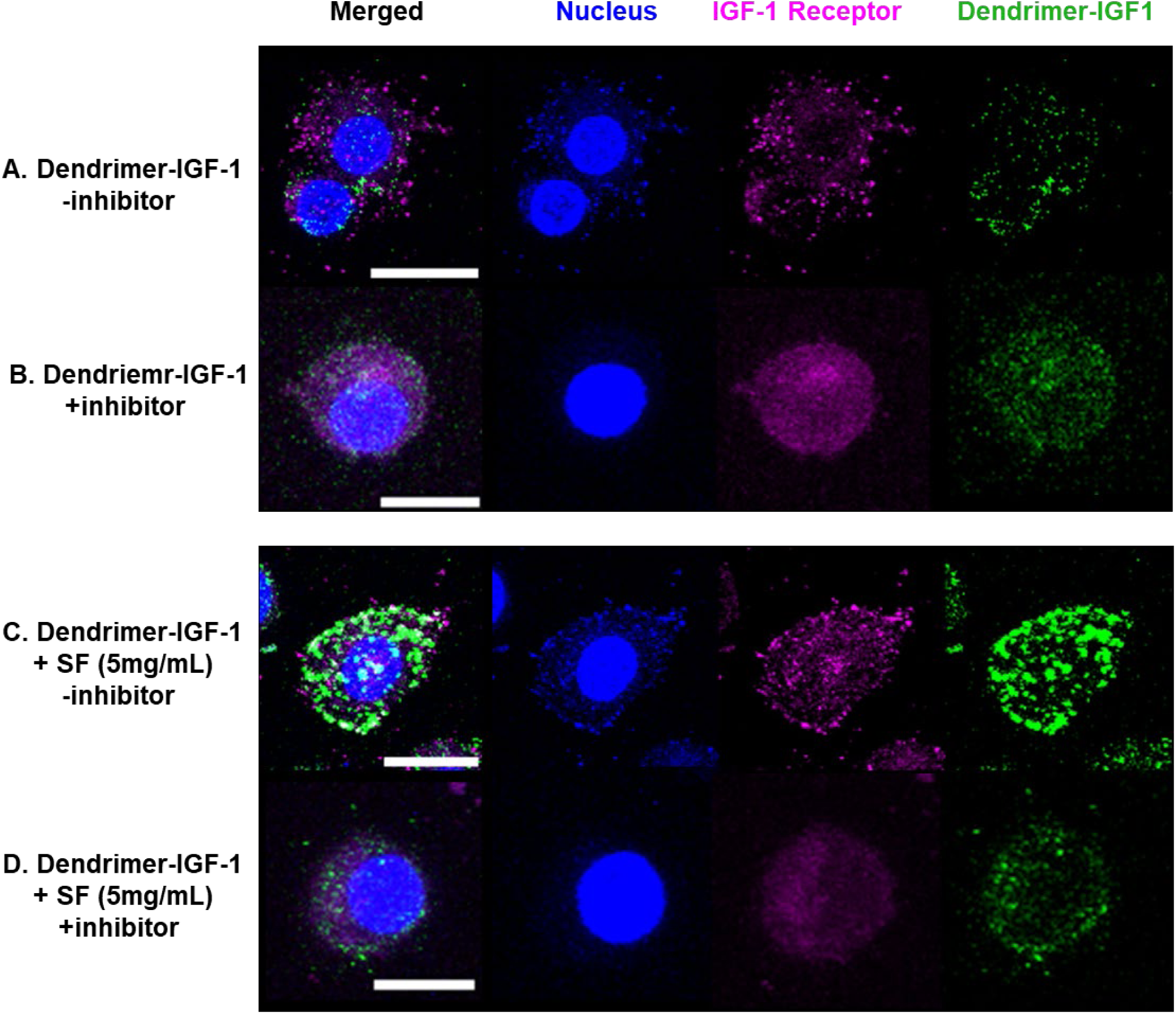
Pre-formed protein coronas aid dendrimer-IGF-1 conjugates with IGF-1 receptor engagement. Dendrimers (green) were incubated in synovial fluid for 1 hour at 37°C. For inhibitor studies, chondrocytes were treated with IGF-1 receptor antibodies for 1 hour before dosing. Confocal was done 4 hours after dosing. Nuclei were visualized with Hoechst (blue) and IGF-1 receptor was visualized with a 488 conjugated secondary antibody. Full images in Supplemental Figure 12. Dendrimer-IGF-1 conjugates are internalized by chondrocytes **(A)** with and **(B)** without IGF-1 inhibitor, suggesting dendrimers do not need to engage with IGF-1 receptors to be internalized. **(C)** However, dendrimer-IGF-1 conjugates with a pre-formed synovial fluid protein corona aggregate to co-localize more with the receptor and membrane (see Supplemental Figure 13); **(D)** this engagement is reduced with IGF-1 inhibitor. Collectively, this demonstrates that protein coronas can improve receptor-ligand targeting. | Scale bar = 10µm.

Both dendrimer-IGF-1 with and without pre-formed synovial fluid protein coronas were internalized by chondrocytes. Results suggest that dendrimer-IGF-1 conjugates without synovial fluid were not engaging with the IGF-1 receptors to be internalized (**Figure 9A**). The dendrimer-IGF-1 conjugates incubated with synovial fluid had more engagement with the IGF-1 receptors, as demonstrated by the punctate spots colocalized with IGF-1 receptor and membrane (**Figure 9C and Supplemental Figure 10**). Taken together, this suggests that the adsorbed synovial fluid proteins help with dendrimer-IGF-1 engagement with the IGF-1 receptor on chondrocytes. We hypothesize that dendrimer-IGF1 without synovial fluid is internalized by chondrocytes due to cationic interactions with the membrane. Further, these results suggest that cationic-mediated internalization could be faster than IGF-1 engagement.

To test this hypothesis, experiments were repeated by treating chondrocytes with IGF-1 receptor antibodies to act as an inhibitor to prevent IGF-1 ligand-receptor interactions. Note that, per the manufacturer (LSBio), the epitope for the IGF-1 Receptor Antibody is pTyr1161, and when IGF-1 binds to IGF-1 receptor, it leads to autophosphorylation of the tyrosine kinase (TK) domain^65^ which suggests that this antibody is a suitable inhibitor of IGF-1. We hypothesized that dendrimer-IGF-1 with a pre-formed synovial fluid protein corona would still be internalized through cationic interactions, even after chondrocytes are treated with an inhibitor. This is what we observed, even with inhibitor, chondrocytes internalized dendrimer-IGF1 conjugates (**Figure 9B**). The same was observed with dendrimer-IGF-1 conjugates with pre-formed synovial fluid protein coronas, where dendrimer-IGF-1 was still internalized, but fewer punctate spots were co-localized with the IGF-1 receptors (**Figure 8D**). Collectively, we demonstrated that dendrimer-IGF-1 conjugates can “bypass” the IGF-1 receptors and still be internalized. However, pre-formed coronas can help with dendrimer-IGF-1 engagement with IGF-1 receptors. We also assessed the effect of protein coronas on dendrimer-IGF-1 conjugates on bioactivity, as measured by percentage of cells in S phase/DNA synthesis (**Supplemental Figure 11**) and determined that preformed synovial fluid protein coronas do not change the bioactivity of dendrimer-IGF-1 conjugates.

## 4. Discussion

In this work, we characterized synovial fluid protein coronas on PEGylated PAMAM dendrimers and assessed how dendrimer-protein interactions altered biological outcomes of dendrimer uptake by cartilage and chondrocytes. PAMAM dendrimers are ideal nanocarriers to overcome delivery challenges to cartilage to deliver therapeutics to treat osteoarthritis. Analysis of dendrimer delivery efficacy must consider how synovial fluid-derived proteins and biomolecules interact with dendrimers. We hypothesized that after injection into the intra-articular space, synovial fluid-derived proteins can adsorb to the surface of nanocarriers, forming a nanocarrier-protein complex that affects targeting and transport properties. We observed that the biological outcomes of PAMAM dendrimer are determined by surface interactions between PEG and synovial fluid proteins, however, the protein corona does not always hinder biological outcomes (e.g., uptake or diffusion). Our results show that the influence of protein corona on biological outcomes with cartilage and chondrocytes depends on the large-scale interactions on the tissue or cellular level and small-scale interactions on the molecular level. Overall, protein coronas do not have a significant effect on small-scale interactions with cartilage; this includes partitioning between cartilage and meniscus, and net diffusion through cartilage. However, protein coronas significantly affect large-scale interactions with cartilage and chondrocytes; this includes uptake, desorption (critical salt concentration), uptake kinetics, and cell-internalization pathway.

When characterizing the protein corona, we observed that dendrimers mostly interact with albumin, as well as other transport proteins (serotransferrin) and complement proteins, including complement C3 and C4 gamma chain. In proof-of-concept studies^28^, we characterized dendrimer interactions with FBS proteins. Although the dendrimer concentrations are the same, we see how changing the biological fluid alters protein coronas. We observed there was an overlap of 4 top hit proteins between FBS and synovial fluid: Complement C3, Cluster of Serotransferrin, Cluster of Alpha-2 macroglobulin, and ceruloplasmin. Despite FBS and synovial fluid having 117 overlapping proteins (∼55%), there is a difference in banding patterns observed in native PAGE, suggesting dendrimer-protein interactions vary based on biological fluid. Nevertheless, regardless of the biological fluid, dendrimers interact with complement proteins, which remains consistent with previously published results^28,35^. A notable difference is that in synovial fluid, dendrimers interact more with albumin, despite both stock solutions of FBS and synovial fluid containing ∼22% albumin. It is important to mention that healthy synovial fluid is highly viscous and has zero shear viscosities ranging from 1 to 175 Pa^66^. We hypothesize that differences in albumin binding in synovial fluid are due to the higher viscosity of the synovial fluid, which can help drive electrostatic interactions between negatively charged albumin and positively charged dendrimers^67^. An interesting observation is that G4P22-34% had the highest amount (total percent spectral counts) of albumin, even compared to unPEGylated dendrimer, G4-0%. A group that used molecular docking to understand how PEG interacts with albumin demonstrated that the affinity of PEG to albumin increased as PEG molecular weight increased, and PEG at ∼10kDa was needed for PEG-albumin interactions^68^. Although the PEG used in our dendrimer conjugates (PEG8 = 550Da and PEG22 = 1000Da) is lower than the minimum molecular weight, our dendrimers contain multiple PEG chains per dendrimer, which would interact with albumin differently than the standard linear PEG used in the aforementioned molecular docking studies. We hypothesize that PEG-BSA binding affinities could explain why we see an increase in albumin in these protein coronas. Collectively, this finding demonstrates how biological environments can influence protein corona formation; these factors should be considered as we introduce dendrimers to osteoarthritic synovial fluid, which has different protein compositions (e.g., upregulated matrix metalloproteases and proinflammatory cytokines) and lower viscosity compared to healthy synovial fluid^69^.

We observed there were proteins identified in dendrimer-synovial fluid protein coronas that were not detected in stock synovial fluid (**Supplemental Figure 3**). In other words, the dendrimers were able to capture synovial fluid proteins that were at undetectable levels for LC-MS/MS only and enrich proteins to detectable levels (when combined with electrophoresis). One protein of interest we identified was collagen type VI alpha chain. A previous study identified that type VI collagen regulates synovial joint physiology and loss of type VI collagen from chondrocytes pericellular matrix alters the mechanical environment of the chondrocytes within the territorial and interterritorial extracellular matrix, which can lead to the progression of osteoarthritis^70^. Collectively, these data demonstrate how dendrimers can enrich lower abundant or harder-to-detect proteins on their surfaces. Although mass spectroscopy is an accurate tool for proteome discovery, abundant proteins can reduce the sensitivity of mass spectroscopy for identifying low abundance proteins that could be potential biomarkers^14^. To overcome low abundance sensitivity issues, protein corona-based mass spectroscopy can be a potential approach, where dendrimers or other nanoparticles can potentially be a powerful tool that can be used to characterize proteomes of various biological fluids.

While protein coronas can be seen as “biological barriers” for drug delivery^26^, incubating nanoparticles in endogenous proteins before delivery could improve therapeutic targeting. Studies on pre-formed coronas with albumin have shown they can be protective in preventing protein adsorption and reducing complement activation^71^. Other work has shown that protein coronas have been used to shield nanoparticles to help retain their targeting ability^72^. We believe we are among the first groups to demonstrate that pre-formed coronas aid nanoparticles containing targeting proteins with engagement of their cellular receptors. When IGF-1 conjugated dendrimers (dendrimer-IGF-1) were pre-incubated in synovial fluid to form a protein corona, dendrimers were internalized by chondrocytes through engagement with IGF-1 receptors. Without a pre-formed corona, IGF-1 dendrimers were still internalized, however, it was not via receptors. We determined that protein coronas can reduce net dendrimer uptake and this is likely due to the protein corona inhibiting cationic-mediated internalization.

This work bridges the gap between *in vivo* and *in vitro* studies where we can better understand how protein interactions affect the biological outcomes of dendrimers. Our work characterized bio-nano interactions throughout the dendrimer delivery pathway-from synovial fluid to the cartilage interface, diffusion through cartilage, and chondrocytes. We better understand the role PEG chain length and density (%) have in mitigating dendrimer-protein interactions and can use this knowledge to select nanocarriers with improved therapeutic efficacy. Overall, we have a better understanding of the effects of the protein corona in a pre-clinical model which can help improve translation of therapeutics.

## 5. Conclusion

In 2022, FDA guidelines reported that nanomaterials could develop different biological properties after delivery that affect their safety and efficacy, and differences in nanomaterial properties are partly due to interactions with plasma proteins forming a protein corona that impacts targeting and transport^73,74^. Our work supports these observations and those reported in literature, and strengthens the importance of characterizing protein coronas on nanoparticles in environments that mimic their delivery location. Overall, we determined that dendrimer uptake into cartilage is a balance between PEG, protein, and cartilage or chondrocyte interactions. Dendrimers mostly interact with albumin in synovial fluid. PEG helps reduce the amount of protein corona formation on dendrimers. Further, PEG shields proteins from adsorption, which has a positive impact on the dendrimers as a therapeutic protein delivery system. However, we also demonstrated the potential benefits of protein coronas for diffusion through cartilage and ligand-receptor engagement on chondrocytes. Specifically, protein adsorption may aid in the function of dendrimers as a therapeutic protein delivery system. This work is evidence that the drug delivery and nanomedicine fields need to better understand how the biological environment alters the intrinsic surface properties of nanocarriers and how protein coronas change biological outcomes (i.e.: targeting and transport) after nanoparticle delivery. Understanding these mechanisms can improve the design of nanocarriers and optimize their targeting and transport capabilities.

## Acknowledgments

The authors would like to thank and acknowledge Rick Schiavoni and the Biopolymers & Proteomics in the Swanson Biotechnology Center at the Koch Institute for Integrative Cancer Research at MIT for preparing and running samples on mass spectroscopy, as well as providing the protocol for the Materials and Methods section. The authors would also like to extend a special thank you to Tamara Dacoba and Margaret Billingsley for sending data files and checking vendor information for the material and methods section, as well as assisting with DLS measurements. Also, thank you to Brandon Johnston and Justin Kaskow for the critical review of this article. Figures created with BioRender.

## Author Contributions

Conceptualization and Methodology: SDG. Investigation and Validation: SDG, JAA (dendrimer-synovial fluid incubation and electrophoresis, uptake and salt screening assays), VMD (meniscus and cartilage co-uptake studies), BM (dendrimer-synovial fluid incubation and electrophoresis), BMJ (meniscus and cartilage co-uptake studies and confocal microscopy for cartilage diffusion studies), and JHP (assistance with bioactivity studies and confocal microscopy for chondrocyte studies). Formal Analysis: SDG, VMD (meniscus and cartilage co-uptake studies), and BMJ (meniscus and cartilage co-uptake studies). Supervision: PTH and AJG. Writing-original draft: SDG. Writing-review and editing: SDG, JAA, VMD, BM, BMJ, JHP, PTH, and AJG. Project funding acquisition: SDG, PTH, and AJG.

## Competing Interests

PTH is a member of the Board of Alector Therapeutics and the Board of Sail Biomedicine, a Flagship company, and a former member of the Scientific Advisory Board of Moderna Therapeutics and the Board of LayerBio. AJG is a consultant to NASA and NGMBio. All other authors report no competing interests.

## Funding Sources

All authors acknowledge financial support from the National Institutes of Health (NIH-NIBIB R01 EB026344) and the Department of Defense (DOD W81XWH2010481). SDG acknowledges financial support from National Institutes of Health Diversity Supplement (3-R01-EB026344-03S1), Ford Foundation Postdoctoral Fellowship, and Burroughs Wellcome Fund Postdoctoral Enrichment Program (Request ID # 1021694). JAA, VMD, and BM acknowledge financial support from the MIT Undergraduate Research Opportunity (UROP) office.

## Author Information

### Corresponding Author

Paula T. Hammond

Department of Chemical Engineering, Massachusetts Institute of Technology,

77 Massachusetts Ave, Cambridge MA, 02139

Koch Institute for Integrative Cancer Research, 500 Main St, Cambridge MA, 02142

### Authors

Simone A. Douglas-Green

Wallace H. Coulter Department of Biomedical Engineering,

Georgia Institute of Technology and Emory University

Whitaker Building

313 Ferst Drive, Atlanta, GA 30332

Juan A. Aleman

Department of Chemical Engineering, Massachusetts Institute of Technology

77 Massachusetts Ave, Cambridge MA, 02139

Victor Damptey

Department of Biological Engineering, Massachusetts Institute of Technology

77 Massachusetts Ave, Cambridge MA, 02139

Bhuvna Murthy

Department of Biological Engineering, Massachusetts Institute of Technology

77 Massachusetts Ave, Cambridge MA, 02139

Brandon Johnston

Department of Chemical Engineering, Massachusetts Institute of Technology

77 Massachusetts Ave, Cambridge MA, 02139

Joon Ho Park

Department of Chemical Engineering, Massachusetts Institute of Technology

77 Massachusetts Ave, Cambridge MA, 02139

Alan J. Grodzinsky

Department of Biological Engineering, Massachusetts Institute of Technology

77 Massachusetts Ave, Cambridge MA, 02139

## Notes

### Summary of Updates

Spelling of author last name corrected; affiliations for authors updated

