## Supplemental Figures for "Investigating Bio-Nano Interactions of PEGylated Cationic Polyamidoamine (PAMAM) Dendrimers within Synovial Joints"

1. Representative full Stain-FreeTM native PAGE gels used to assess electrophoretic migration distance.

2. Representative full regular native PAGE gels used for mass spectroscopy analysis.

3. Proteins found associated with dendrimers but not in synovial fluid stock.

4. Optimizing a “materials characteristic based” protocol to determine how dendrimer-protein interactions affect cartilage uptake.

5. Cartilage uptake and critical salt concentration for G4P8 and G4P22 libraries incubated in varying concentrations of synovial fluid.

6. Representative images of diffusion of unPEGylated and PEGylated dendrimers with or without pre-formed synovial fluid protein coronas.

7. Proteins shield charged amines on dendrimers to facilitate weak reversible interactions within cartilage to aid dendrimer diffusion.

8. Pre-coating dendrimers with synovial fluid-protein corona improves chondrocyte* viability.

9. Confocal microscopy images from experiments with pre-formed protein coronas and dendrimer-IGF-1.

10. Single Z slice confocal microscopy images from experiments with pre-formed protein coronas and dendrimer-IGF-1.

11. Protein coronas do not reduce the bioactivity of dendrimer-IGF1 conjugates.

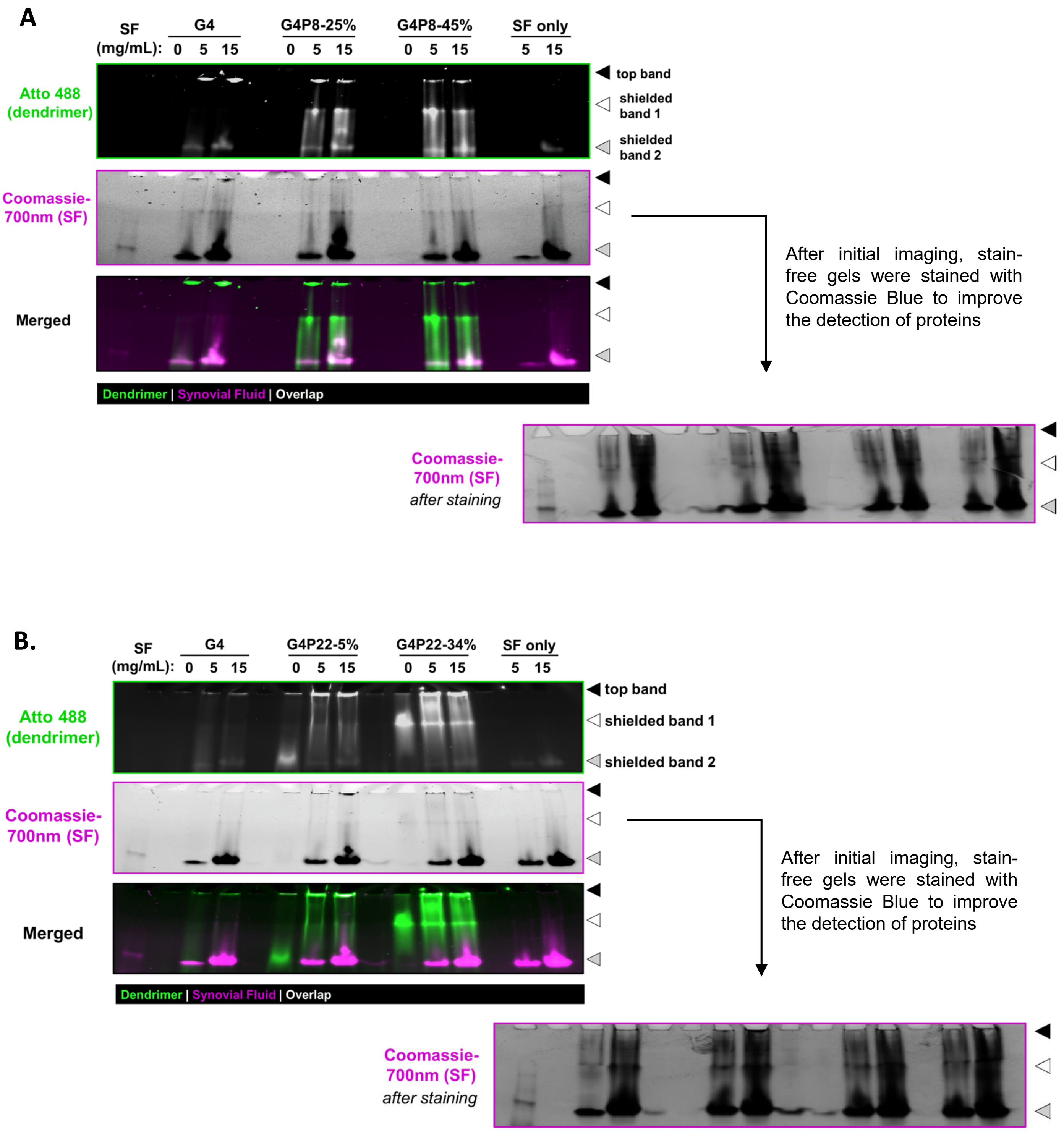

A
SF (mg/mL):
G4
0 5 15
G4P8-25%
0 5 15
G4P8-45%
0 5 15
SF only
5 15
Atto 488 (dendrimer)
top band
shielded band 1
shielded band 2
Coomassie-700nm (SF)
Merged
Dendrimer | Synovial Fluid | Overlap
After initial imaging, stain-free gels were stained with Coomassie Blue to improve the detection of proteins
Coomassie-700nm (SF) after staining
B.
SF (mg/mL):
G4
0 5 15
G4P22-5%
0 5 15
G4P22-34%
0 5 15
SF only
5 15
Atto 488 (dendrimer)
top band
shielded band 1
shielded band 2
Coomassie-700nm (SF)
Merged
Dendrimer | Synovial Fluid | Overlap
After initial imaging, stain-free gels were stained with Coomassie Blue to improve the detection of proteins
Coomassie-700nm (SF) after staining

### Densitometry for G4PEG8

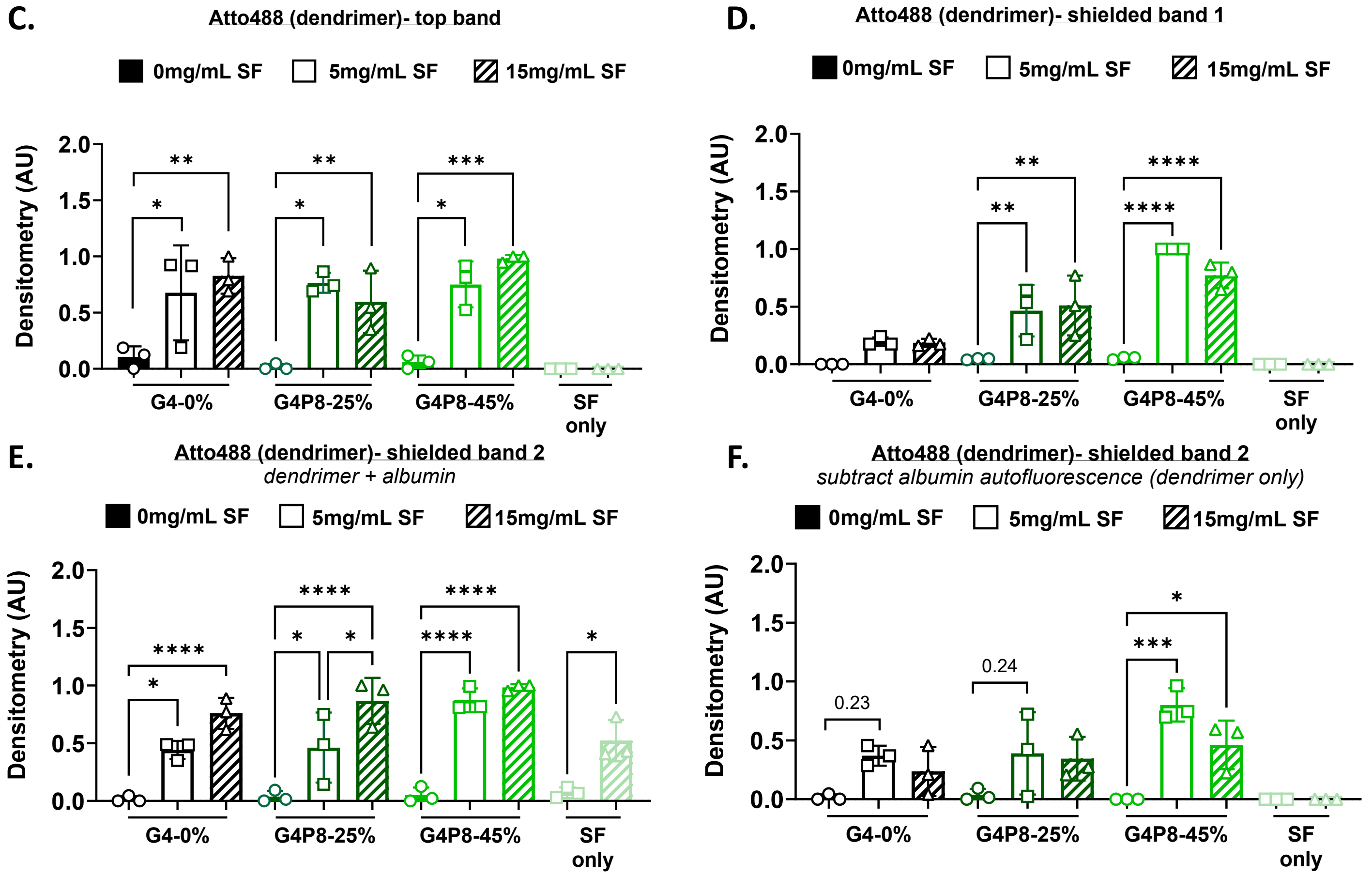


### Densitometry for G4PEG22

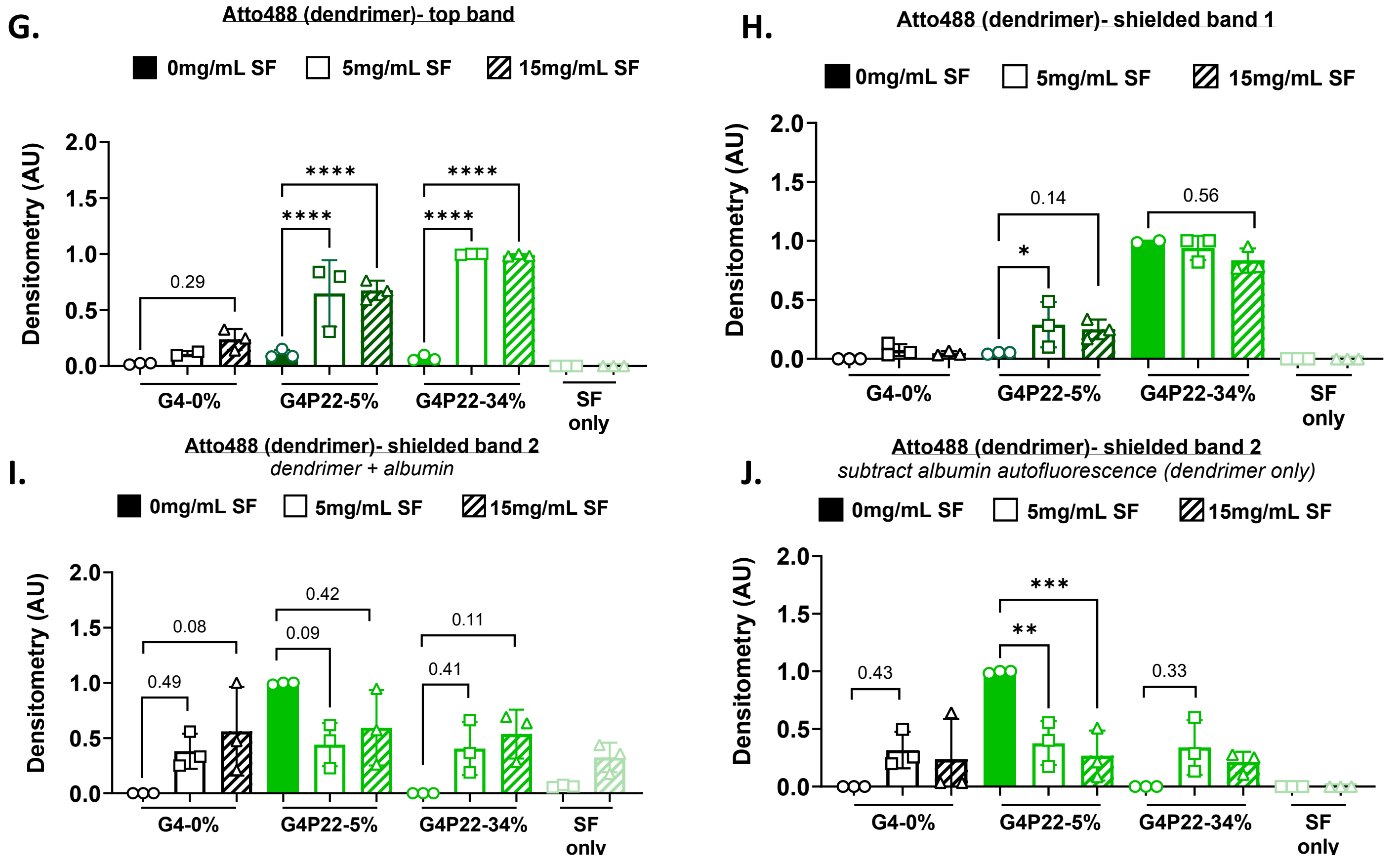

**Supplemental Figure 1: Representative full Stain-Free™ native PAGE gels used to assess electrophoretic migration distance.** Stain-Free™ gels are used to assess the electrophoretic migration of the dendrimers was measured to determine interactions between **(A)** short-chain PEGylated dendrimers or **(B)** long-chain PEGylated dendrimers and synovial fluid proteins. For **(C-E)** short and **(G-I)** long chain PEGylated dendrimers, Fluorescence for the top, shielded band 1, and shielded band 2 in the Atto 488/dendrimer channel (data normalized to max intensity for each gel) was quantified with densitometry. **(F/J)** Shielded band 2 (grey arrow) was identified in all dendrimer formulations; this band auto-fluoresces in the synovial fluid only control and was later confirmed to be albumin with mass spectroscopy. To determine whether fluorescence in shielded band 2 was due to dendrimer and/or albumin auto-fluorescence, the fluorescence from the albumin in the synovial fluid only channel was subtracted from the experimental groups, which confirmed that dendrimer associates with albumin.

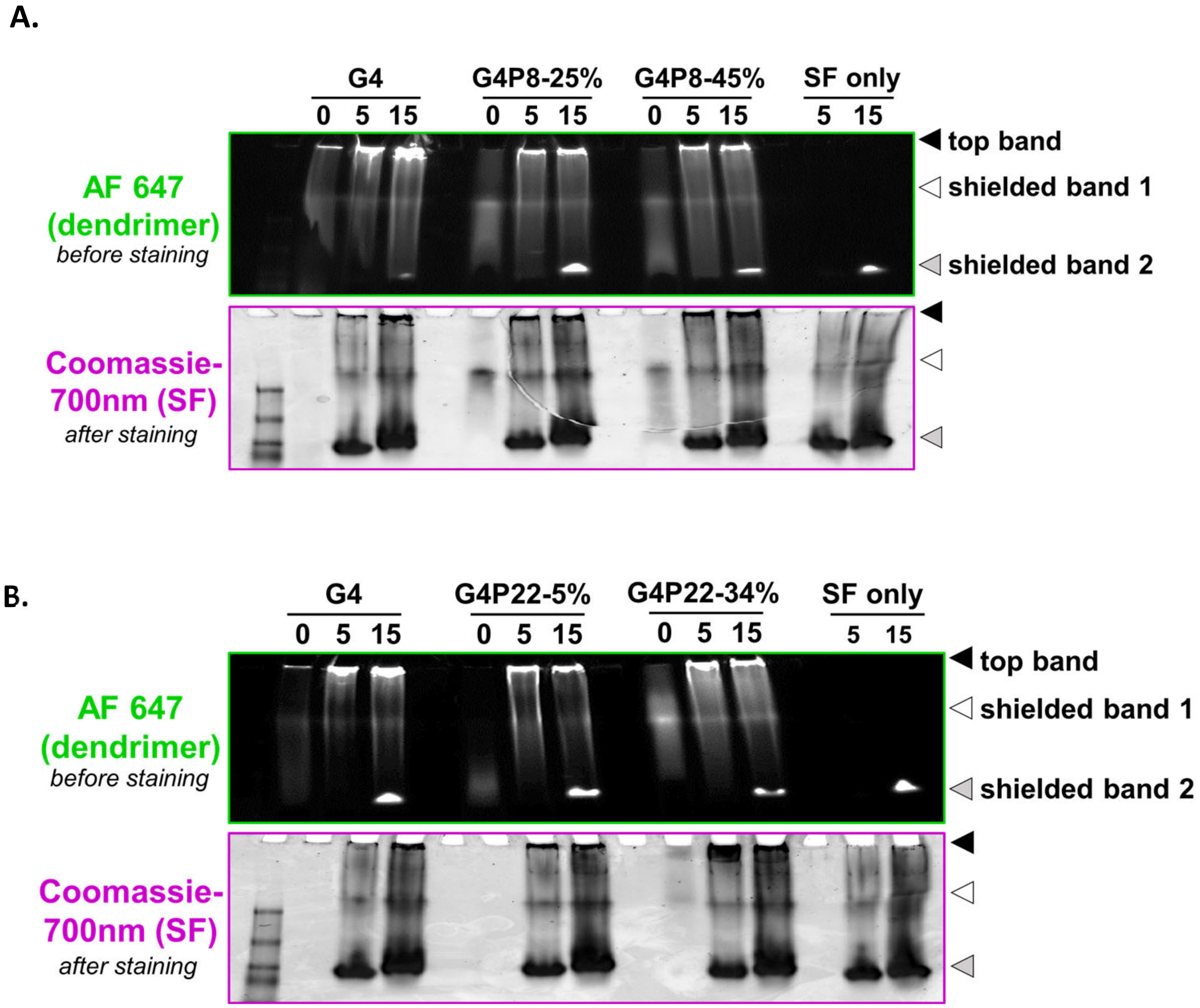


**Supplemental Figure 2: Representative full regular native PAGE gels used for mass spectroscopy analysis.** Regular gels are used for mass spectroscopy analysis to identify proteins, where the top and shielded bands are isolated, digested, and run on mass spectroscopy.

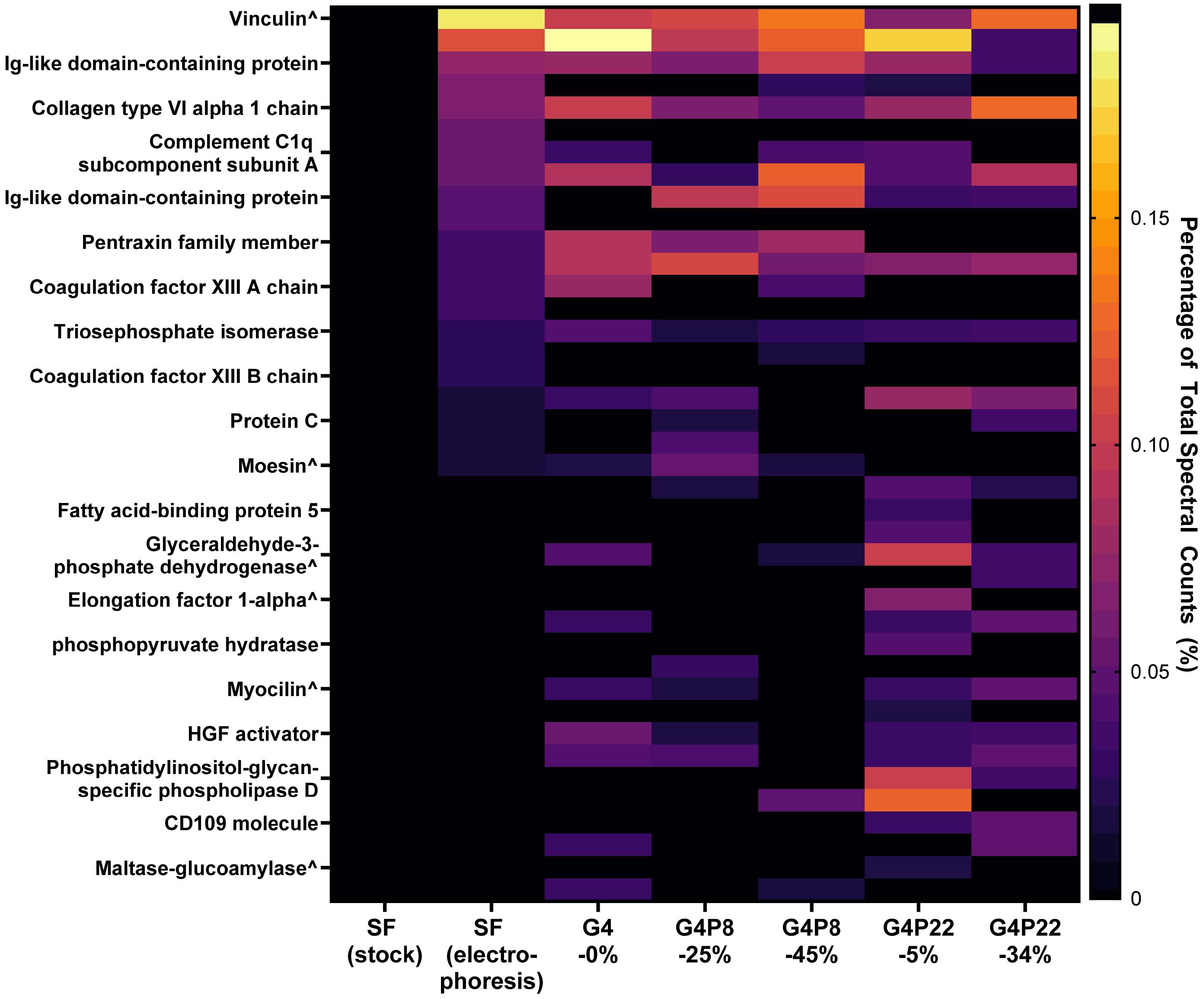


**Supplemental Figure 3: Proteins found associated with dendrimers but not in synovial fluid stock.** Note there were also some proteins not found in either synovial fluid stock or “electrophoresis” control.

^ *denotes found in cluster*

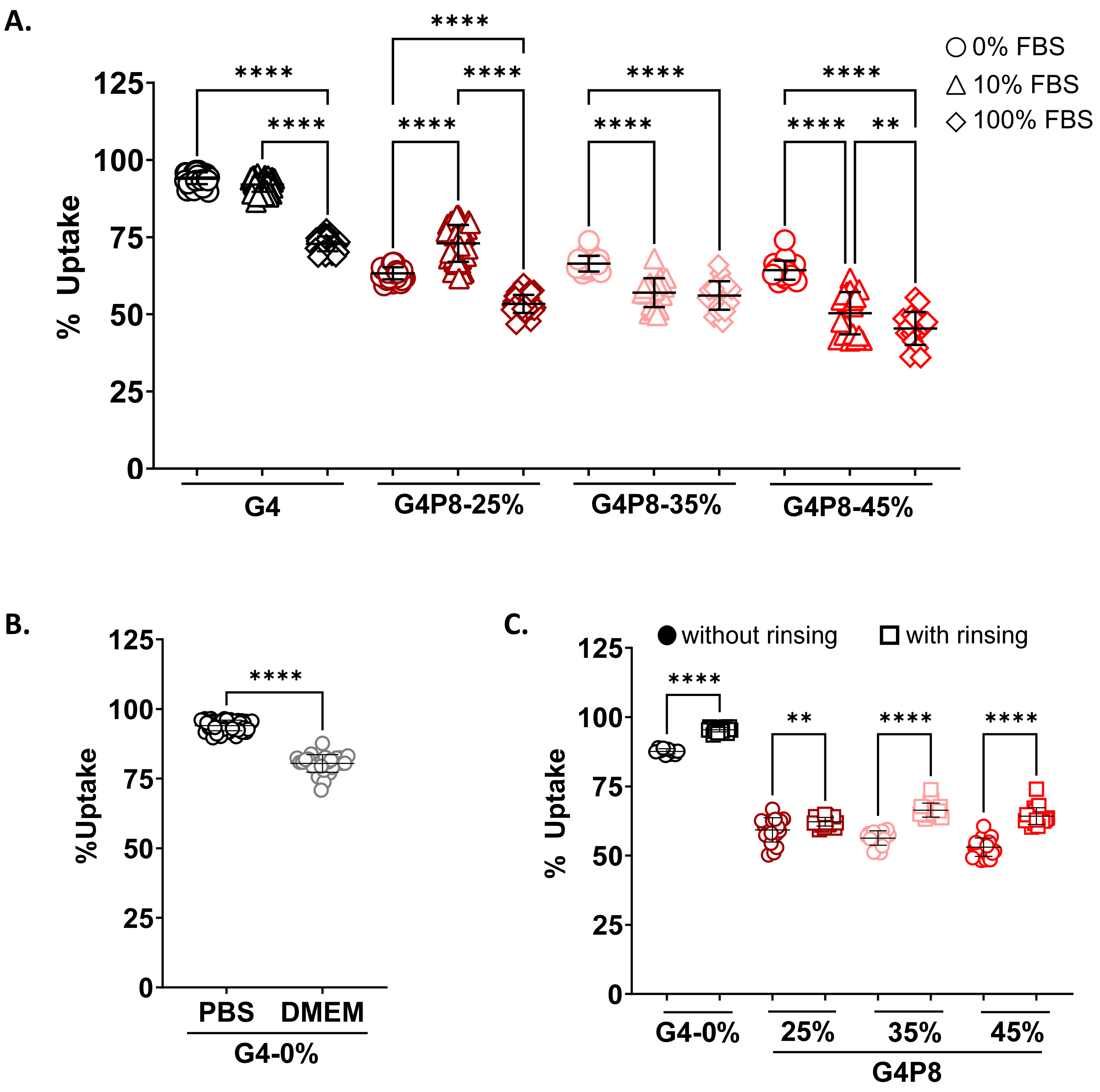


**Supplemental Figure 4: Optimizing a "materials characteristic-based" protocol to determine how dendrimer-protein interactions affect cartilage uptake.** To understand dendrimer-protein interactions and how that affects uptake from a material characteristic perspective we aimed to eliminate and control variables as best as possible. **(A)** FBS was used to troubleshoot the uptake protocol; with notable differences in percent uptake compared to synovial fluid (Figure 3) which demonstrates differences in biological outcomes (i.e.: uptake) of the same nanocarrier in different biological fluids. **(B)** For initial experiments, DMEM without any factors or FBS was used, but it was observed that the nutrients and small biomolecules in DMEM significantly reduced uptake relative to PBS which had nearly 100% uptake. This suggests factors in the DMEM affect the uptake of unPEGylated dendrimer. **(C)** While optimizing the protocol, we determined it was important to rinse cartilage plugs, prior to adding dendrimers. Cartilage explants are kept in DMEM supplemented with FBS and other proteins. We hypothesized that , when not rinsed off, surface proteins prevent uptake. More importantly, rinsing plugs eliminates variables to further characterize dendrimer-protein interactions from a material characteristic perspective.

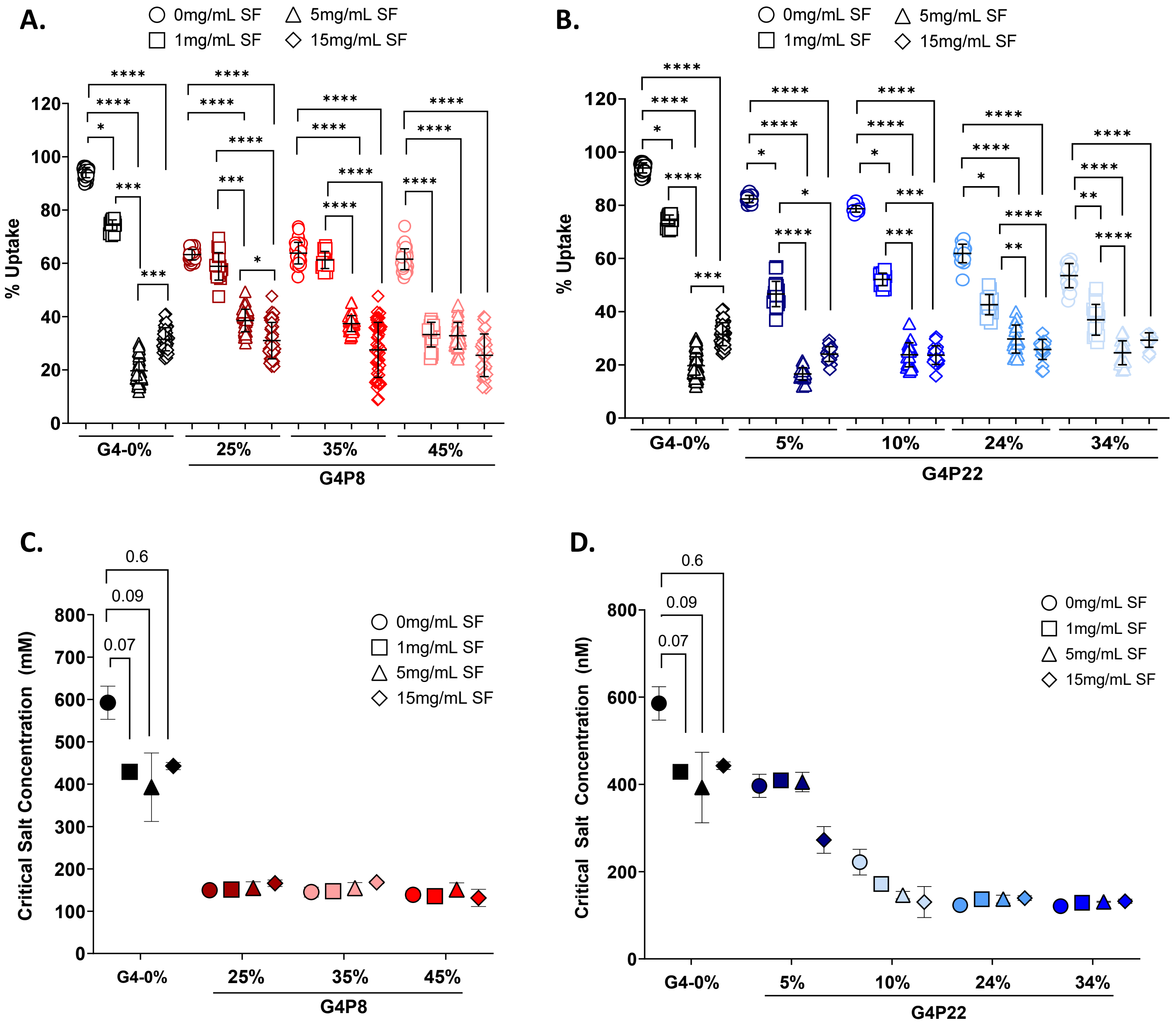


**Supplemental Figure 5: Cartilage uptake and critical salt concentration for G4P8 and G4P22 libraries incubated in varying concentrations of synovial fluid.** Fluorescently tagged dendrimers with varying chain lengths, **(A)** PEG8 or **(B)** PEG22, and densities were incubated in varying concentrations of synovial fluid (SF) for 1 hour at 37°C to form a protein corona, then added to bovine cartilage explants for 24 hours and percent uptake was quantified using fluorescence. Dendrimer uptake into cartilage decreases in synovial fluid, however this is mitigated by varying PEG chain length or density. *2-4 biological replicates with 10 technical replicates; ANOVA, Kruskal-Wallis Test for each dendrimer formulation | ****p<0.0001, ***p<0.001, **p<0.05* **(C and D)** After uptake, a salt screening assay was done by placing cartilage explants in increasing PBS concentrations (0.5X to 9X) for 24 hours to measure dendrimer desorption and calculate critical salt concentration (CSC). For unPEGyalted dendrimers, there is a decrease in the critical salt concentration, suggesting protein adsorption to dendrimer is blocking ACAs on the surface. However, for PEGylated dendrimers, despite synovial fluid proteins and biomolecules being present, there is little to no change in critical salt concentration, which suggests PEG shields dendrimers from protein adsorption while sustaining electrostatic interactions with cartilage. *2 biological replicates with 2 technical replicates; ANOVA, Kruskal-Wallis Test for each dendrimer formulation.*

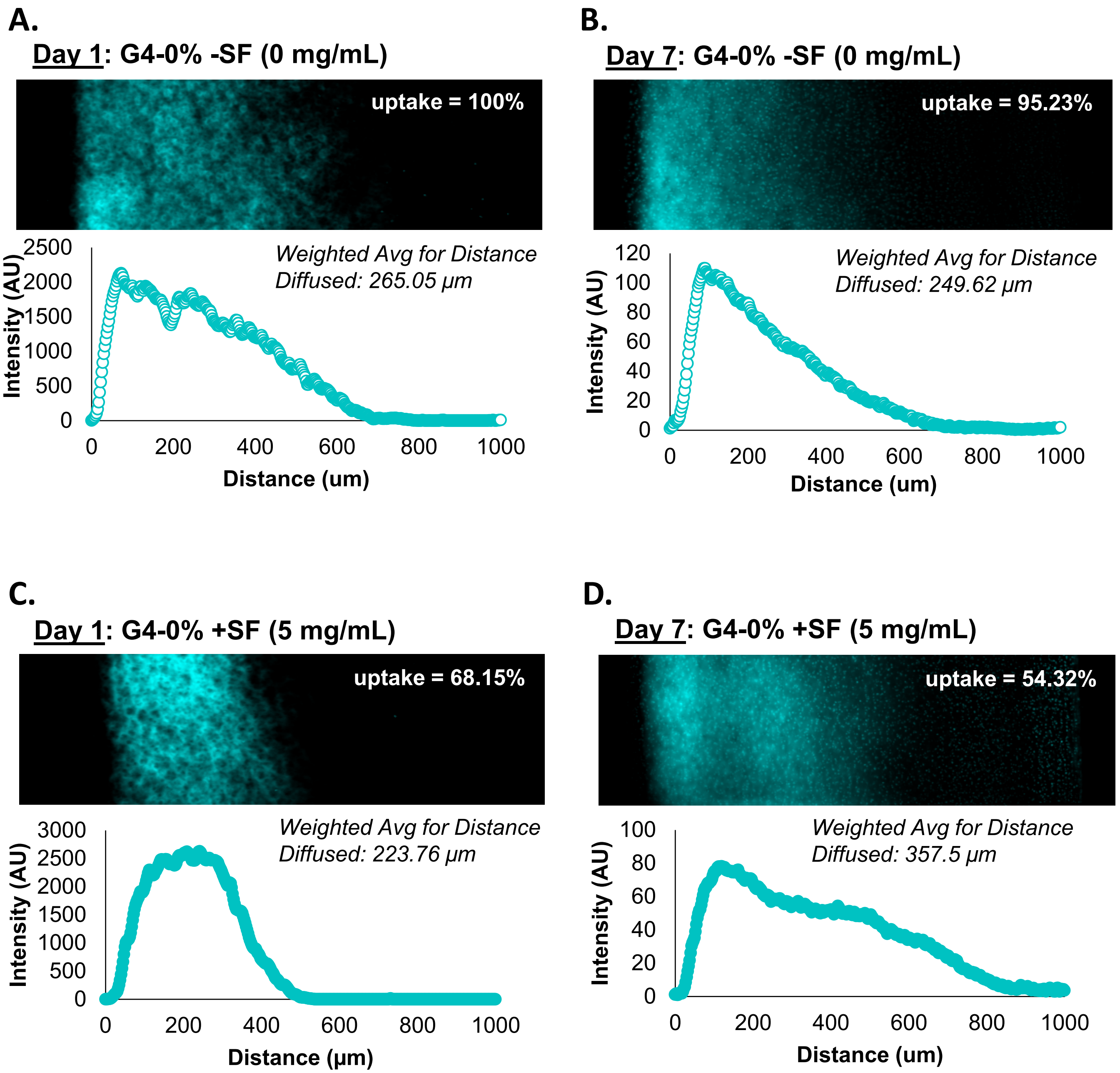

A.
Day 1: G4-0% -SF (0 mg/mL)
uptake = 100%
Weighted Avg for Distance Diffused: 265.05 μm
Intensity (AU)
Distance (um)
B.
Day 7: G4-0% -SF (0 mg/mL)
uptake = 95.23%
Weighted Avg for Distance Diffused: 249.62 μm
Intensity (AU)
Distance (um)
C.
Day 1: G4-0% +SF (5 mg/mL)
uptake = 68.15%
Weighted Avg for Distance Diffused: 223.76 μm
Intensity (AU)
Distance (μm)
D.
Day 7: G4-0% +SF (5 mg/mL)
uptake = 54.32%
Weighted Avg for Distance Diffused: 357.5 μm
Intensity (AU)
Distance (um)

**E.**

**Day 1: G4P8-25% -SF (0 mg/mL)**

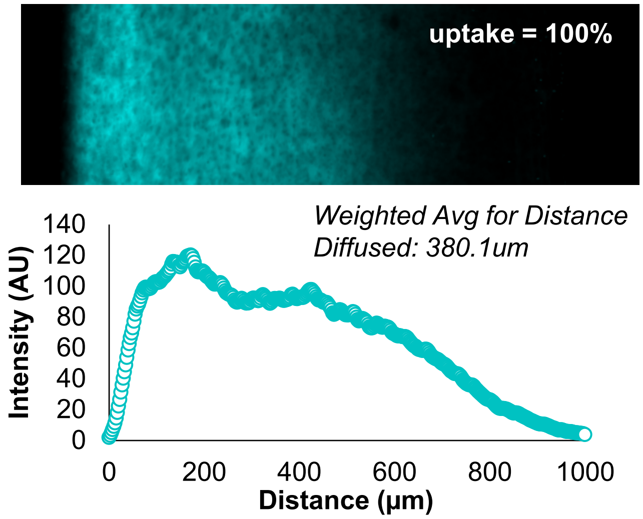


**F.**

**Day 7: G4P8-25% -SF (0 mg/mL)**

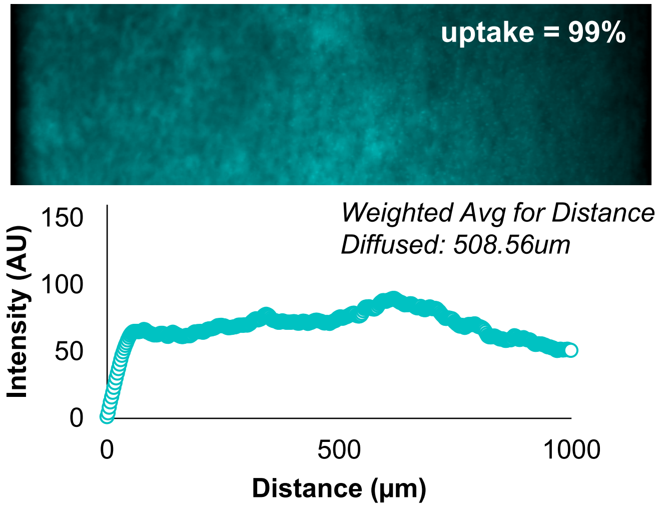


**G.**

**Day 1: G4P8-25% +SF (5 mg/mL)**

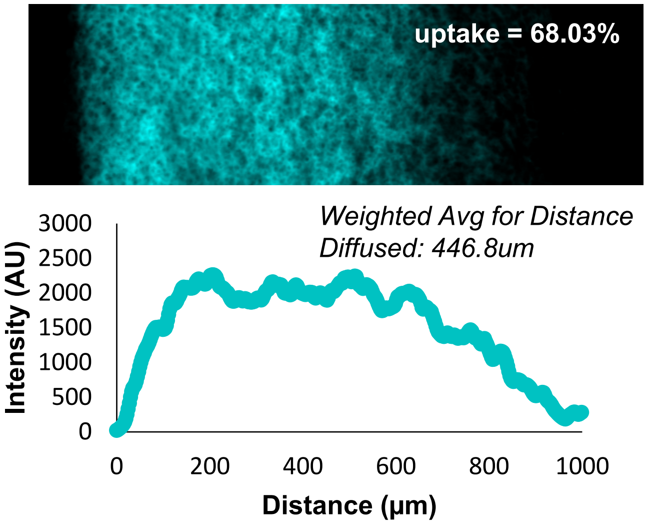


**H.**

**Day 7: G4P8-25% +SF (5 mg/mL)**

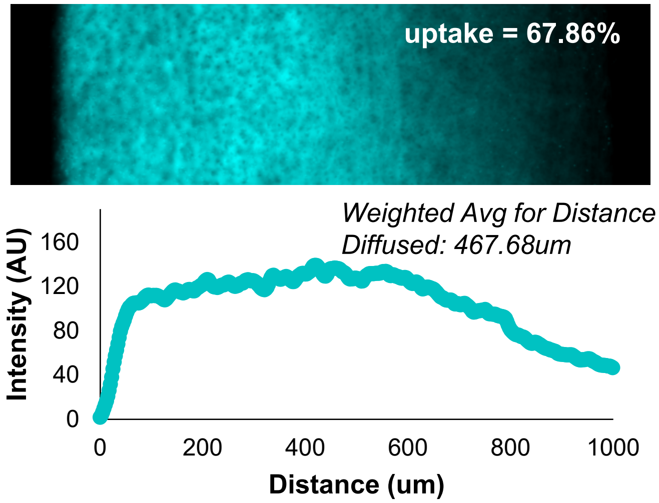

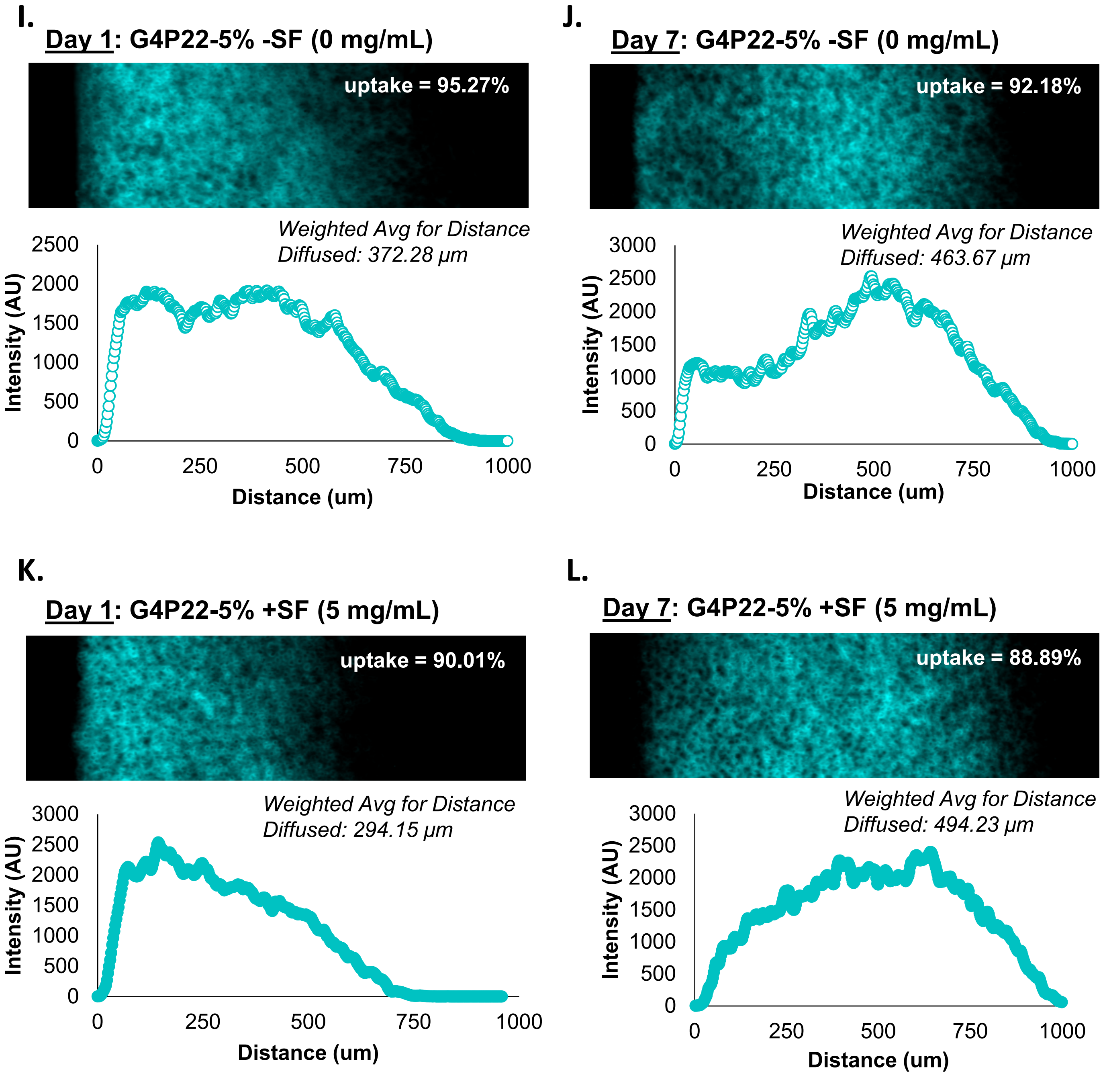


**Supplemental Figure 6: Representative images of diffusion of unPEGylated and PEGylated dendrimers with or without pre-formed synovial fluid protein coronas. (A-D)** UnPEGylated, **(E-H)** short, and **(I-L)** long (chain PEGylated dendrimers were incubated in synovial fluid for 1 hour, then added to cartilage explants. After 24 hours or 7 days, explants were removed, sectioned, and imaged on microscopy to analyze dendrimer diffusion through cartilage. After 24 hours and 7 days, there was no significant increase or decrease or in average diffusion depth for dendrimer formulations with or without synovial fluid, which suggests protein corona does not hinder dendrimer diffusion in cartilage.

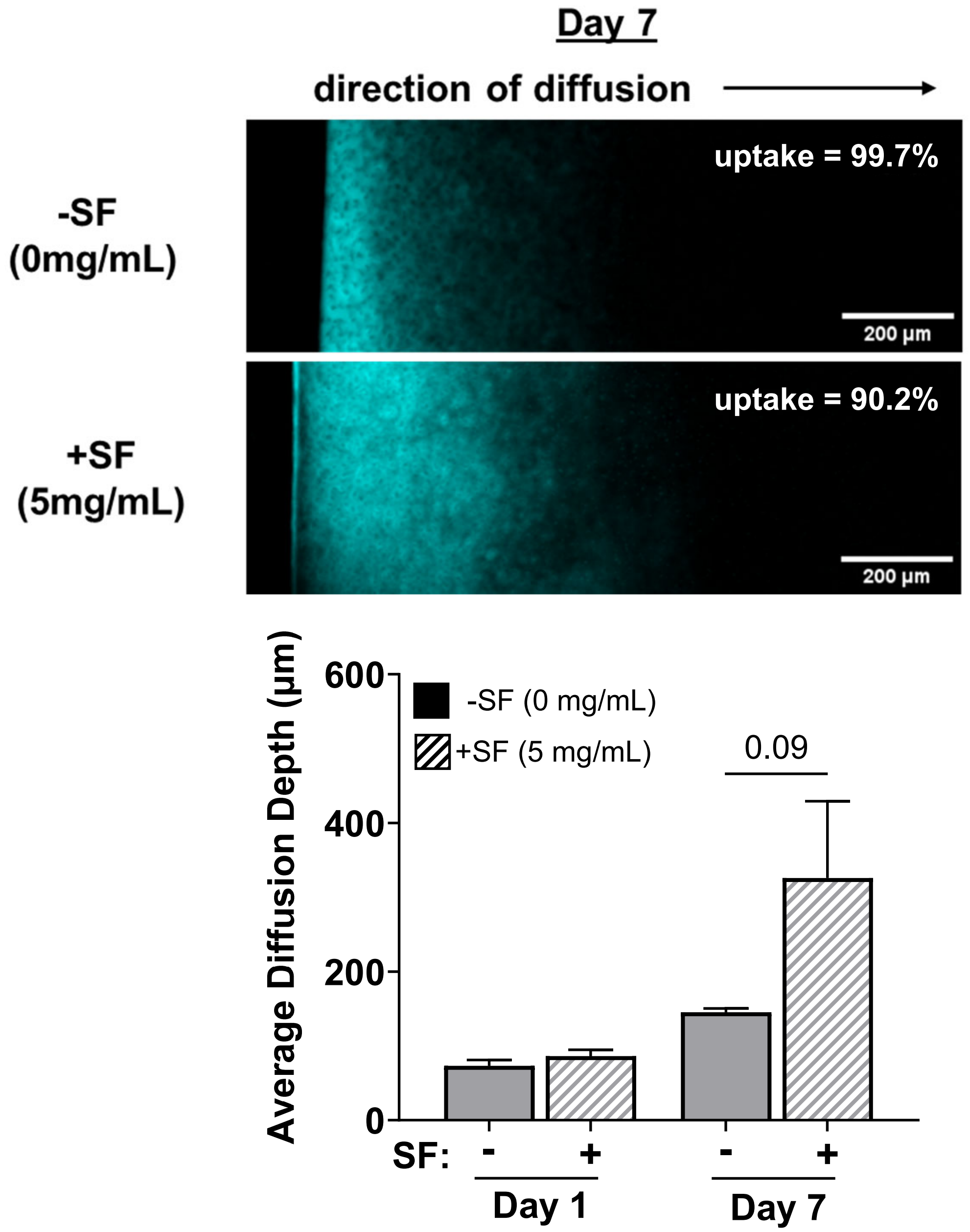


**Supplemental Figure 7: Proteins shield charged amines on dendrimers to facilitate weak reversible interactions within cartilage to aid dendrimer diffusion.** We wanted to test the hypothesis that proteins can shield charges on primary amines and facilitate weak irreversible interactions to aid in dendrimer diffusion. To do this, Generation 6 (Gen6) PAMAM dendrimers, which contain 256 primary amines, were pre-incubated in synovial fluid prior to dosing cartilage explants. Diffusion was measured at day 1 and day 7. After day 1, G6 dendrimers stick well to the surface with little diffusion in both conditions. However, with a pre-formed corona, after 7 days we see an increase ($p = 0.09$) in average diffusion depth.

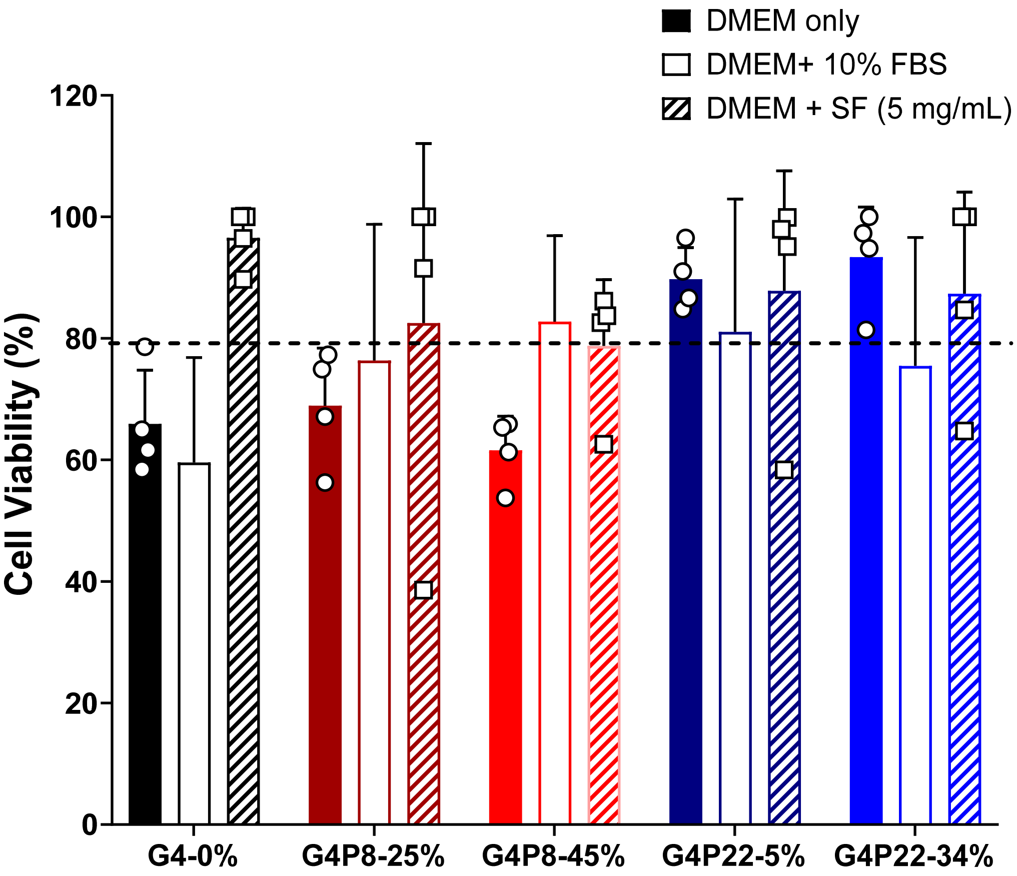


**Supplemental Figure 8: Pre-coating dendrimers with synovial fluid-protein corona improves chondrocyte* viability.** Prior to *in vitro* studies, we wanted to ensure chondrocytes in bovine synovial fluid would not be toxic to cells. Pre-coating dendrimers with synovial fluid improves cell viability for Gen4-0% and Gen4-PEG8 dendrimers. Cell viability was better in synovial fluid than FBS; hypothesize because synovial fluid proteins are more native to chondrocytes compared to FBS. This study also serves as preliminary work for pre-coating dendrimers to improve biocompatibility.

*note: ATCC human chondrocytes were used in these experiments

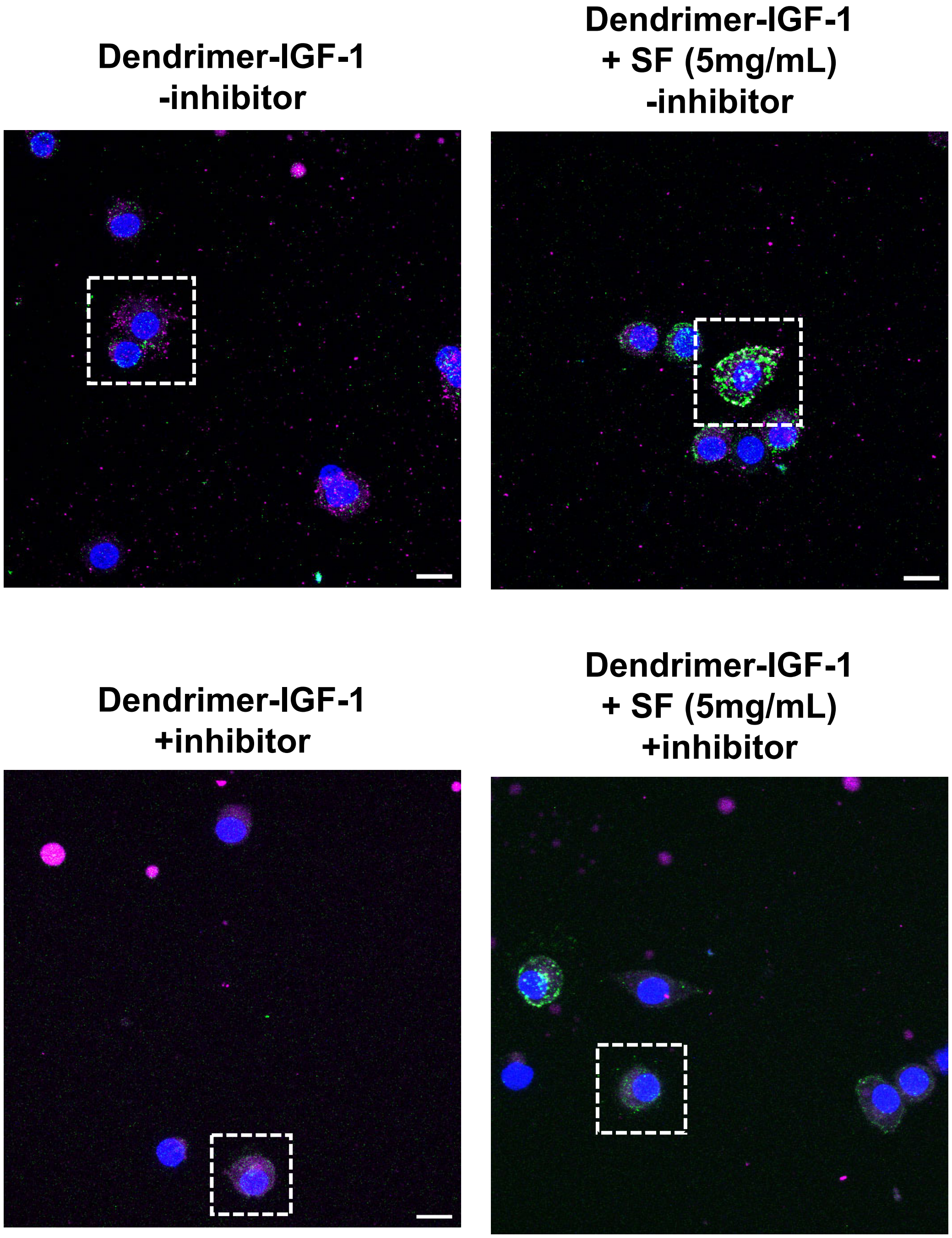


**Supplemental Figure 9: Confocal microscopy images from experiments with pre-formed protein coronas and dendrimer-IGF-1.**

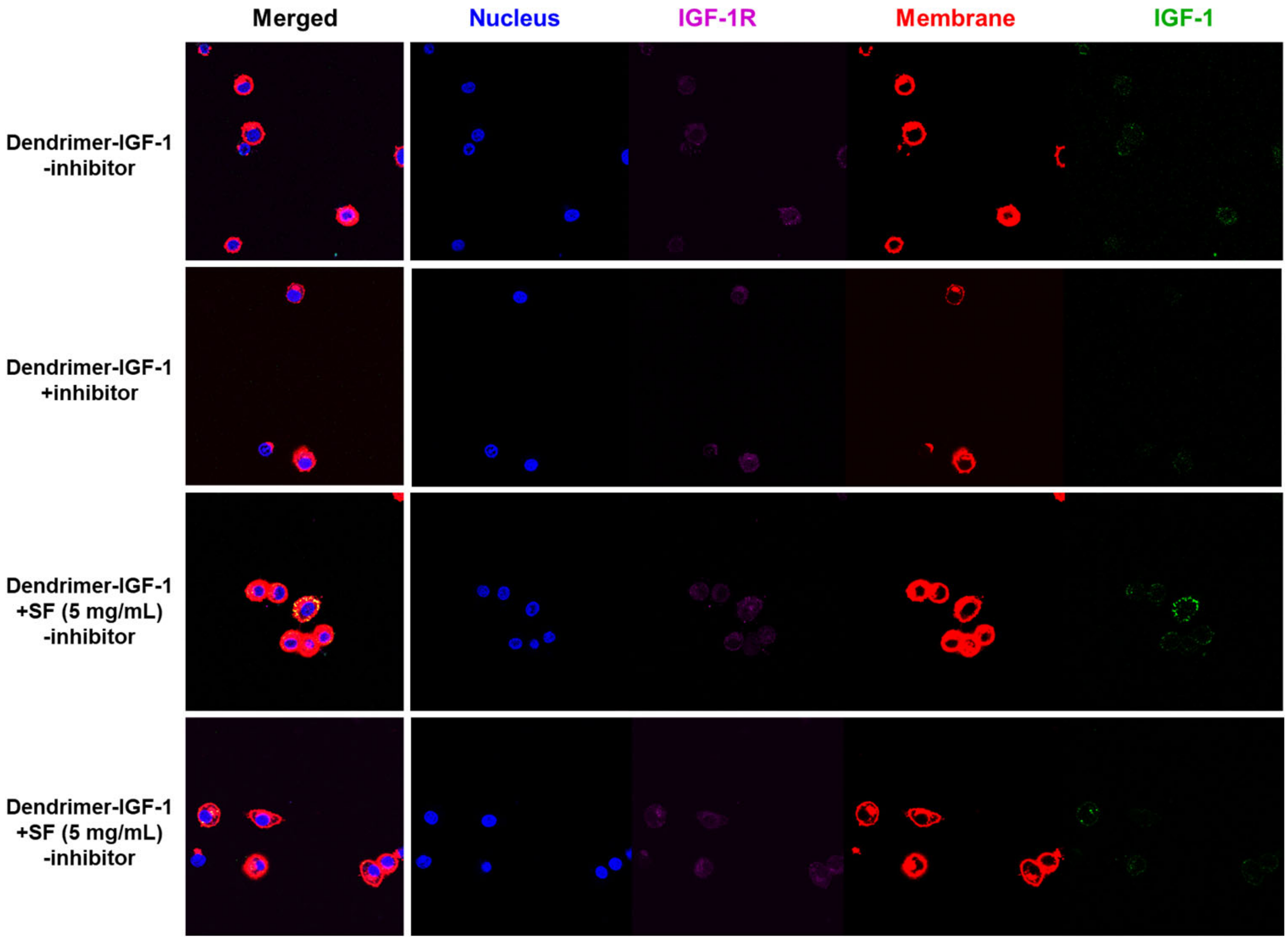


**Supplemental Figure 10: Single Z slice confocal microscopy images from experiments with pre-formed protein coronas and dendrimer-IGF-1.** This illustrates membrane (red) relative to IGF-1 (green) and confirms surface interactions of dendrimer-IGF-1 conjugates incubated with synovial fluid.

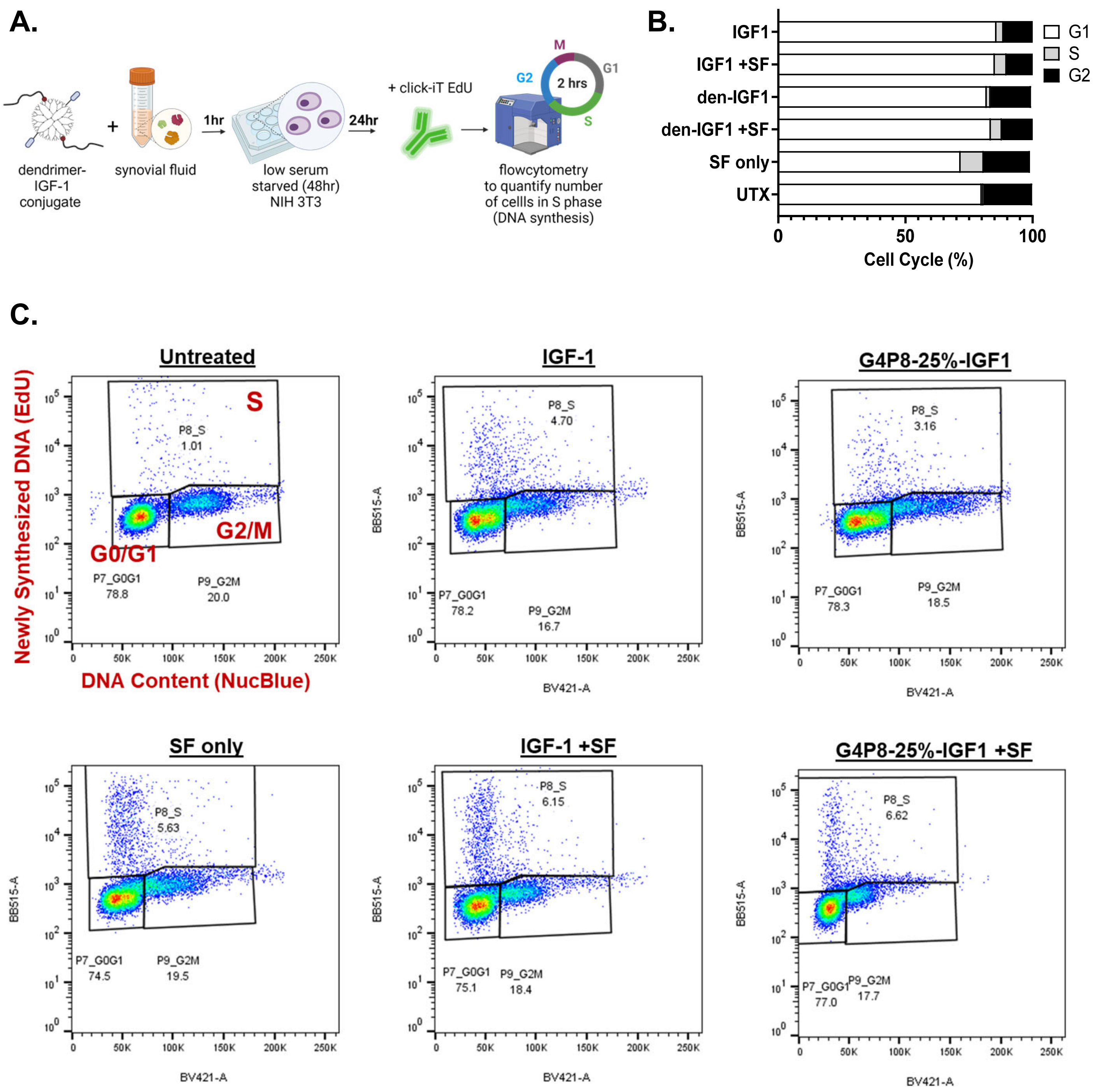
A.
dendrimer-IGF-1 conjugate
+
synovial fluid
1hr
low serum starved (48hr) NIH 3T3
24hr
+ click-iT EdU
flowcytometry to quantify number of cellls in S phase (DNA synthesis)
M
G2
2 hrs
G1
S
B.
IGF1
IGF1 +SF
den-IGF1
den-IGF1 +SF
SF only
UTX
G1
S
G2
0
50
100
Cell Cycle (%)
C.
Untreated
IGF-1
G4P8-25%-IGF1
SF only
IGF-1 +SF
G4P8-25%-IGF1 +SF
Newly Synthesized DNA (EdU)
DNA Content (NucBlue)
S
G0/G1
G2/M
P8_S 1.01
P7_G0G1 78.8
P9_G2M 20.0
P8_S 4.70
P7_G0G1 78.2
P9_G2M 16.7
P8_S 3.16
P7_G0G1 78.3
P9_G2M 18.5
P8_S 5.63
P7_G0G1 74.5
P9_G2M 19.5
P8_S 6.15
P7_G0G1 75.1
P9_G2M 18.4
P8_S 6.62
P7_G0G1 77.0
P9_G2M 17.7
BB515-A
BV421-A

**Supplemental Figure 11: Protein coronas do not reduce the bioactivity of dendrimer-IGF1 conjugates. (A)** To assess how the protein corona affects bioactivity, G4P8-25%-IGF1 conjugates or IGF-1 (conditions matched to 100ng of IGF-1) were incubated in synovial fluid for 1hr at 37C prior to dosing low serum-starved NIH 3T3 cells for 24 hours. A Click-iT EdU Cell Proliferation Assay was used to quantify new synthesized DNA by incorporating EdU into DNA during activity DNA synthesis (S phase). See below for full description of Bioactivity Assay methods. **(B)** Using flow cytometry we determined the percentage of cells in cell growth (G0/G1), DNA synthesis (S phase), and preparing for cell division (G2/M). **(C)** Representative flow cytometry plots are shown for each experimental group. Overall, we determined that pre-formed protein coronas do not impair bioactivity of dendrimer-IGF1 conjugates.

*Bioactivity Assay*: Mouse embryo fibroblasts (NIH/3T3; ATCC) were used for bioactivity assays; cultured in high serum media (DMEM, 4.5 g/L glucose with 10% FBS, 1% Pen-Strep) at 37°C in 5% CO2, and was replaced every 2-3 days. IGF-1 proliferative bioactivity was observed used using the Click-iT EdU cell proliferation kit for flow cytometry (Invitrogen) to observe the cell cycle by quantifying the percent of NIH/3T3 fibroblasts in S phase in response to IGF-1 treatment. In T-75 flasks, cells were seeded overnight in full media. The following day, media was replaced with low serum (0.5% FBS) for 48 hours. 100ng/mL of dendrimer-IGF-1 or (free) IGF-1 was incubated in 1mg/mL of synovial fluid to form a protein corona. Low serum media was aspirated and cells were rinsed with 1x PBS to remove residual proteins before being dosed with dendrimer-IGF-1 or IGF-1 with or without pre-formed protein coronas, or synovia fluid (1mg/mL) only for 24 hours. Next, cells were incubated with 10 µM EdU for 2 hours, fixed, and permeabilized following the manufacturer’s instructions. EdU was labeled with an azide-functionalized AF488 dye and cells were stained with DAPI. Flow cytometry (FACS Celesta, BD) was used to analyze Dual EdU-488/DAPI stained cells with 20,000 events per sample.
