## Supplemental Tables for "Investigating Bio-Nano Interactions of PEGylated Cationic Polyamidoamine (PAMAM) Dendrimers within Synovial Joints"

**Supplemental Table 1: Summary of dendrimer characterization**

| Dendrimer Forumulation | Dn(50) DLS, nm | Accessible Charged Amines (ACA)* | 24hr- Avg Percent Uptake | | 24 hr- Avg Diffusion Depth (um) | | 24 hr- Chondrocyte Internalization | | *In Vivo* Properties* |
|---|---|---|---|---|---|---|---|---|---|
| | | | without protein corona | with protein corona | without protein corona | with protein corona | without protein corona | with protein corona | |
| **G4-0%** | 3.99 ± 0.28 | 64 | 94.07% | 19.70% | 305.09 | 282.6 | yes | little to none, dendrimer aggregated outside cell | not tested due to cytotoxicity |
| **G4P8-25%** | 5.00 ± 0.38 | 36 | 63.25% | 38.53% | 326.34 | 391.44 | yes | reduced | short PEG chains with higher ACA had greatest joint retention (100+ days in rat knee joints) |
| **G4P8-35%** | 5.93 ± 0.24 | 29 | 63.84% | 37.42% | --- | --- | --- | --- | |
| **G4P8-45%** | 4.99 ± 0.54 | 21 | 61.57% | 32.85% | 353.07 | 440.68 | yes | reduced | |
| **G4P22-5%** | 4.37 ± 0.18 | 47 | 82.38% | 16.59% | 456.73 | 296.66 | yes | yes, dendrimer dispersion outside cell | --- |
| **G4P22-10%** | 4.84 ± 0.21 | 40 | 78.71% | 23.78% | --- | --- | --- | --- | --- |
| **G4P22-24%** | 6.35 ± 0.23 | 21 | 61.90% | 29.75% | --- | --- | --- | --- | --- |
| **G4P22-34%** | 6.63 ± 0.15 | 8 | 53.57% | 24.55% | 488.3 | 435.41 | yes | no | high fast rate constant (~2 1/days), low slow half life (~25 days), lowest joint retention |

**Supplemental Table 2: Mass spectroscopy data for dendrimers incubated in 15 mg/mL synovial fluid**

| | Identified Proteins | Accession Number | Molecular Weight (MW) | Theoretical Isoelectric Point (pI) | Biological Process | G4-0% | | G4P8-25% | | G4P8-45% | | G4P22-5% | | G4P22-34% | | Synovial Fluid (electrophoresis) | | Synovial Fluid (stock) | |
|---|---|---|---|---|---|---|---|---|---|---|---|---|---|---|---|---|---|---|---|
| | | | | | | Total Spectrum Count | Percentage of Total Spectral Counts | Total Spectrum Count | Percentage of Total Spectral Counts | Total Spectrum Count | Percentage of Total Spectral Counts | Total Spectrum Count | Percentage of Total Spectral Counts | Total Spectrum Count | Percentage of Total Spectral Counts | Total Spectrum Count | Percentage of Total Spectral Counts | Total Spectrum Count | Percentage of Total Spectral Counts |
| 1 | Albumin OS=Bos taurus OX=9913 GN=ALB PE=4 SV=1 | A0A140T897 | 69 kDa | 5.77 | Transport/Carrier/Binding | 1556 | 18.034% | 1633 | 18.138% | 1655 | 17.375% | 1573 | 18.306% | 1583 | 20.807% | 1662 | 16.147% | 2333 | 22.028% |
| 2 | Cluster of Endopin 2 OS=Bos taurus OX=9913 GN=SERPINA3-7 PE=1 SV=1 (A0A0A0MP92) | A0A0A0MP92 [2] | 47 kDa | 5.61 | Protease Inhibitor | 94 | 1.089% | 112 | 1.244% | 106 | 1.113% | 103 | 1.199% | 145 | 1.906% | 148 | 1.438% | 180 | 1.700% |
| 3 | Serpin peptidase inhibitor, clade A (alpha-1 antiproteinase, antitrypsin), member 3 OS=Bos taurus OX=9913 GN=SERPINA3-1 PE=3 SV=1 | A0A452DJK6 (+1) | 52 kDa | 5.51 | Protease Inhibitor | 90 | 1.043% | 91 | 1.011% | 94 | 0.987% | 90 | 1.047% | 86 | 1.130% | 112 | 1.088% | 224 | 2.115% |
| 4 | Cluster of Serpin A3-3 OS=Bos taurus OX=9913 GN=SERPINA3-3 PE=1 SV=2 (Q3ZEJ6) | Q3ZEJ6 [3] | 46 kDa | 5.78 | Protease Inhibitor | 98 | 1.136% | 97 | 1.077% | 94 | 0.987% | 109 | 1.268% | 116 | 1.525% | 122 | 1.185% | 194 | 1.832% |
| 5 | Alpha-1-acid glycoprotein OS=Bos taurus OX=9913 GN=ORM1 PE=2 SV=1 | Q3SZR3 (+1) | 23 kDa | 5.67 | Transport/Carrier/Binding | 64 | 0.742% | 62 | 0.689% | 50 | 0.525% | 55 | 0.640% | 55 | 0.723% | 61 | 0.593% | 111 | 1.048% |
| 6 | Alpha-1-antiproteinase OS=Bos taurus OX=9913 GN=SERPINA1 PE=1 SV=1 | P34955 | 46 kDa | 5.98 | Protease Inhibitor | 65 | 0.753% | 69 | 0.766% | 94 | 0.987% | 93 | 1.082% | 111 | 1.459% | 112 | 1.088% | 140 | 1.322% |
| 7 | Cluster of Serotransferrin OS=Bos taurus OX=9913 GN=TF PE=2 SV=1 (Q29443) | Q29443 [3] | 78 kDa | 6.5 | Transport/Carrier/Binding | 598 | 6.931% | 660 | 7.331% | 578 | 6.068% | 588 | 6.843% | 441 | 5.797% | 572 | 5.557% | 678 | 6.402% |
| 8 | Serpin domain-containing protein OS=Bos taurus OX=9913 GN=LOC784932 PE=1 SV=1 | A0A0A0MPA0 | 47 kDa | 5.44 | Protease Inhibitor | 25 | 0.290% | 29 | 0.322% | 29 | 0.304% | 44 | 0.512% | 47 | 0.618% | 42 | 0.408% | 47 | 0.444% |
| 9 | Cluster of Trypsin OS=Sus scrofa OX=9823 (P00761) | P00761 [2] | 24 kDa | 8.26 | Enzyme/Protease | 94 | 1.089% | 91 | 1.011% | 87 | 0.913% | 91 | 1.059% | 89 | 1.170% | 112 | 1.088% | 67 | 0.633% |
| 10 | Alpha-2-HS-glycoprotein OS=Bos taurus OX=9913 GN=AHSG PE=1 SV=2 | P12763 | 38 kDa | 5.1 | Protease Inhibitor | 25 | 0.290% | 25 | 0.278% | 38 | 0.399% | 35 | 0.407% | 35 | 0.460% | 39 | 0.379% | 72 | 0.680% |
| 11 | Serpin A3-8 OS=Bos taurus OX=9913 GN=SERPINA3-8 PE=2 SV=1 | A6QPQ2 | 47 kDa | 5.49 | Protease Inhibitor | 21 | 0.243% | 26 | 0.289% | 34 | 0.357% | 31 | 0.361% | 41 | 0.539% | 51 | 0.495% | 52 | 0.491% |
| 12 | Transthyretin OS=Bos taurus OX=9913 GN=TTR PE=1 SV=1 | O46375 | 16 kDa | 5.91 | Transport/Carrier/Binding | 23 | 0.267% | 27 | 0.300% | 31 | 0.325% | 39 | 0.454% | 35 | 0.460% | 31 | 0.301% | 71 | 0.670% |
| 13 | Corticosteroid-binding globulin OS=Bos taurus OX=9913 GN=SERPINA6 PE=3 SV=1 | E1BF81 | 45 kDa | 5.54 | Protease Inhibitor | 11 | 0.127% | 9 | 0.100% | 7 | 0.073% | 8 | 0.093% | 13 | 0.171% | 12 | 0.117% | 6 | 0.057% |
| 14 | Coagulation factor IX OS=Bos taurus OX=9913 GN=F9 PE=4 SV=3 | F1MBC5 (+1) | 50 kDa | 5.9 | Coagulation | 9 | 0.104% | 11 | 0.122% | 11 | 0.115% | 10 | 0.116% | 9 | 0.118% | 9 | 0.087% | 10 | 0.094% |
| 15 | Cluster of Complement C4 gamma chain OS=Bos taurus OX=9913 GN=LOC107131209 PE=4 SV=3 (F1MVK1) | F1MVK1 [4] | 192 kDa | 7.26 | Complement | 245 | 2.840% | 238 | 2.644% | 304 | 3.192% | 197 | 2.293% | 173 | 2.274% | 310 | 3.012% | 180 | 1.700% |
| 16 | Cluster of CD5 molecule like OS=Bos taurus OX=9913 GN=CD5L PE=2 SV=1 (A6QNW7) | A6QNW7 [3] | 50 kDa | 5.24 | Immunity/Immunity Related | 7 | 0.081% | 3 | 0.033% | 8 | 0.084% | 3 | 0.035% | 7 | 0.092% | 5 | 0.049% | 11 | 0.104% |
| 17 | Prothrombin OS=Bos taurus OX=9913 GN=F2 PE=1 SV=2 | P00735 | 71 kDa | 5.4 | Coagulation | 25 | 0.290% | 33 | 0.367% | 62 | 0.651% | 88 | 1.024% | 96 | 1.262% | 74 | 0.719% | 91 | 0.859% |
| 18 | Adiponectin B OS=Bos taurus OX=9913 GN=C1QC PE=4 SV=1 | A0A3B0IZF8 | 26 kDa | 8.62 | Complement | 6 | 0.070% | 4 | 0.044% | 6 | 0.063% | 5 | 0.058% | 5 | 0.066% | 4 | 0.039% | 4 | 0.038% |
| 19 | Adiponectin, C1Q and collagen domain containing OS=Bos taurus OX=9913 GN=ADIPOQ PE=4 SV=1 | A0A3Q1M564 | 35 kDa | 5.36 | Structural/Cytoskeleton | 21 | 0.243% | 18 | 0.200% | 20 | 0.210% | 19 | 0.221% | 19 | 0.250% | 20 | 0.194% | 6 | 0.057% |
| 20 | Afamin OS=Bos taurus OX=9913 GN=AFM PE=1 SV=1 | G3MYZ3 | 70 kDa | 5.57 | Transport/Carrier/Binding | 0 | 0.000% | 0 | 0.000% | 13 | 0.136% | 17 | 0.198% | 24 | 0.315% | 14 | 0.136% | 36 | 0.340% |
| 21 | Alpha-1-B glycoprotein OS=Bos taurus OX=9913 GN=A1BG PE=1 SV=1 | A0A3Q1MJT2 (+1) | 62 kDa | 5.37 | Immunity/Immunity Related | 22 | 0.255% | 19 | 0.211% | 20 | 0.210% | 26 | 0.303% | 40 | 0.526% | 24 | 0.233% | 61 | 0.576% |
| 22 | Alpha-2-antiplasmin OS=Bos taurus OX=9913 GN=SERPINF2 PE=1 SV=2 | P28800 | 55 kDa | 5.71 | Protease Inhibitor | 24 | 0.278% | 24 | 0.267% | 21 | 0.220% | 28 | 0.326% | 32 | 0.421% | 26 | 0.253% | 49 | 0.463% |
| 23 | Alpha-2-glycoprotein 1, zinc-binding OS=Bos taurus OX=9913 GN=AZGP1 PE=3 SV=1 | A0A452DK44 (+1) | 34 kDa | 5.11 | Immunity/Immunity Related | 0 | 0.000% | 0 | 0.000% | 2 | 0.021% | 4 | 0.047% | 4 | 0.053% | 0 | 0.000% | 21 | 0.198% |
| 24 | Angiotensinogen OS=Bos taurus OX=9913 GN=AGT PE=1 SV=2 | P01017 | 51 kDa | 6.49 | Protease Inhibitor | 53 | 0.614% | 61 | 0.678% | 53 | 0.556% | 64 | 0.745% | 48 | 0.631% | 44 | 0.427% | 38 | 0.359% |
| 25 | Apolipoprotein A4 OS=Bos taurus OX=9913 GN=APOA4 PE=3 SV=2 | F1N3Q7 | 52 kDa | 5.5 | Apolipoprotein | 12 | 0.139% | 19 | 0.211% | 13 | 0.136% | 27 | 0.314% | 18 | 0.237% | 7 | 0.068% | 51 | 0.482% |
| 26 | Apolipoprotein A-I OS=Bos taurus OX=9913 GN=APOA1 PE=1 SV=3 | P15497 | 30 kDa | 5.36 | Apolipoprotein | 125 | 1.449% | 123 | 1.366% | 121 | 1.270% | 137 | 1.594% | 114 | 1.498% | 129 | 1.253% | 126 | 1.190% |
| 27 | Apolipoprotein A-II OS=Bos taurus OX=9913 GN=APOA2 PE=1 SV=2 | P81644 | 11 kDa | 5.34 | Apolipoprotein | 0 | 0.000% | 2 | 0.022% | 0 | 0.000% | 4 | 0.047% | 2 | 0.026% | 0 | 0.000% | 0 | 0.000% |
| 28 | Apolipoprotein B OS=Bos taurus OX=9913 GN=APOB PE=1 SV=1 | A0A3Q1MFR4 (+1) | 514 kDa | 6.24 | Apolipoprotein | 8 | 0.093% | 3 | 0.033% | 12 | 0.126% | 4 | 0.047% | 7 | 0.092% | 6 | 0.058% | 0 | 0.000% |
| 29 | Apolipoprotein C-III OS=Bos taurus OX=9913 GN=APOC3 PE=1 SV=2 | P19035 | 11 kDa | 4.73 | Apolipoprotein | 0 | 0.000% | 0 | 0.000% | 0 | 0.000% | 0 | 0.000% | 0 | 0.000% | 0 | 0.000% | 5 | 0.047% |
| 30 | Apolipoprotein D OS=Bos taurus OX=9913 GN=APOD PE=3 SV=3 | F1MS32 | 24 kDa | 5.07 | Apolipoprotein | 4 | 0.046% | 7 | 0.078% | 4 | 0.042% | 4 | 0.047% | 3 | 0.039% | 0 | 0.000% | 6 | 0.057% |
| 31 | Apolipoprotein E OS=Bos taurus OX=9913 GN=APOE PE=1 SV=1 | Q03247 | 36 kDa | 5.55 | Apolipoprotein | 7 | 0.081% | 5 | 0.056% | 8 | 0.084% | 0 | 0.000% | 0 | 0.000% | 5 | 0.049% | 6 | 0.057% |
| 32 | ApoN protein OS=Bos taurus OX=9913 GN=APON PE=2 SV=1 | Q2KIH2 | 29 kDa | 5.36 | Apolipoprotein | 3 | 0.035% | 4 | 0.044% | 0 | 0.000% | 7 | 0.081% | 5 | 0.066% | 2 | 0.019% | 0 | 0.000% |
| 33 | Beta-2-glycoprotein 1 OS=Bos taurus OX=9913 GN=APOH PE=4 SV=1 | A0A140T843 (+1) | 38 kDa | 8.55 | Complement | 3 | 0.035% | 8 | 0.089% | 0 | 0.000% | 7 | 0.081% | 0 | 0.000% | 20 | 0.194% | 43 | 0.406% |
| 34 | Beta-2-microglobulin OS=Bos taurus OX=9913 PE=3 SV=1 | A0A3Q1MU93 (+1) | 14 kDa | 9.1 | Immunity/Immunity Related | 0 | 0.000% | 0 | 0.000% | 0 | 0.000% | 0 | 0.000% | 3 | 0.039% | 0 | 0.000% | 12 | 0.113% |
| 35 | Capping actin protein, gelsolin like OS=Bos taurus OX=9913 GN=CAPG PE=1 SV=1 | A0A3Q1N6D1 (+1) | 41 kDa | 6.03 | Transport/Carrier/Binding | 0 | 0.000% | 0 | 0.000% | 0 | 0.000% | 0 | 0.000% | 3 | 0.039% | 0 | 0.000% | 0 | 0.000% |
| 36 | Carbonic anhydrase 2 OS=Bos taurus OX=9913 GN=CA2 PE=1 SV=3 | P00921 | 29 kDa | 6.4 | Enzyme/Protease | 9 | 0.104% | 9 | 0.100% | 11 | 0.115% | 5 | 0.058% | 5 | 0.066% | 9 | 0.087% | 0 | 0.000% |
| 37 | Carboxypeptidase B2 OS=Bos taurus OX=9913 GN=CPB2 PE=3 SV=1 | A0A3Q1N1A7 (+1) | 45 kDa | 8.69 | Enzyme/Protease | 20 | 0.232% | 21 | 0.233% | 20 | 0.210% | 12 | 0.140% | 15 | 0.197% | 22 | 0.214% | 7 | 0.066% |
| 38 | Cartilage intermediate layer protein 2 OS=Bos taurus OX=9913 GN=CILP2 PE=4 SV=1 | E1BKL4 | 126 kDa | 8.7 | Cartilage Related | 0 | 0.000% | 0 | 0.000% | 3 | 0.031% | 0 | 0.000% | 0 | 0.000% | 2 | 0.019% | 4 | 0.038% |
| 39 | CD109 molecule OS=Bos taurus OX=9913 GN=CD109 PE=3 SV=3 | F1MPE1 | 161 kDa | 5.5 | Cell-Cell Adhesion & Signaling | 0 | 0.000% | 0 | 0.000% | 0 | 0.000% | 3 | 0.035% | 4 | 0.053% | 0 | 0.000% | 0 | 0.000% |
| 40 | chitinase OS=Bos taurus OX=9913 GN=CHIA PE=3 SV=2 | F1MH27 (+1) | 52 kDa | 5.36 | Enzyme/Protease | 3 | 0.035% | 0 | 0.000% | 0 | 0.000% | 3 | 0.035% | 4 | 0.053% | 0 | 0.000% | 0 | 0.000% |
| 41 | Chondroadherin OS=Bos taurus OX=9913 GN=CHAD PE=4 SV=2 | F1MYE4 | 41 kDa | 9.43 | Cartilage Related | 6 | 0.070% | 5 | 0.056% | 11 | 0.115% | 5 | 0.058% | 7 | 0.092% | 11 | 0.107% | 11 | 0.104% |
| 42 | Cluster of Actin, cytoplasmic 1 OS=Bos taurus OX=9913 GN=ACTB PE=1 SV=1 (P60712) | P60712 [7] | 42 kDa | 5.29 | Structural/Cytoskeleton | 52 | 0.603% | 42 | 0.467% | 44 | 0.462% | 53 | 0.617% | 39 | 0.513% | 54 | 0.525% | 33 | 0.312% |
| 43 | Cluster of Aggrecan OS=Bos taurus OX=9913 GN=ACAN PE=3 SV=2 (F1N367) | F1N367 [3] | 243 kDa | 4.17 | Extracellular Matrix | 39 | 0.452% | 32 | 0.355% | 53 | 0.556% | 43 | 0.500% | 26 | 0.342% | 82 | 0.797% | 56 | 0.529% |
| 44 | Cluster of Albumin OS=Oryctolagus cuniculus OX=9986 GN=ALB (P49065) | P49065 [2] | 69 kDa | 5.77 | Transport/Carrier/Binding | 0 | 0.000% | 68 | 0.755% | 69 | 0.724% | 66 | 0.768% | 0 | 0.000% | 66 | 0.641% | 138 | 1.303% |
| 45 | Cluster of Alpha-2-macroglobulin OS=Bos taurus OX=9913 GN=A2M PE=1 SV=2 (Q7SIH1) | Q7SIH1 [2] | 168 kDa | 5.68 | Protease Inhibitor | 312 | 3.616% | 296 | 3.288% | 442 | 4.640% | 231 | 2.688% | 170 | 2.234% | 483 | 4.693% | 338 | 3.191% |

| | | | | | | | | | | | | | | | | | | |
|---|---|---|---|---|---|---|---|---|---|---|---|---|---|---|---|---|---|---|
| 46 | Cluster of Alpha-amylase OS=Bos taurus OX=9913 GN=AMY2B PE=3 SV=1 (F1MJQ3) | F1MJQ3 [2] | 57 kDa | 5.87 | Enzyme/Protease | 0 | 0.000% | 0 | 0.000% | 0 | 0.000% | 0 | 0.000% | 0 | 0.000% | 0 | 0.000% | 18 | 0.170% |
| 47 | Cluster of Antithrombin-III OS=Bos taurus OX=9913 GN=SERPINC1 PE=3 SV=1 (A0A3Q1NJR8) | A0A3Q1NJR8 [2] | 60 kDa | 8.85 | Coagulation | 42 | 0.487% | 51 | 0.566% | 32 | 0.336% | 40 | 0.465% | 49 | 0.644% | 44 | 0.427% | 112 | 1.058% |
| 48 | Cluster of C4b-binding protein alpha chain OS=Bos taurus OX=9913 GN=C4BPA PE=2 SV=1 (Q28065) | Q28065 [3] | 69 kDa | 5.66 | Complement | 0 | 0.000% | 0 | 0.000% | 4 | 0.042% | 12 | 0.140% | 4 | 0.053% | 33 | 0.321% | 31 | 0.293% |
| 49 | Cluster of Cartilage oligomeric matrix protein OS=Bos taurus OX=9913 GN=COMP PE=1 SV=2 (P35445) | P35445 [2] | 82 kDa | 4.36 | Cartilage Related | 41 | 0.475% | 41 | 0.455% | 57 | 0.598% | 35 | 0.407% | 25 | 0.329% | 71 | 0.690% | 67 | 0.633% |
| 50 | Cluster of Cathelicidin-1 OS=Bos taurus OX=9913 GN=CATHL1 PE=1 SV=2 (P22226) | P22226 [6] | 18 kDa | 11.53 | Immunity/Immunity Related | 15 | 0.174% | 10 | 0.111% | 14 | 0.147% | 15 | 0.175% | 13 | 0.171% | 14 | 0.136% | 10 | 0.094% |
| 51 | Cluster of Ceruloplasmin OS=Bos taurus OX=9913 GN=CP PE=1 SV=1 (A0A3Q1NJB1) | A0A3Q1NJB1 [3] | 121 kDa | 5.67 | Transport/Carrier/Binding | 158 | 1.831% | 171 | 1.899% | 141 | 1.480% | 176 | 2.048% | 125 | 1.643% | 152 | 1.477% | 145 | 1.369% |
| 52 | Cluster of Clusterin OS=Bos taurus OX=9913 GN=CLU PE=1 SV=1 (P17697) | P17697 [2] | 51 kDa | 5.72 | Apolipoprotein | 80 | 0.927% | 72 | 0.800% | 70 | 0.735% | 56 | 0.652% | 59 | 0.775% | 81 | 0.787% | 66 | 0.623% |
| 53 | Cluster of Complement C2 OS=Bos taurus OX=9913 GN=C2 PE=4 SV=1 (A0A3S5ZPC6) | A0A3S5ZPC6 [3] | 87 kDa | 8.86 | Complement | 28 | 0.325% | 38 | 0.422% | 36 | 0.378% | 30 | 0.349% | 30 | 0.394% | 36 | 0.350% | 11 | 0.104% |
| 54 | Cluster of Complement C5 OS=Bos taurus OX=9913 GN=C5 PE=1 SV=3 (F1MY85) | F1MY85 [2] | 189 kDa | 6.16 | Complement | 91 | 1.055% | 139 | 1.544% | 175 | 1.837% | 58 | 0.675% | 48 | 0.631% | 184 | 1.788% | 77 | 0.727% |
| 55 | Cluster of Complement C8 gamma chain OS=Bos taurus OX=9913 GN=C8G PE=4 SV=1 (A0A3Q1MR54) | A0A3Q1MR54 [3] | 31 kDa | 10.67 | Complement | 9 | 0.104% | 12 | 0.133% | 16 | 0.168% | 10 | 0.116% | 6 | 0.079% | 20 | 0.194% | 6 | 0.057% |
| 56 | Cluster of Complement component C6 OS=Bos taurus OX=9913 GN=C6 PE=2 SV=1 (Q29RU4) | Q29RU4 [3] | 105 kDa | 6.59 | Complement | 11 | 0.127% | 8 | 0.089% | 16 | 0.168% | 22 | 0.256% | 10 | 0.131% | 26 | 0.253% | 33 | 0.312% |
| 57 | Cluster of Complement component C7 OS=Bos taurus OX=9913 GN=C7 PE=2 SV=1 (Q29RQ1) | Q29RQ1 [3] | 93 kDa | 6.9 | Complement | 15 | 0.174% | 15 | 0.167% | 16 | 0.168% | 15 | 0.175% | 16 | 0.210% | 29 | 0.282% | 25 | 0.236% |
| 58 | Cluster of Complement component C9 OS=Bos taurus OX=9913 GN=C9 PE=2 SV=1 (Q3MHN2) | Q3MHN2 [3] | 62 kDa | 5.57 | Complement | 78 | 0.904% | 80 | 0.889% | 66 | 0.693% | 72 | 0.838% | 72 | 0.946% | 61 | 0.593% | 44 | 0.415% |
| 59 | Cluster of Complement factor H OS=Bos taurus OX=9913 GN=CFH PE=1 SV=3 (Q28085) | Q28085 [6] | 140 kDa | 6.33 | Complement | 25 | 0.290% | 19 | 0.211% | 34 | 0.357% | 58 | 0.675% | 21 | 0.276% | 92 | 0.894% | 88 | 0.831% |
| 60 | Cluster of Complement factor I OS=Bos taurus OX=9913 GN=CFI PE=1 SV=1 (A0A3Q1MF14) | A0A3Q1MF14 [2] | 69 kDa | 8.13 | Complement | 35 | 0.406% | 36 | 0.400% | 36 | 0.378% | 29 | 0.337% | 23 | 0.302% | 41 | 0.398% | 39 | 0.368% |
| 61 | Cluster of Conglutinin OS=Bos taurus OX=9913 GN=CGN1 PE=1 SV=2 (P23805) | P23805 [3] | 38 kDa | 5.63 | Extracellular Matrix | 4 | 0.046% | 4 | 0.044% | 8 | 0.084% | 4 | 0.047% | 5 | 0.066% | 9 | 0.087% | 12 | 0.113% |
| 62 | Cluster of Elongation factor 1-alpha OS=Bos taurus OX=9913 PE=3 SV=3 (E1B9F6) | E1B9F6 [3] | 48 kDa | 9.1 | Other | 0 | 0.000% | 0 | 0.000% | 0 | 0.000% | 6 | 0.070% | 0 | 0.000% | 0 | 0.000% | 0 | 0.000% |
| 63 | Cluster of Fibrinogen beta chain OS=Bos taurus OX=9913 GN=FGB PE=4 SV=1 (A0A3Q1MG04) | A0A3Q1MG04 [3] | 57 kDa | 8.33 | Coagulation | 68 | 0.788% | 42 | 0.467% | 46 | 0.483% | 41 | 0.477% | 28 | 0.368% | 43 | 0.418% | 55 | 0.519% |
| 64 | Cluster of Fibronectin OS=Bos taurus OX=9913 GN=FN1 PE=1 SV=4 (P07589) | P07589 [3] | 272 kDa | 5.28 | Structural/Cytoskeleton | 260 | 3.013% | 244 | 2.710% | 285 | 2.992% | 223 | 2.595% | 177 | 2.326% | 343 | 3.332% | 278 | 2.625% |
| 65 | Cluster of Fructose-bisphosphate aldolase OS=Bos taurus OX=9913 GN=ALDOA PE=1 SV=1 (A0A3Q1LMG1) | A0A3Q1LMG1 [5] | 40 kDa | 8.6 | Enzyme/Protease | 0 | 0.000% | 3 | 0.033% | 4 | 0.042% | 5 | 0.058% | 0 | 0.000% | 7 | 0.068% | 2 | 0.019% |
| 66 | Cluster of Gelsolin OS=Bos taurus OX=9913 GN=GSN PE=1 SV=2 (F1N1I6) | F1N1I6 [2] | 92 kDa | 6.42 | Transport/Carrier/Binding | 162 | 1.878% | 156 | 1.733% | 147 | 1.543% | 159 | 1.850% | 140 | 1.840% | 155 | 1.506% | 144 | 1.360% |
| 67 | Cluster of Glyceraldehyde-3-phosphate dehydrogenase OS=Bos taurus OX=9913 GN=GAPDH PE=1 SV=4 (P10096) | P10096 [2] | 36 kDa | 8.52 | Enzyme/Protease | 4 | 0.046% | 0 | 0.000% | 2 | 0.021% | 9 | 0.105% | 3 | 0.039% | 0 | 0.000% | 0 | 0.000% |
| 68 | Cluster of Hemoglobin subunit alpha OS=Bos taurus OX=9913 GN=HBA PE=1 SV=2 (P01966) | P01966 [2] | 15 kDa | 8.19 | Transport/Carrier/Binding | 29 | 0.336% | 29 | 0.322% | 28 | 0.294% | 35 | 0.407% | 30 | 0.394% | 28 | 0.272% | 51 | 0.482% |
| 69 | Cluster of Hemoglobin subunit beta OS=Bos taurus OX=9913 GN=HBB PE=1 SV=1 (P02070) | P02070 [5] | 16 kDa | 7.02 | Transport/Carrier/Binding | 65 | 0.753% | 59 | 0.655% | 59 | 0.619% | 65 | 0.756% | 63 | 0.828% | 59 | 0.573% | 51 | 0.482% |
| 70 | Cluster of Histidine rich glycoprotein OS=Bos taurus OX=9913 GN=HRG PE=4 SV=3 (F1MKS5) | F1MKS5 [2] | 61 kDa | 7.14 | Protease Inhibitor | 39 | 0.452% | 40 | 0.444% | 38 | 0.399% | 31 | 0.361% | 27 | 0.355% | 38 | 0.369% | 60 | 0.567% |
| 71 | Cluster of Ig-like domain-containing protein OS=Bos taurus OX=9913 GN=LOC104968484 PE=4 SV=1 (A0A3Q1MFI7) | A0A3Q1MFI7 [9] | 24 kDa | 7.69 | Immunoglobulin | 14 | 0.162% | 22 | 0.244% | 20 | 0.210% | 14 | 0.163% | 17 | 0.223% | 34 | 0.330% | 12 | 0.113% |
| 72 | Cluster of Ig-like domain-containing protein OS=Bos taurus OX=9913 PE=1 SV=1 (A0A3Q1LUE9) | A0A3Q1LUE9 [5] | 17 kDa | 8.76 | Immunoglobulin | 28 | 0.325% | 34 | 0.378% | 30 | 0.315% | 26 | 0.303% | 24 | 0.315% | 36 | 0.350% | 22 | 0.208% |
| 73 | Cluster of Ig-like domain-containing protein OS=Bos taurus OX=9913 PE=1 SV=2 (G5E5T5) | G5E5T5 [4] | 56 kDa | 5.87 | Immunoglobulin | 43 | 0.498% | 44 | 0.489% | 53 | 0.556% | 58 | 0.675% | 46 | 0.605% | 64 | 0.622% | 87 | 0.821% |
| 74 | Cluster of Ig-like domain-containing protein OS=Bos taurus OX=9913 PE=4 SV=1 (A0A3Q1LSF0) | A0A3Q1LSF0 [3] | 15 kDa | 9.82 | Immunoglobulin | 17 | 0.197% | 18 | 0.200% | 31 | 0.325% | 19 | 0.221% | 14 | 0.184% | 26 | 0.253% | 17 | 0.161% |
| 75 | Cluster of Ig-like domain-containing protein OS=Bos taurus OX=9913 PE=4 SV=1 (A0A3Q1MBT3) | A0A3Q1MBT3 [5] | 12 kDa | 6.52 | Immunoglobulin | 9 | 0.104% | 4 | 0.044% | 10 | 0.105% | 9 | 0.105% | 0 | 0.000% | 7 | 0.068% | 10 | 0.094% |
| 76 | Cluster of Ig-like domain-containing protein OS=Bos taurus OX=9913 PE=4 SV=3 (F1N160) | F1N160 [4] | 34 kDa | 8.86 | Immunoglobulin | 164 | 1.901% | 212 | 2.355% | 213 | 2.236% | 146 | 1.699% | 134 | 1.761% | 191 | 1.856% | 294 | 2.776% |
| 77 | Cluster of Inter-alpha-trypsin inhibitor heavy chain 4 OS=Bos taurus OX=9913 GN=ITIH4 PE=3 SV=3 (F1MMD7) | F1MMD7 [3] | 102 kDa | 5.99 | Immunity/Immunity Related | 115 | 1.333% | 111 | 1.233% | 116 | 1.218% | 117 | 1.362% | 112 | 1.472% | 107 | 1.040% | 125 | 1.180% |
| 78 | Cluster of Junction plakoglobin OS=Bos taurus OX=9913 GN=JUP PE=1 SV=1 (A0A3Q1M1M7) | A0A3Q1M1M7 [4] | 81 kDa | 5.95 | Transport/Carrier/Binding | 16 | 0.185% | 10 | 0.111% | 22 | 0.231% | 48 | 0.559% | 19 | 0.250% | 0 | 0.000% | 0 | 0.000% |
| 79 | Cluster of Kininogen 1 OS=Bos taurus OX=9913 GN=KNG1 PE=4 SV=2 (F1MNV5) | F1MNV5 [2] | 48 kDa | 5.69 | Coagulation | 33 | 0.382% | 36 | 0.400% | 48 | 0.504% | 61 | 0.710% | 66 | 0.868% | 63 | 0.612% | 75 | 0.708% |
| 80 | Cluster of L-lactate dehydrogenase OS=Bos taurus OX=9913 GN=LDHB PE=1 SV=1 (A0A3Q1M5R4) | A0A3Q1M5R4 [6] | 37 kDa | 5.86 | Enzyme/Protease | 8 | 0.093% | 10 | 0.111% | 6 | 0.063% | 6 | 0.070% | 6 | 0.079% | 4 | 0.039% | 0 | 0.000% |
| 81 | Cluster of Maltase-glucoamylase OS=Bos taurus OX=9913 GN=MGAM PE=3 SV=2 (G3N3S2) | G3N3S2 [2] | 308 kDa | 5.4 | Enzyme/Protease | 0 | 0.000% | 0 | 0.000% | 0 | 0.000% | 2 | 0.023% | 0 | 0.000% | 0 | 0.000% | 0 | 0.000% |
| 82 | Cluster of Moesin OS=Bos taurus OX=9913 GN=MSN PE=2 SV=3 (Q2HJ49) | Q2HJ49 [4] | 68 kDa | 5.9 | Structural/Cytoskeleton | 2 | 0.023% | 5 | 0.056% | 2 | 0.021% | 0 | 0.000% | 0 | 0.000% | 2 | 0.019% | 0 | 0.000% |
| 83 | Cluster of Myocilin OS=Bos taurus OX=9913 GN=MYOC PE=2 SV=1 (Q3SX06) | Q3SX06 [2] | 55 kDa | 5.33 | Structural/Cytoskeleton | 3 | 0.035% | 2 | 0.022% | 0 | 0.000% | 3 | 0.035% | 4 | 0.053% | 0 | 0.000% | 0 | 0.000% |
| 84 | Cluster of Peptidoglycan recognition protein 1 OS=Bos taurus OX=9913 GN=PGLYRP1 PE=1 SV=1 (Q8SPP7) | Q8SPP7 [2] | 21 kDa | 9.38 | Immunity/Immunity Related | 0 | 0.000% | 2 | 0.022% | 0 | 0.000% | 0 | 0.000% | 0 | 0.000% | 0 | 0.000% | 9 | 0.085% |
| 85 | Cluster of Plasma kallikrein OS=Bos taurus OX=9913 GN=KLKB1 PE=2 SV=1 (Q2KJ63) | Q2KJ63 [2] | 71 kDa | 8.62 | Complement | 4 | 0.046% | 4 | 0.044% | 0 | 0.000% | 3 | 0.035% | 4 | 0.053% | 0 | 0.000% | 0 | 0.000% |
| 86 | Cluster of Plasminogen OS=Bos taurus OX=9913 GN=PLG PE=3 SV=2 (E1B726) | E1B726 [2] | 91 kDa | 7.71 | Coagulation | 52 | 0.603% | 67 | 0.744% | 90 | 0.945% | 85 | 0.989% | 47 | 0.618% | 139 | 1.350% | 124 | 1.171% |
| 87 | Cluster of Pregnancy zone protein OS=Bos taurus OX=9913 GN=LOC506828 PE=3 SV=3 (F1MJK3) | F1MJK3 [6] | 163 kDa | 6.91 | Protease Inhibitor | 150 | 1.739% | 141 | 1.566% | 150 | 1.575% | 95 | 1.106% | 83 | 1.091% | 176 | 1.710% | 40 | 0.378% |
| 88 | Cluster of Primary amine oxidase, liver isozyme OS=Bos taurus OX=9913 PE=1 SV=1 (Q29437) | Q29437 [6] | 85 kDa | 5.58 | Enzyme/Protease | 54 | 0.626% | 62 | 0.689% | 56 | 0.588% | 59 | 0.687% | 57 | 0.749% | 58 | 0.563% | 60 | 0.567% |
| 89 | Cluster of Pyruvate kinase OS=Bos taurus OX=9913 GN=PKM PE=1 SV=1 (A5D984) | A5D984 [2] | 58 kDa | 7.96 | Enzyme/Protease | 17 | 0.197% | 9 | 0.100% | 12 | 0.126% | 15 | 0.175% | 3 | 0.039% | 12 | 0.117% | 0 | 0.000% |
| 90 | Cluster of Serpin family G member 1 OS=Bos taurus OX=9913 GN=SERPING1 PE=3 SV=3 (E1BMJ0) | E1BMJ0 [2] | 52 kDa | 6.28 | Complement | 39 | 0.452% | 38 | 0.422% | 40 | 0.420% | 34 | 0.396% | 27 | 0.355% | 43 | 0.418% | 24 | 0.227% |
| 91 | Cluster of SERPINA10 protein OS=Bos taurus OX=9913 GN=SERPINA10 PE=2 SV=1 (A5PJ69) | A5PJ69 [2] | 52 kDa | 6.05 | Protease Inhibitor | 0 | 0.000% | 0 | 0.000% | 0 | 0.000% | 0 | 0.000% | 0 | 0.000% | 0 | 0.000% | 12 | 0.113% |
| 92 | Cluster of Serum amyloid A protein OS=Bos taurus OX=9913 GN=LOC104968478 PE=3 SV=2 (E1BJF9) | E1BJF9 [4] | 20 kDa | 6.8 | Apolipoprotein | 4 | 0.046% | 5 | 0.056% | 0 | 0.000% | 8 | 0.093% | 5 | 0.066% | 3 | 0.029% | 11 | 0.104% |
| 93 | Cluster of Vinculin OS=Bos taurus OX=9913 GN=VCL PE=1 SV=1 (F1N789) | F1N789 [2] | 124 kDa | 5.58 | Structural/Cytoskeleton | 9 | 0.104% | 10 | 0.111% | 13 | 0.136% | 6 | 0.070% | 10 | 0.131% | 19 | 0.185% | 0 | 0.000% |
| 94 | Cluster of Vitamin D-binding protein OS=Bos taurus OX=9913 GN=GC PE=4 SV=2 (F1N5M2) | F1N5M2 [3] | 53 kDa | 5.24 | Transport/Carrier/Binding | 28 | 0.325% | 28 | 0.311% | 37 | 0.388% | 51 | 0.594% | 72 | 0.946% | 48 | 0.466% | 124 | 1.171% |
| 95 | Coagulation factor X OS=Bos taurus OX=9913 GN=F10 PE=4 SV=1 | A0A452DIP8 | 67 kDa | 5.96 | Coagulation | 0 | 0.000% | 4 | 0.044% | 0 | 0.000% | 0 | 0.000% | 0 | 0.000% | 2 | 0.019% | 0 | 0.000% |

| | | | | | | | | | | | | | | | | | | | |
|---|---|---|---|---|---|---|---|---|---|---|---|---|---|---|---|---|---|---|---|
| 96 | Coagulation factor XIII A chain OS=Bos taurus OX=9913 GN=F13A1 PE=3 SV=1 | A0A3Q1LTB9 (+1) | 76 kDa | 6 | Coagulation | 7 | 0.081% | 0 | 0.000% | 4 | 0.042% | 0 | 0.000% | 0 | 0.000% | 4 | 0.039% | 0 | 0.000% |
| 97 | Coagulation factor XIII B chain OS=Bos taurus OX=9913 GN=F13B PE=4 SV=1 | A0A3Q1MWQ1 (+1) | 68 kDa | 6 | Coagulation | 0 | 0.000% | 0 | 0.000% | 0 | 0.000% | 0 | 0.000% | 0 | 0.000% | 3 | 0.029% | 0 | 0.000% |
| 98 | Collagen type VI alpha 1 chain OS=Bos taurus OX=9913 GN=COL6A1 PE=1 SV=1 | E1BI98 | 109 kDa | 5.18 | Extracellular Matrix | 9 | 0.104% | 6 | 0.067% | 5 | 0.052% | 7 | 0.081% | 10 | 0.131% | 7 | 0.068% | 0 | 0.000% |
| 99 | Collagen type VI alpha 3 chain OS=Bos taurus OX=9913 GN=COL6A3 PE=1 SV=2 | E1BB91 | 340 kDa | 5.87 | Extracellular Matrix | 19 | 0.220% | 14 | 0.156% | 14 | 0.147% | 11 | 0.128% | 18 | 0.237% | 22 | 0.214% | 3 | 0.028% |
| 100 | Complement C1q subcomponent subunit A OS=Bos taurus OX=9913 GN=C1QA PE=2 SV=1 | Q5E9E3 | 26 kDa | 9.12 | Complement | 3 | 0.035% | 0 | 0.000% | 4 | 0.042% | 4 | 0.047% | 0 | 0.000% | 6 | 0.058% | 0 | 0.000% |
| 101 | Complement C1q subcomponent subunit B OS=Bos taurus OX=9913 GN=C1QB PE=1 SV=1 | Q2KIV9 | 26 kDa | 9.26 | Complement | 2 | 0.023% | 0 | 0.000% | 2 | 0.021% | 0 | 0.000% | 0 | 0.000% | 5 | 0.049% | 3 | 0.028% |
| 102 | Complement C1r OS=Bos taurus OX=9913 GN=C1R PE=4 SV=1 | A0A3Q1MGP1 (+1) | 81 kDa | 5.8 | Complement | 7 | 0.081% | 4 | 0.044% | 5 | 0.052% | 2 | 0.023% | 2 | 0.026% | 7 | 0.068% | 4 | 0.038% |
| 103 | Complement C1s OS=Bos taurus OX=9913 GN=C1S PE=4 SV=1 | F1MJ12 (+1) | 77 kDa | 4.97 | Complement | 22 | 0.255% | 22 | 0.244% | 20 | 0.210% | 23 | 0.268% | 15 | 0.197% | 19 | 0.185% | 9 | 0.085% |
| 104 | Complement C3 OS=Bos taurus OX=9913 GN=C3 PE=1 SV=2 | Q2UVX4 | 187 kDa | 6.37 | Complement | 698 | 8.090% | 761 | 8.453% | 827 | 8.682% | 621 | 7.227% | 554 | 7.282% | 886 | 8.608% | 501 | 4.730% |
| 105 | Complement C3 OS=Bos taurus OX=9913 GN=LOC528040 PE=4 SV=3 | E1B805 | 186 kDa | 6.54 | Complement | 138 | 1.599% | 129 | 1.433% | 141 | 1.480% | 111 | 1.292% | 100 | 1.314% | 154 | 1.496% | 7 | 0.066% |
| 106 | Complement C3d receptor 2 OS=Bos taurus OX=9913 GN=CR2 PE=4 SV=1 | A0A3Q1M7W3 | 163 kDa | 7.13 | Complement | 0 | 0.000% | 0 | 0.000% | 0 | 0.000% | 0 | 0.000% | 0 | 0.000% | 4 | 0.039% | 0 | 0.000% |
| 107 | Complement C8 alpha chain OS=Bos taurus OX=9913 GN=C8A PE=3 SV=1 | A0A3Q1LSP4 (+1) | 65 kDa | 6.24 | Complement | 19 | 0.220% | 20 | 0.222% | 31 | 0.325% | 18 | 0.209% | 13 | 0.171% | 36 | 0.350% | 16 | 0.151% |
| 108 | Complement C8 beta chain OS=Bos taurus OX=9913 GN=C8B PE=3 SV=3 | F1N102 | 67 kDa | 8.15 | Complement | 11 | 0.127% | 15 | 0.167% | 24 | 0.252% | 13 | 0.151% | 8 | 0.105% | 31 | 0.301% | 24 | 0.227% |
| 109 | Complement factor B OS=Bos taurus OX=9913 GN=CFB PE=1 SV=2 | P81187 | 85 kDa | 7.68 | Complement | 90 | 1.043% | 105 | 1.166% | 108 | 1.134% | 89 | 1.036% | 72 | 0.946% | 131 | 1.273% | 106 | 1.001% |
| 110 | Complement factor D OS=Bos taurus OX=9913 GN=CFD PE=2 SV=1 | Q3T0A3 | 28 kDa | 6.85 | Complement | 2 | 0.023% | 8 | 0.089% | 8 | 0.084% | 0 | 0.000% | 0 | 0.000% | 8 | 0.078% | 7 | 0.066% |
| 111 | Complement factor properdin OS=Bos taurus OX=9913 GN=CFP PE=4 SV=1 | A0A3Q1M2R9 (+1) | 63 kDa | 8.33 | Complement | 0 | 0.000% | 0 | 0.000% | 0 | 0.000% | 0 | 0.000% | 0 | 0.000% | 0 | 0.000% | 3 | 0.028% |
| 112 | C-type lectin domain family 3 member B OS=Bos taurus OX=9913 GN=CLEC3B PE=4 SV=1 | A0A452DIU4 | 31 kDa | 6.52 | Cell-Cell Adhesion & Signaling | 5 | 0.058% | 7 | 0.078% | 4 | 0.042% | 8 | 0.093% | 5 | 0.066% | 0 | 0.000% | 17 | 0.161% |
| 113 | Cystatin-C OS=Bos taurus OX=9913 GN=CST3 PE=1 SV=2 | P01035 | 16 kDa | 9.03 | Protease Inhibitor | 0 | 0.000% | 0 | 0.000% | 0 | 0.000% | 0 | 0.000% | 0 | 0.000% | 0 | 0.000% | 16 | 0.151% |
| 114 | Desmoglein 1 OS=Bos taurus OX=9913 GN=DSG1 PE=4 SV=2 | F1MIW8 (+1) | 112 kDa | 4.88 | Cell-Cell Adhesion & Signaling | 0 | 0.000% | 0 | 0.000% | 5 | 0.052% | 11 | 0.128% | 0 | 0.000% | 0 | 0.000% | 0 | 0.000% |
| 115 | EGF containing fibulin extracellular matrix protein 1 OS=Bos taurus OX=9913 GN=EFEMP1 PE=1 SV=1 | A2VE41 | 55 kDa | 4.83 | Extracellular Matrix | 0 | 0.000% | 3 | 0.033% | 0 | 0.000% | 0 | 0.000% | 0 | 0.000% | 0 | 0.000% | 0 | 0.000% |
| 116 | Extracellular matrix protein 1 OS=Bos taurus OX=9913 GN=ECM1 PE=4 SV=1 | A0A3Q1M5Q6 (+1) | 63 kDa | 6.6 | Extracellular Matrix | 5 | 0.058% | 7 | 0.078% | 5 | 0.052% | 5 | 0.058% | 4 | 0.053% | 13 | 0.126% | 21 | 0.198% |
| 117 | Fatty acid-binding protein 5 OS=Bos taurus OX=9913 GN=FABP5 PE=1 SV=4 | P55052 | 15 kDa | 7.8 | Transport/Carrier/Binding | 0 | 0.000% | 0 | 0.000% | 0 | 0.000% | 3 | 0.035% | 0 | 0.000% | 0 | 0.000% | 0 | 0.000% |
| 118 | Fetuin-B OS=Bos taurus OX=9913 GN=FETUB PE=1 SV=1 | Q58D62 | 43 kDa | 5.59 | Protease Inhibitor | 0 | 0.000% | 0 | 0.000% | 0 | 0.000% | 0 | 0.000% | 0 | 0.000% | 0 | 0.000% | 23 | 0.217% |
| 119 | Fibrinogen alpha chain OS=Bos taurus OX=9913 GN=FGA PE=4 SV=1 | F6QND5 | 95 kDa | 5.69 | Coagulation | 120 | 1.391% | 87 | 0.966% | 94 | 0.987% | 88 | 1.024% | 77 | 1.012% | 96 | 0.933% | 63 | 0.595% |
| 120 | Fibrinogen gamma chain OS=Bos taurus OX=9913 GN=FGG PE=4 SV=1 | F1MGU7 (+1) | 50 kDa | 5.38 | Coagulation | 68 | 0.788% | 59 | 0.655% | 62 | 0.651% | 41 | 0.477% | 43 | 0.565% | 60 | 0.583% | 42 | 0.397% |
| 121 | Fibromodulin OS=Bos taurus OX=9913 GN=FMOD PE=1 SV=2 | P13605 | 43 kDa | 5.57 | Extracellular Matrix | 0 | 0.000% | 0 | 0.000% | 0 | 0.000% | 0 | 0.000% | 0 | 0.000% | 0 | 0.000% | 3 | 0.028% |
| 122 | Fibulin-1 OS=Bos taurus OX=9913 GN=FBLN1 PE=1 SV=1 | A0A3Q1MG73 | 74 kDa | 4.9 | Coagulation | 61 | 0.707% | 60 | 0.666% | 61 | 0.640% | 41 | 0.477% | 38 | 0.499% | 57 | 0.554% | 19 | 0.179% |
| 123 | Glutathione peroxidase 3 OS=Bos taurus OX=9913 GN=GPX3 PE=2 SV=2 | P37141 | 26 kDa | 6 | Enzyme/Protease | 0 | 0.000% | 0 | 0.000% | 0 | 0.000% | 0 | 0.000% | 0 | 0.000% | 0 | 0.000% | 6 | 0.057% |
| 124 | Haptoglobin OS=Bos taurus OX=9913 GN=HP PE=3 SV=1 | A0A3Q1MB98 (+1) | 44 kDa | 8.09 | Complement | 12 | 0.139% | 8 | 0.089% | 12 | 0.126% | 17 | 0.198% | 12 | 0.158% | 25 | 0.243% | 16 | 0.151% |
| 125 | Heat shock protein beta-1 OS=Bos taurus OX=9913 GN=HSPB1 PE=2 SV=1 | Q3T149 | 22 kDa | 5.98 | Other | 0 | 0.000% | 0 | 0.000% | 0 | 0.000% | 4 | 0.047% | 0 | 0.000% | 0 | 0.000% | 0 | 0.000% |
| 126 | Hemopexin OS=Bos taurus OX=9913 GN=HPX PE=2 SV=1 | Q3SZV7 | 52 kDa | 7.1 | Transport/Carrier/Binding | 101 | 1.171% | 105 | 1.166% | 90 | 0.945% | 97 | 1.129% | 65 | 0.854% | 85 | 0.826% | 172 | 1.624% |
| 127 | HGF activator OS=Bos taurus OX=9913 GN=HGFAC PE=4 SV=3 | E1BCW0 | 70 kDa | 7.66 | Coagulation | 5 | 0.058% | 2 | 0.022% | 0 | 0.000% | 3 | 0.035% | 3 | 0.039% | 0 | 0.000% | 0 | 0.000% |
| 128 | Hyaluronan and proteoglycan link protein 1 OS=Bos taurus OX=9913 GN=HAPLN1 PE=2 SV=1 | P55252 | 40 kDa | 7.66 | Cartilage Related | 11 | 0.127% | 3 | 0.033% | 10 | 0.105% | 4 | 0.047% | 3 | 0.039% | 15 | 0.146% | 10 | 0.094% |
| 129 | Ig-like domain-containing protein OS=Bos taurus OX=9913 PE=1 SV=1 | A0A3Q1M3L6 | 40 kDa | 5.16 | Immunoglobulin | 272 | 3.153% | 303 | 3.366% | 309 | 3.244% | 254 | 2.956% | 189 | 2.484% | 334 | 3.245% | 392 | 3.701% |
| 130 | Ig-like domain-containing protein OS=Bos taurus OX=9913 PE=1 SV=1 | G3N0V0 | 36 kDa | 8.05 | Immunoglobulin | 186 | 2.156% | 197 | 2.188% | 210 | 2.205% | 153 | 1.781% | 163 | 2.142% | 204 | 1.982% | 358 | 3.380% |
| 131 | Ig-like domain-containing protein OS=Bos taurus OX=9913 PE=1 SV=1 | A0A3Q1LPG0 | 36 kDa | 7.59 | Immunoglobulin | 47 | 0.545% | 52 | 0.578% | 58 | 0.609% | 49 | 0.570% | 43 | 0.565% | 56 | 0.544% | 45 | 0.425% |
| 132 | Ig-like domain-containing protein OS=Bos taurus OX=9913 PE=1 SV=1 | A0A3Q1LRW4 (+1) | 47 kDa | 8.2 | Immunoglobulin | 22 | 0.255% | 28 | 0.311% | 26 | 0.273% | 19 | 0.221% | 15 | 0.197% | 28 | 0.272% | 21 | 0.198% |
| 133 | Ig-like domain-containing protein OS=Bos taurus OX=9913 PE=1 SV=3 | F1MZ96 | 27 kDa | 5.94 | Immunoglobulin | 70 | 0.811% | 74 | 0.822% | 75 | 0.787% | 76 | 0.884% | 50 | 0.657% | 84 | 0.816% | 60 | 0.567% |
| 134 | Ig-like domain-containing protein OS=Bos taurus OX=9913 PE=4 SV=1 | A0A3Q1LJT1 | 25 kDa | 4.73 | Immunoglobulin | 0 | 0.000% | 5 | 0.056% | 3 | 0.031% | 3 | 0.035% | 3 | 0.039% | 0 | 0.000% | 2 | 0.019% |
| 135 | Ig-like domain-containing protein OS=Bos taurus OX=9913 PE=4 SV=1 | A0A3Q1MR79 | 21 kDa | 10.28 | Immunoglobulin | 7 | 0.081% | 6 | 0.067% | 10 | 0.105% | 7 | 0.081% | 3 | 0.039% | 8 | 0.078% | 0 | 0.000% |
| 136 | Ig-like domain-containing protein OS=Bos taurus OX=9913 PE=4 SV=1 | A0A3Q1LLT0 (+1) | 24 kDa | 9.39 | Immunoglobulin | 0 | 0.000% | 0 | 0.000% | 0 | 0.000% | 0 | 0.000% | 0 | 0.000% | 6 | 0.058% | 0 | 0.000% |
| 137 | Ig-like domain-containing protein OS=Bos taurus OX=9913 PE=4 SV=2 | G3MXG6 | 15 kDa | 8.56 | Immunoglobulin | 0 | 0.000% | 9 | 0.100% | 11 | 0.115% | 3 | 0.035% | 3 | 0.039% | 5 | 0.049% | 0 | 0.000% |
| 138 | Ig-like domain-containing protein OS=Bos taurus OX=9913 PE=4 SV=2 | G3N342 | 55 kDa | 6.55 | Immunoglobulin | 0 | 0.000% | 0 | 0.000% | 2 | 0.021% | 0 | 0.000% | 0 | 0.000% | 3 | 0.029% | 0 | 0.000% |
| 139 | Immunoglobulin J chain OS=Bos taurus OX=9913 GN=JCHAIN PE=1 SV=1 | Q3SYR8 | 18 kDa | 4.92 | Immunoglobulin | 4 | 0.046% | 4 | 0.044% | 4 | 0.042% | 4 | 0.047% | 4 | 0.053% | 8 | 0.078% | 7 | 0.066% |
| 140 | Inhibitor of carbonic anhydrase OS=Bos taurus OX=9913 GN=INHCA PE=1 SV=1 | A0A3Q1NEQ0 | 78 kDa | 6.8 | Transport/Carrier/Binding | 49 | 0.568% | 57 | 0.633% | 54 | 0.567% | 39 | 0.454% | 29 | 0.381% | 57 | 0.554% | 16 | 0.151% |
| 141 | Insulin-like growth factor binding protein acid labile subunit OS=Bos taurus OX=9913 GN=IGFALS PE=2 SV=1 | Q09TE3 | 66 kDa | 6.73 | Structural/Cytoskeleton | 13 | 0.151% | 12 | 0.133% | 7 | 0.073% | 10 | 0.116% | 10 | 0.131% | 9 | 0.087% | 3 | 0.028% |
| 142 | Inter-alpha-trypsin inhibitor heavy chain 2 OS=Bos taurus OX=9913 GN=ITIH2 PE=1 SV=1 | A0A3Q1LK49 | 97 kDa | 8.72 | Transport/Carrier/Binding | 115 | 1.333% | 103 | 1.144% | 100 | 1.050% | 109 | 1.268% | 98 | 1.288% | 92 | 0.894% | 49 | 0.463% |
| 143 | Inter-alpha-trypsin inhibitor heavy chain H1 OS=Bos taurus OX=9913 GN=ITIH1 PE=1 SV=1 | Q0VCM5 | 101 kDa | 7.52 | Cartilage Related | 127 | 1.472% | 130 | 1.444% | 124 | 1.302% | 126 | 1.466% | 125 | 1.643% | 120 | 1.166% | 63 | 0.595% |
| 144 | LDL receptor related protein 1 OS=Bos taurus OX=9913 GN=LRP1 PE=1 SV=3 | E1BGJ0 | 505 kDa | 5.12 | Apolipoprotein | 0 | 0.000% | 0 | 0.000% | 0 | 0.000% | 0 | 0.000% | 0 | 0.000% | 5 | 0.049% | 0 | 0.000% |
| 145 | Leucine rich alpha-2-glycoprotein 1 OS=Bos taurus OX=9913 GN=LRG1 PE=1 SV=1 | F6RMV5 | 39 kDa | 6.26 | Structural/Cytoskeleton | 0 | 0.000% | 0 | 0.000% | 0 | 0.000% | 0 | 0.000% | 5 | 0.066% | 0 | 0.000% | 41 | 0.387% |

| | | | | | | | | | | | | | | | | | | | |
|---|---|---|---|---|---|---|---|---|---|---|---|---|---|---|---|---|---|---|---|
| 146 | Lipopolysaccharide-binding protein OS=Bos taurus OX=9913 GN=LBP PE=3 SV=2 | F1MNN7 (+1) | 54 kDa | 6.38 | Immunity/Immunity Related | 13 | 0.151% | 17 | 0.189% | 10 | 0.105% | 21 | 0.244% | 10 | 0.131% | 11 | 0.107% | 15 | 0.142% |
| 147 | Lumican OS=Bos taurus OX=9913 GN=LUM PE=1 SV=1 | Q05443 | 39 kDa | 5.93 | Extracellular Matrix | 20 | 0.232% | 19 | 0.211% | 19 | 0.199% | 24 | 0.279% | 21 | 0.276% | 20 | 0.194% | 24 | 0.227% |
| 148 | Lymphocyte cytosolic protein 1 OS=Bos taurus OX=9913 GN=LCP1 PE=1 SV=1 | A0A3Q1LSN0 (+1) | 70 kDa | 5.17 | Transport/Carrier/Binding | 0 | 0.000% | 0 | 0.000% | 0 | 0.000% | 0 | 0.000% | 0 | 0.000% | 0 | 0.000% | 3 | 0.028% |
| 149 | Mannose receptor C type 2 OS=Bos taurus OX=9913 GN=MRC2 PE=4 SV=1 | A0A3Q1M5H7 (+1) | 167 kDa | 5.62 | Transport/Carrier/Binding | 3 | 0.035% | 0 | 0.000% | 0 | 0.000% | 0 | 0.000% | 4 | 0.053% | 0 | 0.000% | 0 | 0.000% |
| 150 | Mannose-binding protein C OS=Bos taurus OX=9913 GN=MBL PE=2 SV=1 | O02659 | 26 kDa | 5.11 | Complement | 0 | 0.000% | 0 | 0.000% | 3 | 0.031% | 2 | 0.023% | 0 | 0.000% | 7 | 0.068% | 0 | 0.000% |
| 151 | Pantetheinase OS=Bos taurus OX=9913 GN=VNN1 PE=1 SV=1 | Q58CQ9 | 57 kDa | 5.26 | Enzyme/Protease | 0 | 0.000% | 0 | 0.000% | 0 | 0.000% | 2 | 0.023% | 3 | 0.039% | 3 | 0.029% | 8 | 0.076% |
| 152 | Paraoxonase OS=Bos taurus OX=9913 GN=PON1 PE=1 SV=1 | Q2KIW1 | 40 kDa | 5.17 | Enzyme/Protease | 15 | 0.174% | 15 | 0.167% | 13 | 0.136% | 15 | 0.175% | 8 | 0.105% | 13 | 0.126% | 14 | 0.132% |
| 153 | Pentraxin family member OS=Bos taurus OX=9913 GN=CRP PE=2 SV=1 | C4T8B4 | 25 kDa | 6.48 | Immunity/Immunity Related | 8 | 0.093% | 6 | 0.067% | 8 | 0.084% | 0 | 0.000% | 0 | 0.000% | 4 | 0.039% | 0 | 0.000% |
| 154 | Peptidase D OS=Bos taurus OX=9913 GN=PEPD PE=3 SV=1 | A0A3Q1NGC5 (+1) | 55 kDa | 5.53 | Enzyme/Protease | 0 | 0.000% | 0 | 0.000% | 0 | 0.000% | 2 | 0.023% | 0 | 0.000% | 0 | 0.000% | 0 | 0.000% |
| 155 | Peptidoglycan recognition protein 2 OS=Bos taurus OX=9913 GN=PGLYRP2 PE=3 SV=3 | E1BH94 | 63 kDa | 6.46 | Immunity/Immunity Related | 0 | 0.000% | 0 | 0.000% | 0 | 0.000% | 0 | 0.000% | 0 | 0.000% | 0 | 0.000% | 10 | 0.094% |
| 156 | Phosphatidylinositol-glycan-specific phospholipase D OS=Bos taurus OX=9913 GN=GPLD1 PE=3 SV=1 | A0A3Q1MEX6 | 91 kDa | 6.06 | Enzyme/Protease | 0 | 0.000% | 0 | 0.000% | 0 | 0.000% | 9 | 0.105% | 3 | 0.039% | 0 | 0.000% | 0 | 0.000% |
| 157 | Phospholipid transfer protein OS=Bos taurus OX=9913 GN=PLTP PE=2 SV=1 | Q58DL9 | 56 kDa | 6.32 | Immunity/Immunity Related | 22 | 0.255% | 23 | 0.255% | 19 | 0.199% | 27 | 0.314% | 22 | 0.289% | 17 | 0.165% | 11 | 0.104% |
| 158 | phosphopyruvate hydratase OS=Bos taurus OX=9913 GN=ENO1 PE=3 SV=2 | F1MB08 | 54 kDa | 9.2 | Enzyme/Protease | 0 | 0.000% | 0 | 0.000% | 0 | 0.000% | 4 | 0.047% | 0 | 0.000% | 0 | 0.000% | 0 | 0.000% |
| 159 | Pigment epithelium-derived factor OS=Bos taurus OX=9913 GN=SERPINF1 PE=1 SV=1 | Q95121 | 46 kDa | 6.31 | Coagulation | 71 | 0.823% | 70 | 0.778% | 85 | 0.892% | 74 | 0.861% | 62 | 0.815% | 89 | 0.865% | 40 | 0.378% |
| 160 | Plasma retinol-binding protein OS=Bos taurus OX=9913 GN=RBP4 PE=3 SV=1 | A0A3Q1MSW9 | 29 kDa | 5.66 | Transport/Carrier/Binding | 3 | 0.035% | 5 | 0.056% | 0 | 0.000% | 0 | 0.000% | 0 | 0.000% | 0 | 0.000% | 17 | 0.161% |
| 161 | Procollagen C-endopeptidase enhancer OS=Bos taurus OX=9913 GN=PCOLCE PE=2 SV=1 | Q2HJB6 | 48 kDa | 8.15 | Enzyme/Protease | 6 | 0.070% | 4 | 0.044% | 7 | 0.073% | 5 | 0.058% | 6 | 0.079% | 7 | 0.068% | 10 | 0.094% |
| 162 | Prostaglandin-H2 D-isomerase OS=Bos taurus OX=9913 GN=PTGDS PE=1 SV=1 | O02853 | 21 kDa | 6.12 | Transport/Carrier/Binding | 0 | 0.000% | 0 | 0.000% | 0 | 0.000% | 0 | 0.000% | 0 | 0.000% | 0 | 0.000% | 3 | 0.028% |
| 163 | Protein AMBP OS=Bos taurus OX=9913 GN=KIF12 PE=3 SV=3 | F1MMK9 | 53 kDa | 8.87 | Transport/Carrier/Binding | 18 | 0.209% | 18 | 0.200% | 15 | 0.157% | 13 | 0.151% | 11 | 0.145% | 22 | 0.214% | 19 | 0.179% |
| 164 | Protein C, inactivator of coagulation factors Va and VIIIa OS=Bos taurus OX=9913 GN=PROC PE=4 SV=2 | A0A140T851 (+1) | 51 kDa | 6.06 | Coagulation | 0 | 0.000% | 2 | 0.022% | 0 | 0.000% | 0 | 0.000% | 3 | 0.039% | 2 | 0.019% | 0 | 0.000% |
| 165 | Protein HP-20 homolog OS=Bos taurus OX=9913 PE=2 SV=1 | Q2KIT0 | 21 kDa | 8.86 | Structural/Cytoskeleton | 45 | 0.522% | 51 | 0.566% | 55 | 0.577% | 69 | 0.803% | 48 | 0.631% | 54 | 0.525% | 31 | 0.293% |
| 166 | Protein HP-25 homolog 1 OS=Bos taurus OX=9913 PE=1 SV=1 | Q2KIX7 | 23 kDa | 5.71 | Structural/Cytoskeleton | 19 | 0.220% | 25 | 0.278% | 30 | 0.315% | 34 | 0.396% | 32 | 0.421% | 28 | 0.272% | 16 | 0.151% |
| 167 | Protein HP-25 homolog 2 OS=Bos taurus OX=9913 PE=2 SV=1 | Q2KIU3 | 23 kDa | 5.29 | Structural/Cytoskeleton | 21 | 0.243% | 21 | 0.233% | 30 | 0.315% | 20 | 0.233% | 17 | 0.223% | 22 | 0.214% | 22 | 0.208% |
| 168 | Protein S OS=Bos taurus OX=9913 GN=PROS1 PE=4 SV=1 | A0A3Q1MMH1 (+1) | 71 kDa | 5.42 | Coagulation | 4 | 0.046% | 6 | 0.067% | 7 | 0.073% | 4 | 0.047% | 0 | 0.000% | 2 | 0.019% | 6 | 0.057% |
| 169 | Proteoglycan 4 OS=Bos taurus OX=9913 GN=PRG4 PE=4 SV=1 | A0A3Q1LZ67 | 158 kDa | 9.2 | Extracellular Matrix | 42 | 0.487% | 33 | 0.367% | 46 | 0.483% | 48 | 0.559% | 33 | 0.434% | 50 | 0.486% | 46 | 0.434% |
| 170 | Regakine-1 OS=Bos taurus OX=9913 PE=1 SV=2 | P82943 | 10 kDa | 8.81 | Immunity/Immunity Related | 0 | 0.000% | 0 | 0.000% | 0 | 0.000% | 0 | 0.000% | 0 | 0.000% | 0 | 0.000% | 4 | 0.038% |
| 171 | Serpin family A member 5 OS=Bos taurus OX=9913 GN=SERPINA5 PE=3 SV=1 | A0A452DHZ3 (+1) | 44 kDa | 9.36 | Protease Inhibitor | 12 | 0.139% | 14 | 0.156% | 10 | 0.105% | 18 | 0.209% | 14 | 0.184% | 3 | 0.029% | 13 | 0.123% |
| 172 | Serpin family A member 7 OS=Bos taurus OX=9913 GN=SERPINA7 PE=2 SV=1 | Q3SYR0 (+1) | 46 kDa | 5.51 | Protease Inhibitor | 0 | 0.000% | 0 | 0.000% | 5 | 0.052% | 9 | 0.105% | 14 | 0.184% | 9 | 0.087% | 19 | 0.179% |
| 173 | Serpin family D member 1 OS=Bos taurus OX=9913 GN=SERPIND1 PE=3 SV=1 | F6R4P6 | 63 kDa | 6.23 | Protease Inhibitor | 25 | 0.290% | 34 | 0.378% | 22 | 0.231% | 19 | 0.221% | 12 | 0.158% | 25 | 0.243% | 48 | 0.453% |
| 174 | Serum amyloid A-4 protein OS=Bos taurus OX=9913 GN=SAA4 PE=2 SV=1 | Q32L76 | 15 kDa | 6.8 | Apolipoprotein | 11 | 0.127% | 8 | 0.089% | 12 | 0.126% | 10 | 0.116% | 10 | 0.131% | 5 | 0.049% | 4 | 0.038% |
| 175 | Serum amyloid P-component OS=Bos taurus OX=9913 GN=APCS PE=2 SV=1 | Q3T004 | 25 kDa | 8.42 | Apolipoprotein | 40 | 0.464% | 41 | 0.455% | 41 | 0.430% | 39 | 0.454% | 42 | 0.552% | 40 | 0.389% | 11 | 0.104% |
| 176 | SHBG protein OS=Bos taurus OX=9913 GN=SHBG PE=2 SV=1 | A5PKC2 | 43 kDa | 5.43 | Transport/Carrier/Binding | 0 | 0.000% | 0 | 0.000% | 0 | 0.000% | 0 | 0.000% | 0 | 0.000% | 0 | 0.000% | 13 | 0.123% |
| 177 | Superoxide dismutase [Cu-Zn] OS=Bos taurus OX=9913 GN=SOD3 PE=1 SV=1 | F6R4N7 | 27 kDa | 6.63 | Enzyme/Protease | 4 | 0.046% | 5 | 0.056% | 3 | 0.031% | 6 | 0.070% | 9 | 0.118% | 3 | 0.029% | 7 | 0.066% |
| 178 | Sushi domain-containing protein OS=Bos taurus OX=9913 PE=4 SV=2 | G3N0S9 | 21 kDa | 5.37 | Complement | 21 | 0.243% | 22 | 0.244% | 23 | 0.241% | 23 | 0.268% | 14 | 0.184% | 39 | 0.379% | 17 | 0.161% |
| 179 | Transforming growth factor-beta-induced protein ig-h3 OS=Bos taurus OX=9913 GN=TGFBI PE=1 SV=2 | P55906 | 74 kDa | 6.69 | Cell-Cell Adhesion | 10 | 0.116% | 7 | 0.078% | 0 | 0.000% | 8 | 0.093% | 9 | 0.118% | 0 | 0.000% | 4 | 0.038% |
| 180 | Triosephosphate isomerase OS=Bos taurus OX=9913 GN=TPI1 PE=2 SV=3 | Q5E956 | 27 kDa | 5.89 | Enzyme/Protease | 4 | 0.046% | 2 | 0.022% | 3 | 0.031% | 3 | 0.035% | 3 | 0.039% | 3 | 0.029% | 0 | 0.000% |
| 181 | Versican OS=Bos taurus OX=9913 GN=VCAN PE=1 SV=2 | F1MZ85 | 369 kDa | 4.46 | Extracellular Matrix | 3 | 0.035% | 0 | 0.000% | 2 | 0.021% | 0 | 0.000% | 0 | 0.000% | 0 | 0.000% | 0 | 0.000% |
| 182 | Vitronectin OS=Bos taurus OX=9913 GN=VTN PE=1 SV=1 | Q3ZBS7 | 54 kDa | 5.79 | Cell-Cell Adhesion | 14 | 0.162% | 17 | 0.189% | 21 | 0.220% | 10 | 0.116% | 13 | 0.171% | 18 | 0.175% | 19 | 0.179% |

**Supplemental Table 3: Mass spectroscopy data for synovial fluid stock and FBS stock**

| | | | | | | Synovial Fluid (stock) | | FBS (stock) | |
|---|---|---|---|---|---|---|---|---|---|
| | Identified Proteins | Accession Number | Molecular Weight (MW) | Theoretical Isoelectric Point (pI) | Biological Process | Total Spectrum Count | Percentage of Total Spectral Counts | Total Spectrum Count | Percentage of Total Spectral Counts |
| 1 | Cluster of Albumin OS=Bos taurus OX=9913 GN=ALB PE=4 SV=1 (A0A140T897) | A0A140T897 [2] | 69 kDa | 5.77 | Transport/Carrier/Binding | 2261 | 21.932% | 2484 | 22.656% |
| 2 | Cluster of Beta-1 metal-binding globulin OS=Bos taurus OX=9913 GN=TF PE=1 SV=2 (G3X6N3) | G3X6N3 [3] | 78 kDa | 6.63 | Transport/Carrier/Binding | 643 | 6.237% | 743 | 6.777% |
| 3 | Complement C3 OS=Bos taurus OX=9913 GN=C3 PE=1 SV=2 | Q2UVX4 | 187 kDa | 6.37 | Complement | 458 | 4.443% | 353 | 3.220% |
| 4 | Uncharacterized protein OS=Bos taurus OX=9913 PE=1 SV=1 | G3N0V0 | 36 kDa | 8.05 | Complement | 341 | 3.308% | 21 | 0.192% |
| 5 | Alpha-2-macroglobulin OS=Bos taurus OX=9913 GN=A2M PE=1 SV=2 | Q7SIH1 | 168 kDa | 5.68 | Protease Inhibitor | 320 | 3.104% | 537 | 4.898% |
| 6 | Uncharacterized protein OS=Bos taurus OX=9913 PE=1 SV=1 | A0A3Q1M3L6 | 40 kDa | 5.16 | Immunoglobulin | 286 | 2.774% | 51 | 0.465% |
| 7 | Cluster of Fibronectin OS=Bos taurus OX=9913 GN=FN1 PE=1 SV=4 (P07589) | P07589 [3] | 272 kDa | 5.28 | Structural/Cytoskeleton | 282 | 2.735% | 42 | 0.383% |
| 8 | Cluster of Uncharacterized protein OS=Bos taurus OX=9913 PE=4 SV=3 (F1N160) | F1N160 [3] | 34 kDa | 8.86 | Immunity/Immunity Related | 247 | 2.396% | 21 | 0.192% |
| 9 | Actin-depolymerizing factor OS=Bos taurus OX=9913 GN=GSN PE=1 SV=2 | F1N1I6 | 92 kDa | 6.42 | Structural/Cytoskeleton | 225 | 2.183% | 52 | 0.474% |
| 10 | Serpin A3-1 OS=Bos taurus OX=9913 GN=SERPINA3-1 PE=3 SV=1 | A0A452DJK6 (+1) | 52 kDa | 5.99 | Protease Inhibitor | 197 | 1.911% | 161 | 1.468% |
| 11 | Cluster of C4a anaphylatoxin OS=Bos taurus OX=9913 GN=LOC107131209 PE=4 SV=3 (F1MVK1) | F1MVK1 [3] | 192 kDa | 7.26 | Protease Inhibitor | 194 | 1.882% | 174 | 1.587% |
| 12 | Cluster of Serpin A3-7 OS=Bos taurus OX=9913 GN=SERPINA3-7 PE=1 SV=1 (A0A0A0MP92) | A0A0A0MP92 [2] | 47 kDa | 5.61 | Protease Inhibitor | 183 | 1.775% | 117 | 1.067% |
| 13 | Cluster of Ceruloplasmin OS=Bos taurus OX=9913 GN=CP PE=1 SV=1 (A0A3Q1NJB1) | A0A3Q1NJB1 [2] | 121 kDa | 5.67 | Transport/Carrier/Binding | 162 | 1.571% | 5 | 0.046% |
| 14 | Cluster of Hemopexin OS=Bos taurus OX=9913 GN=HPX PE=2 SV=1 (Q3SZV7) | Q3SZV7 [2] | 52 kDa | 7.1 | Transport/Carrier/Binding | 157 | 1.523% | 99 | 0.903% |
| 15 | Cluster of Serpin A3-3 OS=Bos taurus OX=9913 GN=SERPINA3-3 PE=1 SV=2 (Q3ZEJ6) | Q3ZEJ6 [3] | 46 kDa | 5.73 | Protease Inhibitor | 149 | 1.445% | 82 | 0.748% |
| 16 | Cluster of Plasminogen OS=Bos taurus OX=9913 GN=PLG PE=3 SV=2 (E1B726) | E1B726 [2] | 91 kDa | 7.71 | Coagulation | 142 | 1.377% | 153 | 1.395% |
| 17 | Cluster of Gc-globulin OS=Bos taurus OX=9913 GN=GC PE=4 SV=2 (F1N5M2) | F1N5M2 [2] | 53 kDa | 5.24 | Transport/Carrier/Binding | 130 | 1.261% | 225 | 2.052% |
| 18 | Cluster of Inter-alpha-trypsin inhibitor heavy chain H4 OS=Bos taurus OX=9913 GN=ITIH4 PE=3 SV=3 (F1MMD7) | F1MMD7 [3] | 102 kDa | 5.99 | Protease Inhibitor | 128 | 1.242% | 143 | 1.304% |
| 19 | Cluster of Apolipoprotein A-I OS=Bos taurus OX=9913 GN=APOA1 PE=1 SV=3 (P15497) | P15497 [2] | 30 kDa | 5.39 | Apolipoprotein | 118 | 1.145% | 124 | 1.131% |
| 20 | Alpha-1-antiproteinase OS=Bos taurus OX=9913 GN=SERPINA1 PE=1 SV=1 | P34955 | 46 kDa | 5.98 | Protease Inhibitor | 110 | 1.067% | 833 | 7.598% |
| 21 | Cluster of Complement factor H OS=Bos taurus OX=9913 GN=CFH PE=1 SV=3 (Q28085) | Q28085 [6] | 140 kDa | 6.33 | Complement | 108 | 1.048% | 85 | 0.775% |
| 22 | Cluster of Antithrombin-III OS=Bos taurus OX=9913 GN=SERPINC1 PE=3 SV=1 (A0A3Q1NJR8) | A0A3Q1NJR8 [2] | 60 kDa | 8.85 | Protease Inhibitor | 106 | 1.028% | 120 | 1.094% |
| 23 | Prothrombin OS=Bos taurus OX=9913 GN=F2 PE=1 SV=2 | P00735 | 71 kDa | 5.4 | Coagulation | 105 | 1.019% | 85 | 0.775% |
| 24 | Complement factor B OS=Bos taurus OX=9913 GN=CFB PE=1 SV=2 | P81187 | 85 kDa | 7.68 | Complement | 103 | 0.999% | 86 | 0.784% |
| 25 | Cluster of Fibrinogen beta chain OS=Bos taurus OX=9913 GN=FGB PE=4 SV=1 (A0A3Q1MG04) | A0A3Q1MG04 [3] | 57 kDa | 8.33 | Coagulation | 80 | 0.776% | 12 | 0.109% |
| 26 | Alpha-1-acid glycoprotein OS=Bos taurus OX=9913 GN=ORM1 PE=2 SV=1 | Q3SZR3 (+1) | 23 kDa | 5.67 | Transport/Carrier/Binding | 79 | 0.766% | 62 | 0.565% |
| 27 | Cluster of Bradykinin OS=Bos taurus OX=9913 GN=KNG1 PE=4 SV=1 (A0A140T8C8) | A0A140T8C8 [2] | 69 kDa | 6.09 | Protease Inhibitor | 78 | 0.757% | 93 | 0.848% |
| 28 | Complement C5a anaphylatoxin OS=Bos taurus OX=9913 GN=C5 PE=1 SV=3 | F1MY85 | 189 kDa | 6.16 | Complement | 77 | 0.747% | 61 | 0.556% |
| 29 | Inter-alpha-trypsin inhibitor heavy chain H1 OS=Bos taurus OX=9913 GN=ITIH1 PE=1 SV=1 | Q0VCM5 | 101 kDa | 7.52 | Protease Inhibitor | 75 | 0.728% | 45 | 0.410% |
| 30 | Fibrinogen alpha chain OS=Bos taurus OX=9913 GN=FGA PE=4 SV=1 | F6QND5 | 95 kDa | 5.69 | Coagulation | 73 | 0.708% | 24 | 0.219% |

| | | | | | | | | | |
|---|---|---|---|---|---|---|---|---|---|
| 31 | Cluster of Uncharacterized protein OS=Bos taurus OX=9913 PE=1 SV=1 (A0A3Q1M032) | A0A3Q1M032 [2] | 40 kDa | 5.77 | N/A | 70 | 0.679% | 25 | 0.228% |
| 32 | Primary amine oxidase, liver isozyme OS=Bos taurus OX=9913 PE=1 SV=1 | Q29437 | 85 kDa | 5.58 | Enzyme/Protease | 70 | 0.679% | 4 | 0.036% |
| 33 | Apolipoprotein A-IV OS=Bos taurus OX=9913 GN=APOA4 PE=3 SV=2 | F1N3Q7 | 52 kDa | 5.5 | Apolipoprotein | 64 | 0.621% | 13 | 0.119% |
| 34 | Fibrinogen gamma-B chain OS=Bos taurus OX=9913 GN=FGG PE=4 SV=1 | F1MGU7 (+1) | 50 kDa | 5.38 | Coagulation | 62 | 0.601% | 9 | 0.082% |
| 36 | Proteoglycan 4 OS=Bos taurus OX=9913 GN=PRG4 PE=4 SV=3 | F1MDS0 | 129 kDa | 9.2 | Extracellular Matrix | 60 | 0.582% | 0 | 0.000% |
| 37 | ITIH2 protein OS=Bos taurus OX=9913 GN=ITIH2 PE=1 SV=1 | A5D7R6 | 106 kDa | 7.75 | Protease Inhibitor | 59 | 0.572% | 135 | 1.231% |
| 38 | Aggrecan core protein OS=Bos taurus OX=9913 GN=ACAN PE=3 SV=2 | F1N367 (+1) | 243 kDa | 4.17 | Extracellular Matrix | 57 | 0.553% | 0 | 0.000% |
| 39 | Cluster of Clusterin OS=Bos taurus OX=9913 GN=CLU PE=1 SV=1 (P17697) | P17697 [2] | 51 kDa | 5.72 | Apolipoprotein | 57 | 0.553% | 26 | 0.237% |
| 40 | Alpha-2-HS-glycoprotein OS=Bos taurus OX=9913 GN=AHSG PE=1 SV=2 | P12763 | 38 kDa | 5.1 | Protease Inhibitor | 55 | 0.534% | 772 | 7.041% |
| 41 | Transthyretin OS=Bos taurus OX=9913 GN=TTR PE=1 SV=1 | O46375 | 16 kDa | 5.91 | Transport/Carrier/Binding | 55 | 0.534% | 51 | 0.465% |
| 42 | Hemoglobin subunit beta OS=Bos taurus OX=9913 GN=HBB PE=1 SV=1 | P02070 | 16 kDa | 7.02 | Transport/Carrier/Binding | 54 | 0.524% | 32 | 0.292% |
| 43 | Alpha-1B-glycoprotein OS=Bos taurus OX=9913 GN=A1BG PE=1 SV=1 | Q2KJF1 | 54 kDa | 5.37 | Immunity/Immunity Related | 53 | 0.514% | 60 | 0.547% |
| 44 | Serpin A3-8 OS=Bos taurus OX=9913 GN=SERPINA3-8 PE=2 SV=1 | A6QPQ2 | 47 kDa | 5.49 | Protease Inhibitor | 52 | 0.504% | 19 | 0.173% |
| 45 | Uncharacterized protein OS=Bos taurus OX=9913 PE=1 SV=1 | A0A3Q1LPG0 | 36 kDa | 7.59 | Immunity/Immunity Related | 51 | 0.495% | 0 | 0.000% |
| 46 | Cartilage oligomeric matrix protein OS=Bos taurus OX=9913 GN=COMP PE=1 SV=2 | P35445 | 82 kDa | 4.36 | Extracellular Matrix | 49 | 0.475% | 18 | 0.164% |
| 47 | Cluster of Actin, cytoplasmic 1 OS=Bos taurus OX=9913 GN=ACTB PE=1 SV=1 (P60712) | P60712 [4] | 42 kDa | 5.29 | Structural/Cytoskeleton | 47 | 0.456% | 35 | 0.319% |
| 48 | Cluster of Complement component C9 OS=Bos taurus OX=9913 GN=C9 PE=2 SV=1 (Q3MHN2) | Q3MHN2 [2] | 62 kDa | 5.57 | Complement | 47 | 0.456% | 19 | 0.173% |
| 50 | SERPIN domain-containing protein OS=Bos taurus OX=9913 GN=LOC784932 PE=1 SV=1 | A0A0A0MPA0 | 47 kDa | 5.44 | Protease Inhibitor | 47 | 0.456% | 39 | 0.356% |
| 51 | Cluster of Uncharacterized protein OS=Bos taurus OX=9913 GN=LOC506828 PE=3 SV=3 (F1MJK3) | F1MJK3 [3] | 163 kDa | 6.91 | Protease Inhibitor | 46 | 0.446% | 122 | 1.113% |
| 52 | Cluster of Uncharacterized protein OS=Bos taurus OX=9913 PE=1 SV=3 (F1MZ96) | F1MZ96 [2] | 27 kDa | 5.94 | Immunity/Immunity Related | 46 | 0.446% | 14 | 0.128% |
| 53 | Apolipoprotein H OS=Bos taurus OX=9913 GN=APOH PE=4 SV=1 | A0A140T843 (+1) | 38 kDa | 8.55 | Apolipoprotein | 43 | 0.417% | 57 | 0.520% |
| 54 | Complement component C6 OS=Bos taurus OX=9913 GN=C6 PE=3 SV=2 | F1MM86 | 105 kDa | 6.74 | Complement | 43 | 0.417% | 2 | 0.018% |
| 55 | Cluster of Complement factor I OS=Bos taurus OX=9913 GN=CFI PE=1 SV=1 (A0A3Q1LGM4) | A0A3Q1LGM4 [4] | 65 kDa | 8.13 | Complement | 42 | 0.407% | 35 | 0.319% |
| 56 | Cluster of Histidine-rich glycoprotein OS=Bos taurus OX=9913 GN=HRG PE=4 SV=3 (F1MKS5) | F1MKS5 [2] | 61 kDa | 7.14 | Protease Inhibitor | 42 | 0.407% | 5 | 0.046% |
| 57 | Alpha-2-antiplasmin OS=Bos taurus OX=9913 GN=SERPINF2 PE=1 SV=2 | P28800 | 55 kDa | 5.71 | Protease Inhibitor | 40 | 0.388% | 107 | 0.976% |
| 58 | Pigment epithelium-derived factor OS=Bos taurus OX=9913 GN=SERPINF1 PE=1 SV=1 | Q95121 | 46 kDa | 6.31 | Coagulation | 40 | 0.388% | 47 | 0.429% |
| 59 | Serpin family D member 1 OS=Bos taurus OX=9913 GN=SERPIND1 PE=3 SV=1 | F6R4P6 | 63 kDa | 6.23 | Protease Inhibitor | 38 | 0.369% | 62 | 0.565% |
| 61 | Afamin OS=Bos taurus OX=9913 GN=AFM PE=1 SV=1 | G3MYZ3 | 70 kDa | 5.57 | Transport/Carrier/Binding | 36 | 0.349% | 48 | 0.438% |
| 62 | Complement component 8 subunit beta OS=Bos taurus OX=9913 GN=C8B PE=3 SV=3 | F1N102 | 67 kDa | 8.15 | Complement | 36 | 0.349% | 8 | 0.073% |
| 63 | Extracellular matrix protein 1 OS=Bos taurus OX=9913 GN=ECM1 PE=4 SV=1 | A0A3Q1M5Q6 | 63 kDa | 6.6 | Extracellular Matrix | 35 | 0.340% | 6 | 0.055% |
| 64 | Angiotensinogen OS=Bos taurus OX=9913 GN=AGT PE=1 SV=2 | P01017 | 51 kDa | 6.49 | Protease Inhibitor | 34 | 0.330% | 67 | 0.611% |
| 65 | C4b-binding protein alpha chain OS=Bos taurus OX=9913 GN=C4BPA PE=2 SV=1 | Q28065 | 69 kDa | 5.66 | Complement | 34 | 0.330% | 16 | 0.146% |
| 66 | Hemoglobin subunit alpha OS=Bos taurus OX=9913 GN=HBA PE=1 SV=2 | P01966 | 15 kDa | 8.19 | Transport/Carrier/Binding | 34 | 0.330% | 49 | 0.447% |
| 67 | Leucine rich alpha-2-glycoprotein 1 OS=Bos taurus OX=9913 GN=LRG1 PE=1 SV=1 | F6RMV5 | 39 kDa | 6.26 | Structural/Cytoskeleton | 33 | 0.320% | 26 | 0.237% |

| | | | | | | | | | |
|---|---|---|---|---|---|---|---|---|---|
| 68 | Cluster of Serpin family G member 1 OS=Bos taurus OX=9913 GN=SERPING1 PE=3 SV=3 (E1BMJ0) | E1BMJ0 [2] | 52 kDa | 6.28 | Protease Inhibitor | 32 | 0.310% | 41 | 0.374% |
| 69 | Protein HP-20 homolog OS=Bos taurus OX=9913 PE=2 SV=1 | Q2KIT0 | 21 kDa | 8.86 | Structural/Cytoskeleton | 31 | 0.301% | 7 | 0.064% |
| 70 | Protein HP-25 homolog 2 OS=Bos taurus OX=9913 PE=2 SV=1 | Q2KIU3 | 23 kDa | 5.29 | Structural/Cytoskeleton | 30 | 0.291% | 5 | 0.046% |
| 71 | Uncharacterized protein OS=Bos taurus OX=9913 PE=1 SV=1 | A0A3Q1LRW4 (+1) | 47 kDa | 8.2 | Immunity/Immunity Related | 28 | 0.272% | 0 | 0.000% |
| 72 | Alpha-1-microglobulin OS=Bos taurus OX=9913 GN=KIF12 PE=3 SV=3 | F1MMK9 | 53 kDa | 8.87 | N/A | 27 | 0.262% | 41 | 0.374% |
| 73 | Complement C8 alpha chain OS=Bos taurus OX=9913 GN=C8A PE=3 SV=1 | F1MX87 | 66 kDa | 6.24 | Complement | 25 | 0.243% | 8 | 0.073% |
| 74 | Plasma retinol-binding protein OS=Bos taurus OX=9913 GN=RBP4 PE=3 SV=1 | A0A3Q1MSW9 (+2) | 29 kDa | 5.66 | Transport/Carrier/Binding | 25 | 0.243% | 21 | 0.192% |
| 75 | Zinc-alpha-2-glycoprotein OS=Bos taurus OX=9913 GN=AZGP1 PE=3 SV=1 | A0A452DK44 (+1) | 34 kDa | 5.11 | Immunity/Immunity Related | 24 | 0.233% | 2 | 0.018% |
| 76 | Cluster of Ig-like domain-containing protein OS=Bos taurus OX=9913 PE=4 SV=1 (A0A3Q1ML26) | A0A3Q1ML26 [6] | 23 kDa | 9.08 | Immunoglobulin | 23 | 0.223% | 5 | 0.046% |
| 77 | Cluster of Trypsin OS=Sus scrofa OX=9823 (P00761) | P00761 [2] | 24 kDa | 8.26 | Enzyme/Protease | 23 | 0.223% | 37 | 0.337% |
| 78 | Cluster of Complement component C7 OS=Bos taurus OX=9913 GN=C7 PE=2 SV=1 (Q29RQ1) | Q29RQ1 [3] | 93 kDa | 6.9 | Complement | 22 | 0.213% | 24 | 0.219% |
| 79 | Lumican OS=Bos taurus OX=9913 GN=LUM PE=1 SV=1 | Q05443 | 39 kDa | 5.93 | Extracellular Matrix | 22 | 0.213% | 21 | 0.192% |
| 80 | Protein HP-25 homolog 1 OS=Bos taurus OX=9913 PE=1 SV=1 | Q2KIX7 | 23 kDa | 5.71 | Structural/Cytoskeleton | 22 | 0.213% | 3 | 0.027% |
| 81 | Haptoglobin OS=Bos taurus OX=9913 GN=HP PE=3 SV=1 | A0A3Q1MB98 (+1) | 44 kDa | 8.09 | Complement | 21 | 0.204% | 0 | 0.000% |
| 82 | Vitronectin OS=Bos taurus OX=9913 GN=VTN PE=1 SV=1 | Q3ZBS7 | 54 kDa | 5.79 | Cell-Cell Adhesion | 20 | 0.194% | 32 | 0.292% |
| 83 | Cystatin-C OS=Bos taurus OX=9913 GN=CST3 PE=1 SV=2 | P01035 | 16 kDa | 9.03 | Protease Inhibitor | 19 | 0.184% | 14 | 0.128% |
| 84 | Uncharacterized protein OS=Bos taurus OX=9913 GN=LOC525947 PE=1 SV=1 | A0A3Q1NEQ0 | 78 kDa | 6.8 | Transport/Carrier/Binding | 19 | 0.184% | 58 | 0.529% |
| 85 | Uncharacterized protein OS=Bos taurus OX=9913 GN=LOC528040 PE=4 SV=3 | E1B805 | 186 kDa | 6.54 | N/A | 19 | 0.184% | 4 | 0.036% |
| 86 | Amine oxidase OS=Bos taurus OX=9913 GN=LOC100138645 PE=3 SV=1 | A0A452DHX8 | 87 kDa | 5.56 | Oxidase | 18 | 0.175% | 5 | 0.046% |
| 87 | Cluster of CD5 molecule like OS=Bos taurus OX=9913 GN=CD5L PE=2 SV=1 (A6QNW7) | A6QNW7 [3] | 50 kDa | 5.24 | Immunity/Immunity Related | 18 | 0.175% | 4 | 0.036% |
| 88 | Cluster of Ig-like domain-containing protein OS=Bos taurus OX=9913 PE=4 SV=1 (A0A3Q1N2B9) | A0A3Q1N2B9 [4] | 15 kDa | 8.96 | Immunoglobulin | 17 | 0.165% | 0 | 0.000% |
| 89 | Alpha-amylase OS=Bos taurus OX=9913 GN=AMY2B PE=3 SV=1 | F1MJQ3 | 57 kDa | 5.87 | Enzyme/Protease | 16 | 0.155% | 13 | 0.119% |
| 90 | Cluster of Tetranectin OS=Bos taurus OX=9913 GN=CLEC3B PE=4 SV=1 (A0A452DIU4) | A0A452DIU4 [4] | 31 kDa | 6.52 | sport/Carrier/Binding/Carrier/Bi | 16 | 0.155% | 15 | 0.137% |
| 91 | Fibulin-1 OS=Bos taurus OX=9913 GN=FBLN1 PE=1 SV=1 | A0A3Q1MG73 | 74 kDa | 4.9 | Coagulation | 16 | 0.155% | 36 | 0.328% |
| 92 | Beta-2-microglobulin OS=Bos taurus OX=9913 PE=3 SV=1 | A0A3Q1MU93 (+1) | 14 kDa | 9.1 | Immunity/Immunity Related | 15 | 0.146% | 4 | 0.036% |
| 93 | Fetuin-B OS=Bos taurus OX=9913 GN=FETUB PE=1 SV=1 | Q58D62 | 43 kDa | 5.59 | Protease Inhibitor | 15 | 0.146% | 135 | 1.231% |
| 94 | Paraoxonase OS=Bos taurus OX=9913 GN=PON1 PE=1 SV=1 | Q2KIW1 | 40 kDa | 5.17 | Enzyme/Protease | 15 | 0.146% | 0 | 0.000% |
| 95 | Phospholipid transfer protein OS=Bos taurus OX=9913 GN=PLTP PE=2 SV=1 | Q58DL9 | 56 kDa | 6.32 | Immunity/Immunity Related | 15 | 0.146% | 6 | 0.055% |
| 96 | C3/C5 convertase OS=Bos taurus OX=9913 GN=C2 PE=4 SV=1 | A0A3S5ZPC6 (+1) | 87 kDa | 8.86 | Complement | 14 | 0.136% | 0 | 0.000% |
| 97 | Apolipoprotein E OS=Bos taurus OX=9913 GN=APOE PE=3 SV=2 | A0A140T881 (+2) | 37 kDa | 5.55 | Apolipoprotein | 13 | 0.126% | 27 | 0.246% |
| 98 | Complement C1s subcomponent OS=Bos taurus OX=9913 GN=C1S PE=2 SV=2 | Q0VCX1 | 77 kDa | 4.97 | Complement | 13 | 0.126% | 0 | 0.000% |
| 99 | Hyaluronan and proteoglycan link protein 1 OS=Bos taurus OX=9913 GN=HAPLN1 PE=2 SV=1 | P55252 | 40 kDa | 7.66 | Extracellular Matrix | 13 | 0.126% | 0 | 0.000% |
| 100 | Cluster of Conglutinin OS=Bos taurus OX=9913 GN=CGN1 PE=1 SV=2 (P23805) | P23805 [2] | 38 kDa | 5.63 | Extracellular Matrix | 12 | 0.116% | 0 | 0.000% |
| 101 | Pantetheinase OS=Bos taurus OX=9913 GN=VNN1 PE=3 SV=1 | A0A140T891 (+1) | 57 kDa | 5.26 | Enzyme/Protease | 12 | 0.116% | 30 | 0.274% |

| | | | | | | | | | |
|---|---|---|---|---|---|---|---|---|---|
| 102 | Serpin peptidase inhibitor, clade A (Alpha-1 antiproteinase, antitrypsin), member 7 OS=Bos taurus OX=9913 GN=SERPINA7 PE=2 SV=1 | Q3SYR0 (+1) | 46 kDa | 5.51 | Protease Inhibitor | 12 | 0.116% | 23 | 0.210% |
| 103 | Uncharacterized protein OS=Bos taurus OX=9913 PE=4 SV=2 | G3N0S9 | 21 kDa | 5.37 | Complement | 12 | 0.116% | 8 | 0.073% |
| 104 | Cathelicidin-1 OS=Bos taurus OX=9913 GN=CATHL1 PE=1 SV=2 | P22226 | 18 kDa | 11.53 | Immunity/Immunity Related | 11 | 0.107% | 0 | 0.000% |
| 105 | Chondroadherin OS=Bos taurus OX=9913 GN=CHAD PE=4 SV=2 | F1MYE4 (+1) | 41 kDa | 9.43 | Extracellular Matrix | 11 | 0.107% | 0 | 0.000% |
| 106 | Cluster of Christmas factor OS=Bos taurus OX=9913 GN=F9 PE=4 SV=3 (F1MBC5) | F1MBC5 [3] | 50 kDa | 5.9 | Enzyme/Protease | 11 | 0.107% | 9 | 0.082% |
| 107 | Cluster of Serum amyloid A protein OS=Bos taurus OX=9913 GN=LOC104968478 PE=3 SV=2 (E1BJF9) | E1BJF9 [2] | 20 kDa | 8.9 | Apolipoprotein | 11 | 0.107% | 0 | 0.000% |
| 108 | Peptidoglycan recognition protein 1 OS=Bos taurus OX=9913 GN=PGLYRP1 PE=1 SV=1 | Q8SPP7 | 21 kDa | 9.38 | Immunity/Immunity Related | 11 | 0.107% | 0 | 0.000% |
| 109 | Procollagen C-endopeptidase enhancer OS=Bos taurus OX=9913 GN=PCOLCE PE=2 SV=1 | Q2HJB6 | 48 kDa | 8.15 | Enzyme/Protease | 11 | 0.107% | 0 | 0.000% |
| 110 | Serum amyloid P-component OS=Bos taurus OX=9913 GN=APCS PE=2 SV=1 | Q3T004 | 25 kDa | 8.42 | Apolipoprotein | 11 | 0.107% | 0 | 0.000% |
| 111 | Cluster of Plasma serine protease inhibitor OS=Bos taurus OX=9913 GN=SERPINA5 PE=3 SV=1 (A0A452DHZ3) | A0A452DHZ3 [3] | 44 kDa | 9.36 | Coagulation | 10 | 0.097% | 31 | 0.283% |
| 112 | Ig-like domain-containing protein OS=Bos taurus OX=9913 PE=4 SV=2 | G3MXG6 | 15 kDa | 8.56 | Immunoglobulin | 10 | 0.097% | 4 | 0.036% |
| 113 | SHBG protein OS=Bos taurus OX=9913 GN=SHBG PE=2 SV=1 | A5PKC2 | 43 kDa | 5.43 | Transport/Carrier/Binding | 10 | 0.097% | 9 | 0.082% |
| 114 | Lipopolysaccharide-binding protein OS=Bos taurus OX=9913 GN=LBP PE=3 SV=1 | A0A3Q1LKB0 (+2) | 53 kDa | 6.38 | Immunity/Immunity Related | 9 | 0.087% | 0 | 0.000% |
| 115 | Adiponectin OS=Bos taurus OX=9913 GN=ADIPOQ PE=4 SV=1 | A0A3Q1M564 (+1) | 35 kDa | 5.36 | Cell-Cell Adhesion | 8 | 0.078% | 15 | 0.137% |
| 116 | Apolipoprotein C-III OS=Bos taurus OX=9913 GN=APOC3 PE=1 SV=2 | P19035 | 11 kDa | 4.73 | Apolipoprotein | 8 | 0.078% | 6 | 0.055% |
| 117 | Apolipoprotein D OS=Bos taurus OX=9913 GN=APOD PE=3 SV=3 | F1MS32 (+1) | 24 kDa | 5.07 | Apolipoprotein | 8 | 0.078% | 7 | 0.064% |
| 118 | Ig-like domain-containing protein OS=Bos taurus OX=9913 PE=4 SV=1 | A0A3Q1MSF6 | 11 kDa | 5 | Immunoglobulin | 8 | 0.078% | 0 | 0.000% |
| 119 | Ig-like domain-containing protein OS=Bos taurus OX=9913 PE=4 SV=1 | A0A3Q1LSF0 (+1) | 15 kDa | 9.82 | Immunoglobulin | 8 | 0.078% | 4 | 0.036% |
| 120 | Serum amyloid A-4 protein OS=Bos taurus OX=9913 GN=SAA4 PE=2 SV=1 | Q32L76 | 15 kDa | 6.8 | Apolipoprotein | 8 | 0.078% | 0 | 0.000% |
| 121 | Superoxide dismutase [Cu-Zn] OS=Bos taurus OX=9913 GN=SOD3 PE=1 SV=1 | F6R4N7 | 27 kDa | 6.63 | Enzyme/Protease | 8 | 0.078% | 0 | 0.000% |
| 122 | Carboxypeptidase B2 OS=Bos taurus OX=9913 GN=CPB2 PE=3 SV=1 | A0A3Q1N1A7 (+1) | 45 kDa | 8.69 | Enzyme/Protease | 7 | 0.068% | 5 | 0.046% |
| 123 | Complement factor D OS=Bos taurus OX=9913 GN=CFD PE=2 SV=1 | Q3T0A3 | 28 kDa | 6.85 | Complement | 7 | 0.068% | 4 | 0.036% |
| 124 | Glutathione peroxidase 3 OS=Bos taurus OX=9913 GN=GPX3 PE=2 SV=2 | P37141 | 26 kDa | 6 | Enzyme/Protease | 7 | 0.068% | 0 | 0.000% |
| 125 | Ig-like domain-containing protein OS=Bos taurus OX=9913 GN=LOC100300716 PE=4 SV=2 | G3N1H5 | 18 kDa | 9.86 | Immunoglobulin | 7 | 0.068% | 0 | 0.000% |
| 126 | Ig-like domain-containing protein OS=Bos taurus OX=9913 PE=4 SV=1 | A0A3Q1LI44 | 15 kDa | 8.94 | Immunoglobulin | 7 | 0.068% | 0 | 0.000% |
| 127 | Ig-like domain-containing protein OS=Bos taurus OX=9913 PE=4 SV=1 | A0A3Q1M667 (+1) | 25 kDa | 9.01 | Immunoglobulin | 7 | 0.068% | 0 | 0.000% |
| 128 | Prostaglandin-H2 D-isomerase OS=Bos taurus OX=9913 GN=PTGDS PE=1 SV=1 | O02853 | 21 kDa | 6.12 | Transport/Carrier/Binding | 7 | 0.068% | 0 | 0.000% |
| 129 | Adiponectin B OS=Bos taurus OX=9913 GN=C1QC PE=4 SV=1 | A0A3B0IZF8 | 26 kDa | 8.62 | Complement | 6 | 0.058% | 0 | 0.000% |
| 130 | ApoN protein OS=Bos taurus OX=9913 GN=APON PE=2 SV=1 | Q2KIH2 | 29 kDa | 5.36 | Apolipoprotein | 6 | 0.058% | 5 | 0.046% |
| 131 | Cluster of Ig-like domain-containing protein OS=Bos taurus OX=9913 PE=4 SV=1 (A0A3Q1MBT3) | A0A3Q1MBT3 [5] | 12 kDa | 6.52 | Immunoglobulin | 6 | 0.058% | 0 | 0.000% |
| 132 | Corticosteroid-binding globulin OS=Bos taurus OX=9913 GN=SERPINA6 PE=3 SV=1 | E1BF81 | 45 kDa | 5.54 | Protease Inhibitor | 6 | 0.058% | 16 | 0.146% |
| 133 | EGF containing fibulin extracellular matrix protein 1 OS=Bos taurus OX=9913 GN=EFEMP1 PE=1 SV=1 | A0A3Q1MIU1 (+2) | 50 kDa | 4.83 | Extracellular Matrix | 6 | 0.058% | 3 | 0.027% |
| 134 | Ig-like domain-containing protein OS=Bos taurus OX=9913 PE=4 SV=2 | G3N148 (+1) | 13 kDa | 9.07 | Immunoglobulin | 6 | 0.058% | 0 | 0.000% |
| 135 | Monocyte differentiation antigen CD14 OS=Bos taurus OX=9913 GN=CD14 PE=2 SV=1 | A6QNL0 (+1) | 40 kDa | 5.24 | Immunity/Immunity Related | 6 | 0.058% | 3 | 0.027% |

| | | | | | | | | | |
|---|---|---|---|---|---|---|---|---|---|
| 136 | Regakine-1 OS=Bos taurus OX=9913 PE=1 SV=2 | P82943 | 10 kDa | 8.81 | Immunity/Immunity Related | 6 | 0.058% | 3 | 0.027% |
| 137 | SERPINA10 protein OS=Bos taurus OX=9913 GN=SERPINA10 PE=2 SV=1 | A5PJ69 | 52 kDa | 6.05 | Protease Inhibitor | 6 | 0.058% | 18 | 0.164% |
| 138 | Complement subcomponent C1r OS=Bos taurus OX=9913 GN=C1R PE=4 SV=1 | A0A3Q1MGP1 (+1) | 81 kDa | 5.8 | Complement | 5 | 0.049% | 0 | 0.000% |
| 139 | Ig-like domain-containing protein OS=Bos taurus OX=9913 PE=4 SV=1 | A0A3Q1MI29 | 23 kDa | 8.13 | Immunoglobulin | 5 | 0.049% | 0 | 0.000% |
| 140 | Vitamin K-dependent protein S OS=Bos taurus OX=9913 GN=PROS1 PE=4 SV=1 | A0A3Q1MMH1 (+1) | 71 kDa | 5.42 | Coagulation | 5 | 0.049% | 11 | 0.100% |
| 141 | Apolipoprotein A-II OS=Bos taurus OX=9913 GN=APOA2 PE=1 SV=2 | P81644 | 11 kDa | 5.34 | Apolipoprotein | 4 | 0.039% | 11 | 0.100% |
| 142 | Cartilage intermediate layer protein 2 OS=Bos taurus OX=9913 GN=CILP2 PE=4 SV=1 | E1BKL4 | 126 kDa | 8.7 | Extracellular Matrix | 4 | 0.039% | 0 | 0.000% |
| 143 | Cluster of Chitinase OS=Bos taurus OX=9913 GN=CHIA PE=3 SV=2 (F1MH27) | F1MH27 [2] | 52 kDa | 5.36 | Enzyme/Protease | 4 | 0.039% | 21 | 0.192% |
| 144 | Complement factor properdin OS=Bos taurus OX=9913 GN=CFP PE=4 SV=1 | A0A3Q1MHU8 (+1) | 50 kDa | 8.33 | Complement | 4 | 0.039% | 4 | 0.036% |
| 145 | Decorin OS=Bos taurus OX=9913 GN=DCN PE=1 SV=2 | P21793 | 40 kDa | 8.74 | Extracellular Matrix | 4 | 0.039% | 0 | 0.000% |
| 146 | Ig-like domain-containing protein OS=Bos taurus OX=9913 PE=4 SV=1 | A0A3Q1LU84 (+1) | 15 kDa | 8.21 | Immunoglobulin | 4 | 0.039% | 0 | 0.000% |
| 147 | Lymphocyte cytosolic protein 1 OS=Bos taurus OX=9913 GN=LCP1 PE=1 SV=1 | A0A3Q1LSN0 (+1) | 70 kDa | 5.17 | Transport/Carrier/Binding | 4 | 0.039% | 0 | 0.000% |
| 148 | Coagulation factor X OS=Bos taurus OX=9913 GN=F10 PE=4 SV=1 | A0A452DIP8 (+1) | 67 kDa | 5.96 | Coagulation | 3 | 0.029% | 6 | 0.055% |
| 149 | Collagen type VI alpha 1 chain OS=Bos taurus OX=9913 GN=COL6A1 PE=1 SV=1 | E1BI98 | 109 kDa | 5.18 | Extracellular Matrix | 3 | 0.029% | 14 | 0.128% |
| 150 | Collagen type VI alpha 3 chain OS=Bos taurus OX=9913 GN=COL6A3 PE=1 SV=2 | E1BB91 | 340 kDa | 5.87 | Extracellular Matrix | 3 | 0.029% | 0 | 0.000% |
| 151 | Complement C1q subcomponent subunit A OS=Bos taurus OX=9913 GN=C1QA PE=2 SV=1 | Q5E9E3 | 26 kDa | 9.12 | Complement | 3 | 0.029% | 0 | 0.000% |
| 152 | Cytokine like 1 OS=Bos taurus OX=9913 GN=CYTL1 PE=4 SV=2 | E1BN89 | 16 kDa | 6.73 | Immunity/Immunity Related | 3 | 0.029% | 0 | 0.000% |
| 153 | Fibromodulin OS=Bos taurus OX=9913 GN=FMOD PE=1 SV=2 | P13605 | 43 kDa | 5.57 | Extracellular Matrix | 3 | 0.029% | 0 | 0.000% |
| 154 | Immunoglobulin J chain OS=Bos taurus OX=9913 GN=JCHAIN PE=1 SV=1 | Q3SYR8 | 18 kDa | 4.92 | Immunoglobulin | 3 | 0.029% | 0 | 0.000% |
| 155 | Insulin-like growth factor binding protein acid labile subunit OS=Bos taurus OX=9913 GN=IGFALS PE=2 SV=1 | Q09TE3 | 66 kDa | 6.73 | Structural/Cytoskeleton | 3 | 0.029% | 7 | 0.064% |
| 156 | Insulin-like growth factor-binding protein 6 OS=Bos taurus OX=9913 GN=IGFBP6 PE=4 SV=1 | A0A3Q1LTP5 (+2) | 25 kDa | 8.5 | Protease Inhibitor | 3 | 0.029% | 0 | 0.000% |
| 157 | L-lactate dehydrogenase OS=Bos taurus OX=9913 GN=LDHB PE=1 SV=1 | A0A3Q1M5R4 (+1) | 37 kDa | 5.86 | Enzyme/Protease | 3 | 0.029% | 0 | 0.000% |
| 158 | Peptidase D OS=Bos taurus OX=9913 GN=PEPD PE=3 SV=1 | A0A3Q1NGC5 (+1) | 55 kDa | 5.53 | Enzyme/Protease | 3 | 0.029% | 0 | 0.000% |
| 159 | Peptidoglycan recognition protein 2 OS=Bos taurus OX=9913 GN=PGLYRP2 PE=3 SV=3 | E1BH94 | 63 kDa | 6.46 | Immunity/Immunity Related | 3 | 0.029% | 0 | 0.000% |
| 160 | Uncharacterized protein OS=Bos taurus OX=9913 GN=C8G PE=4 SV=1 | A0A3Q1NH68 | 35 kDa | 10.52 | N/A | 3 | 0.029% | 3 | 0.027% |
| 161 | Histone H4 OS=Bos taurus OX=9913 GN=LOC112445649 PE=3 SV=1 | A0A3Q1LM20 (+17) | 15 kDa | 11.12 | sport/Carrier/Binding/Carrier/Bi | 2 | 0.019% | 0 | 0.000% |
| 162 | Phosphoglycerate mutase 1 OS=Bos taurus OX=9913 GN=PGAM1 PE=2 SV=3 | Q3SZ62 | 29 kDa | 6.67 | Enzyme/Protease | 2 | 0.019% | 0 | 0.000% |
| 163 | Protein S100-A9 OS=Bos taurus OX=9913 GN=S100A9 PE=3 SV=3 | F1MHS5 | 17 kDa | 6.29 | Immunity/Immunity Related | 2 | 0.019% | 0 | 0.000% |
| 164 | Transforming growth factor-beta-induced protein ig-h3 OS=Bos taurus OX=9913 GN=TGFBI PE=1 SV=2 | P55906 | 74 kDa | 6.69 | Cell-Cell Adhesion | 2 | 0.019% | 3 | 0.027% |
| 165 | Vitamin K-dependent protein C OS=Bos taurus OX=9913 GN=PROC PE=4 SV=2 | A0A140T851 (+2) | 51 kDa | 6.06 | Coagulation | 2 | 0.019% | 4 | 0.036% |
| 166 | Adenosylhomocysteinase OS=Bos taurus OX=9913 GN=AHCY PE=1 SV=1 | A0A3Q1LW27 (+1) | 47 kDa | 5.89 | Enzyme/Protease | 0 | 0.000% | 6 | 0.055% |
| 167 | Alpha-fetoprotein OS=Bos taurus OX=9913 GN=AFP PE=2 SV=1 | Q3SZ57 | 69 kDa | 5.92 | Transport/Carrier/Binding | 0 | 0.000% | 268 | 2.444% |
| 168 | Apolipoprotein B OS=Bos taurus OX=9913 GN=APOB PE=1 SV=3 | E1BNR0 | 516 kDa | 6.24 | Apolipoprotein | 0 | 0.000% | 275 | 2.508% |
| 169 | Apolipoprotein M OS=Bos taurus OX=9913 GN=APOM PE=3 SV=1 | A0A3Q1MB70 (+1) | 25 kDa | 6.89 | Apolipoprotein | 0 | 0.000% | 8 | 0.073% |

| | | | | | | | | | |
|---|---|---|---|---|---|---|---|---|---|
| 170 | Argininosuccinate synthase OS=Bos taurus OX=9913 GN=ASS1 PE=1 SV=1 | A0A3Q1MM52 (+1) | 48 kDa | 6.81 | Enzyme/Protease | 0 | 0.000% | 2 | 0.018% |
| 171 | Cadherin-5 OS=Bos taurus OX=9913 GN=CDH5 PE=4 SV=2 | G3MZL1 (+1) | 87 kDa | 5.3 | Cell-Cell Adhesion | 0 | 0.000% | 16 | 0.146% |
| 172 | Carboxypeptidase N catalytic chain OS=Bos taurus OX=9913 GN=CPN1 PE=3 SV=1 | G5E5V0 (+1) | 53 kDa | 8.76 | Enzyme/Protease | 0 | 0.000% | 4 | 0.036% |
| 173 | Cation-independent mannose-6-phosphate receptor OS=Bos taurus OX=9913 GN=IGF2R PE=4 SV=3 | F1MIE6 | 275 kDa | 5.62 | Transport/Carrier/Binding | 0 | 0.000% | 10 | 0.091% |
| 174 | CD109 molecule OS=Bos taurus OX=9913 GN=CD109 PE=3 SV=1 | A0A3Q1LI93 (+6) | 163 kDa | 5.5 | Protease Inhibitor | 0 | 0.000% | 2 | 0.018% |
| 175 | Cluster of 15-oxoprostaglandin 13-reductase OS=Bos taurus OX=9913 GN=PTGR1 PE=3 SV=1 (F1N2W0) | F1N2W0 [3] | 36 kDa | 7.65 | Enzyme/Protease | 0 | 0.000% | 2 | 0.018% |
| 176 | Cluster of 2-phospho-D-glycerate hydro-lyase OS=Bos taurus OX=9913 GN=ENO1 PE=3 SV=1 (A0A3Q1MXQ0) | A0A3Q1MXQ0 [5] | 54 kDa | 9.12 | Enzyme/Protease | 0 | 0.000% | 4 | 0.036% |
| 177 | Cluster of Thrombospondin-1 OS=Bos taurus OX=9913 GN=THBS1 PE=3 SV=1 (A0A3Q1MQV3) | A0A3Q1MQV3 [4] | 130 kDa | 4.69 | Cell-Cell Adhesion | 0 | 0.000% | 28 | 0.255% |
| 178 | Cluster of Thrombospondin-4 OS=Bos taurus OX=9913 GN=THBS4 PE=3 SV=1 (A0A452DJ62) | A0A452DJ62 [2] | 106 kDa | 4.41 | Cell-Cell Adhesion | 0 | 0.000% | 11 | 0.100% |
| 179 | Cluster of von Willebrand factor OS=Bos taurus OX=9913 GN=VWF PE=4 SV=1 (A0A3Q1LLU1) | A0A3Q1LLU1 [3] | 308 kDa | 5.35 | Coagulation | 0 | 0.000% | 5 | 0.046% |
| 180 | Coagulation factor V OS=Bos taurus OX=9913 GN=F5 PE=3 SV=3 | F1N0I3 | 240 kDa | 5.74 | Coagulation | 0 | 0.000% | 13 | 0.119% |
| 181 | Coagulation factor XI OS=Bos taurus OX=9913 GN=F11 PE=2 SV=1 | Q5NTB3 | 70 kDa | 8.32 | Coagulation | 0 | 0.000% | 2 | 0.018% |
| 182 | Coagulation factor XII OS=Bos taurus OX=9913 GN=F12 PE=4 SV=2 | F1MTT3 (+1) | 66 kDa | 7.61 | Coagulation | 0 | 0.000% | 3 | 0.027% |
| 183 | Coagulation factor XIII B chain OS=Bos taurus OX=9913 GN=F13B PE=4 SV=1 | A0A3Q1MWQ1 (+1) | 68 kDa | 6 | Coagulation | 0 | 0.000% | 6 | 0.055% |
| 184 | Collagen alpha-1(I) chain OS=Bos taurus OX=9913 GN=COL1A1 PE=1 SV=3 | P02453 | 139 kDa | 9.28 | Extracellular Matrix | 0 | 0.000% | 8 | 0.073% |
| 185 | Collagen alpha-1(II) chain OS=Bos taurus OX=9913 GN=COL2A1 PE=1 SV=4 | P02459 | 142 kDa | 8.38 | Extracellular Matrix | 0 | 0.000% | 2 | 0.018% |
| 186 | Collagen alpha-1(III) chain OS=Bos taurus OX=9913 GN=COL3A1 PE=2 SV=1 | Q08E14 | 138 kDa | 5.96 | Extracellular Matrix | 0 | 0.000% | 5 | 0.046% |
| 187 | Collagen alpha-2(I) chain OS=Bos taurus OX=9913 GN=COL1A2 PE=1 SV=1 | A0A3Q1LZN8 (+2) | 120 kDa | 9.3 | Extracellular Matrix | 0 | 0.000% | 12 | 0.109% |
| 188 | Collectin-43 OS=Bos taurus OX=9913 GN=CL43 PE=1 SV=2 | P42916 | 34 kDa | 5.04 | Immunity/Immunity Related | 0 | 0.000% | 7 | 0.064% |
| 189 | CPN2 protein OS=Bos taurus OX=9913 GN=CPN2 PE=2 SV=1 | A6QP30 | 60 kDa | 5.66 | Enzyme/Protease | 0 | 0.000% | 14 | 0.128% |
| 190 | CTRB1 protein OS=Bos taurus OX=9913 GN=CTRB1 PE=2 SV=1 | A5PJB8 | 28 kDa | 5.17 | Enzyme/Protease | 0 | 0.000% | 10 | 0.091% |
| 191 | C-X-C motif chemokine OS=Bos taurus OX=9913 GN=PPBP PE=1 SV=2 | F1MD83 | 32 kDa | 9.41 | Immunity/Immunity Related | 0 | 0.000% | 12 | 0.109% |
| 192 | Cysteine-rich secretory protein 2 OS=Bos taurus OX=9913 GN=CRISP3 PE=2 SV=1 | Q3ZCL0 | 27 kDa | 8.55 | Immunity/Immunity Related | 0 | 0.000% | 3 | 0.027% |
| 193 | Elongation factor 1-alpha 1 OS=Bos taurus OX=9913 GN=EEF1A1 PE=1 SV=1 | P68103 | 50 kDa | 9.1 | Other | 0 | 0.000% | 7 | 0.064% |
| 194 | Ferritin OS=Bos taurus OX=9913 PE=3 SV=1 | A0A3Q1MM35 (+1) | 20 kDa | 5.87 | Storage | 0 | 0.000% | 3 | 0.027% |
| 195 | Galectin-3-binding protein OS=Bos taurus OX=9913 GN=LGALS3BP PE=1 SV=1 | A7E3W2 | 62 kDa | 5.25 | Immunity/Immunity Related | 0 | 0.000% | 14 | 0.128% |
| 196 | Gluconolactonase OS=Bos taurus OX=9913 GN=RGN PE=3 SV=1 | A0A3Q1MLX2 (+1) | 34 kDa | 5.54 | Enzyme/Protease | 0 | 0.000% | 10 | 0.091% |
| 197 | Glutathione S-transferase A2 OS=Bos taurus OX=9913 GN=GSTA2 PE=2 SV=4 | O18879 | 26 kDa | 8.67 | Enzyme/Protease | 0 | 0.000% | 3 | 0.027% |
| 198 | Glutathione S-transferase OS=Bos taurus OX=9913 GN=GSTA1 PE=3 SV=1 | A0A3Q1MEC8 | 32 kDa | 7.01 | Enzyme/Protease | 0 | 0.000% | 3 | 0.027% |
| 199 | Glyceraldehyde-3-phosphate dehydrogenase OS=Bos taurus OX=9913 GN=GAPDH PE=1 SV=4 | P10096 | 36 kDa | 8.52 | Enzyme/Protease | 0 | 0.000% | 5 | 0.046% |
| 200 | Hemoglobin fetal subunit beta OS=Bos taurus OX=9913 PE=1 SV=1 | P02081 | 16 kDa | 6.51 | Transport/Carrier/Binding | 0 | 0.000% | 83 | 0.757% |
| 201 | Hepatocyte growth factor-like protein OS=Bos taurus OX=9913 GN=MST1 PE=3 SV=2 | E1BDW7 (+1) | 80 kDa | 8.42 | Enzyme/Protease | 0 | 0.000% | 6 | 0.055% |
| 202 | HGF activator OS=Bos taurus OX=9913 GN=HGFAC PE=4 SV=3 | E1BCW0 | 70 kDa | 7.66 | Coagulation | 0 | 0.000% | 16 | 0.146% |
| 203 | Hyaluronan-binding protein 2 OS=Bos taurus OX=9913 GN=HABP2 PE=2 SV=1 | Q5E9Z2 | 62 kDa | 5.94 | Extracellular Matrix | 0 | 0.000% | 4 | 0.036% |

| | | | | | | | | | |
|---|---|---|---|---|---|---|---|---|---|
| 204 | Ig-like domain-containing protein OS=Bos taurus OX=9913 PE=4 SV=2 | G3MY71 (+1) | 23 kDa | 9.42 | Immunoglobulin | 0 | 0.000% | 3 | 0.027% |
| 205 | Insulin-like growth factor II OS=Bos taurus OX=9913 GN=IGF2 PE=3 SV=1 | A0A3Q1LM00 (+3) | 21 kDa | 9.27 | Other | 0 | 0.000% | 3 | 0.027% |
| 206 | Insulin-like growth factor-binding protein 2 OS=Bos taurus OX=9913 GN=IGFBP2 PE=1 SV=2 | P13384 | 34 kDa | 6.93 | Protease Inhibitor | 0 | 0.000% | 8 | 0.073% |
| 207 | Inter-alpha-trypsin inhibitor heavy chain H3 OS=Bos taurus OX=9913 GN=ITIH3 PE=3 SV=1 | A0A3Q1LQ21 (+1) | 99 kDa | 5.91 | Transport/Carrier/Binding | 0 | 0.000% | 93 | 0.848% |
| 208 | Mannose-binding protein C OS=Bos taurus OX=9913 GN=MBL PE=2 SV=1 | O02659 | 26 kDa | 5.11 | Complement | 0 | 0.000% | 5 | 0.046% |
| 209 | Mimecan OS=Bos taurus OX=9913 GN=OGN PE=2 SV=1 | A5D9E8 (+1) | 34 kDa | 5.21 | Extracellular Matrix | 0 | 0.000% | 2 | 0.018% |
| 210 | Myoglobin OS=Bos taurus OX=9913 GN=MB PE=1 SV=3 | P02192 | 17 kDa | 6.97 | Transport/Carrier/Binding | 0 | 0.000% | 5 | 0.046% |
| 211 | Osteomodulin OS=Bos taurus OX=9913 GN=OMD PE=4 SV=1 | G3X6Y4 (+1) | 49 kDa | 5.21 | Extracellular Matrix | 0 | 0.000% | 2 | 0.018% |
| 212 | Osteonectin OS=Bos taurus OX=9913 GN=SPARC PE=3 SV=1 | A0A3Q1N541 | 44 kDa | 5.05 | Extracellular Matrix | 0 | 0.000% | 2 | 0.018% |
| 213 | Peptidyl-prolyl cis-trans isomerase A OS=Bos taurus OX=9913 GN=PPIA PE=1 SV=2 | P62935 | 18 kDa | 8.34 | Enzyme/Protease | 0 | 0.000% | 3 | 0.027% |
| 214 | Periostin OS=Bos taurus OX=9913 GN=POSTN PE=1 SV=1 | A0A3Q1NNK1 | 90 kDa | 7.91 | Extracellular Matrix | 0 | 0.000% | 25 | 0.228% |
| 215 | Phosphatidylethanolamine-binding protein 1 OS=Bos taurus OX=9913 GN=PEBP1 PE=1 SV=2 | P13696 | 21 kDa | 7.39 | Protease Inhibitor | 0 | 0.000% | 3 | 0.027% |
| 216 | Phosphatidylinositol-glycan-specific phospholipase D OS=Bos taurus OX=9913 GN=GPLD1 PE=1 SV=1 | P80109 | 93 kDa | 6.06 | Enzyme/Protease | 0 | 0.000% | 4 | 0.036% |
| 217 | Plasma kallikrein OS=Bos taurus OX=9913 GN=KLKB1 PE=2 SV=1 | Q2KJ63 | 71 kDa | 8.62 | Complement | 0 | 0.000% | 18 | 0.164% |
| 218 | Profilin-1 OS=Bos taurus OX=9913 GN=PFN1 PE=1 SV=2 | P02584 | 15 kDa | 8.5 | Structural/Cytoskeleton | 0 | 0.000% | 3 | 0.027% |
| 219 | Sulfhydryl oxidase OS=Bos taurus OX=9913 GN=QSOX1 PE=3 SV=3 | F1MM32 | 67 kDa | 9.29 | Enzyme/Protease | 0 | 0.000% | 4 | 0.036% |
| 220 | Thyroglobulin OS=Bos taurus OX=9913 GN=TG PE=3 SV=1 | A0A3Q1LKN2 (+5) | 275 kDa | 5.42 | Storage | 0 | 0.000% | 2 | 0.018% |
| 221 | Triosephosphate isomerase OS=Bos taurus OX=9913 GN=TPI1 PE=3 SV=1 | A0A452DIX3 (+1) | 31 kDa | 5.89 | Enzyme/Protease | 0 | 0.000% | 2 | 0.018% |
| 222 | Tubulin alpha chain OS=Bos taurus OX=9913 GN=TUBA4A PE=3 SV=1 | A0A3Q1M1Z2 (+2) | 46 kDa | 5.91 | Structural/Cytoskeleton | 0 | 0.000% | 2 | 0.018% |
| 223 | Tubulin beta-5 chain OS=Bos taurus OX=9913 GN=TUBB5 PE=2 SV=1 | Q2KJD0 | 50 kDa | 4.78 | Structural/Cytoskeleton | 0 | 0.000% | 4 | 0.036% |
| 224 | VASN protein OS=Bos taurus OX=9913 GN=VASN PE=2 SV=1 | A4IFA5 | 72 kDa | 8.63 | Cell-Cell Adhesion | 0 | 0.000% | 3 | 0.027% |
